# Susceptibility and Regulation of Biomolecular Condensates by Solutes

**DOI:** 10.64898/2026.01.14.699531

**Authors:** Takumi Matsuzawa, Kaarthik Varma, Teagan Bate, Charlotta Lorenz, Katherine Larina, Jonathan Bauermann, Dana Matthias, Tarik Grubić, Robert W. Style, Michel O. Steinmetz, Eric R. Dufresne

**Affiliations:** Department of Physics, Cornell University, Ithaca, NY 14853, USA; Department of Physics, Harvard University, Cambridge, MA 02138, USA; Department of Materials, ETH Zürich, 8093 Zurich, Switzerland; PSI Center for Life Sciences, Villigen PSI, Switzerland; University of Basel, Biozentrum, Basel, Switzerland; Department of Materials Science and Engineering, Cornell University, Ithaca, NY 14853, USA

## Abstract

Biomolecular condensates compartmentalize biochemistry in living cells. While *in vitro* models of condensates involve only a few components, the cytoplasm is a complex mixture with thousands of components, including many small molecules. While many macromolecular drivers of phase separation have been revealed, the contributions from small molecules have received little attention. To quantify the impact of solutes on biomolecular condensates, we introduce susceptibility, a dimensionless descriptor of condensate response to solute perturbations. We measured how three model condensates, assembled by distinct cohesive mechanisms, respond to diverse solutes including amino acids and nucleotides. Generically, solutes shift condensate phase equilibria, with susceptibilities spanning over four orders of magnitude. These values reflect underlying molecular interactions, consistent with theoretical descriptions including Flory-Huggins and polyphasic linkages. As one example of the predictive power of susceptibility, we exploit enzymatic activity to induce condensation and modulate material properties. Our work establishes susceptibility as an indicator of the sensitivity of biomolecular condensates to solutes, with implications for cell physiology and therapeutic design.

**SIGNIFICANCE STATEMENT:** Cells compartmentalize biochemistry using biomolecular condensates formed through phase separation. Although cells contain thousands of small molecules, little is known about their influence on condensation. In such complex mixtures, mapping full phase diagrams is infeasible. Alternatively, we introduce susceptibility to characterize system response around a working composition. Using three distinct model condensates and over a dozen solutes, we observe susceptibilities varying over four orders of magnitude. We provide a general thermodynamic framework that clarifies the driving forces behind these responses, rationalizing their magnitude and their dependence on location in the phase diagram. Our work provides a framework for understanding and harnessing solutes to regulate biomolecular condensation, with implications for cell physiology and therapeutic design.

## Introduction

Cells harness biomolecular condensates to spatially and temporally organize biochemical activities. These membraneless organelles can form through liquid–liquid phase separation [1, 2], which concentrates specific proteins, RNAs, and other intracellular components into dynamic, liquid-like compartments [3, 4]. Examples include nuclear bodies, P bodies, and stress granules, whose size, shape, and internal dynamics are tightly coupled to their phase behavior and biochemical activity [5–9]. These condensates assemble and dissolve in response to physicochemical cues, including compositional changes [10–12], pH [13, 14], redox balance [15–17], and ATP levels [18–20], and their dysregulation has been implicated in diverse pathologies [21, 22].

Macromolecules condense through a balance of cohesive interactions and the entropic costs of demixing. Because different condensates assemble through distinct molecular grammars [23, 24], they exhibit highly variable sensitivities to biochemical perturbations. This biochemical diversity in response has motivated efforts to identify ‘condensate modulators’ (CMods) [25], including metabolites, peptides, and small molecules that can either promote or dissolve specific condensates [26–29]. Such modulators hold promise both for therapeutics that target condensates [30–32] and for engineering condensates as chemical microreactors [33–35]. Despite the growing numbers of CMods [36–38], systematic quantitative comparisons and broad design principles are lacking.

A critical barrier to understanding the response of condensates to small molecules is the complex composition of the cytoplasm, a crowded mixture including thousands of proteins, nucleotides, metabolites, and ions. A direct mapping of the underlying phase behavior in such a system is impossible because the number of measurements required to map phase behavior grows exponentially with the number of components [39]. For a system with only eighty components, an astronomically large number of measurements, comparable to the number of atoms in the observable universe, would be required. A more tractable strategy is to characterize the system response to small chemical perturbations about a working composition. Experiments can be further simplified by focusing on the response of the dilute phase [40–43].

In this paper, we measure solute responses for multiple phase-separating proteins driven by distinct cohesive mechanisms. By measuring how the dilute phase changes when solutes are added, we quantify their impact on condensate stability and unify these effects using a single descriptor, the *susceptibility*. Susceptibility classifies solutes as promoters, inhibitors (or dissolvers), or inert with respect to phase separation, and provides a quantitative basis for comparing their effects across condensates. We show that solute responses can be classified by the underlying protein-solute interactions, and interpret their magnitudes using corresponding phenomenological models including Flory-Huggins theory and mass action. The resulting framework is readily generalizable to multicomponent mixtures, enabling design of enzymatic reactions to modulate condensate stability and suggesting roles for metabolites in the regulation of cellular condensates.

### Condensate responses to solute perturbations

We examine how added solutes shift the phase boundaries of three model condensates: the widely studied intrinsically disordered protein ‘fused in sarcoma’ (FUS) [44], the budding yeast clip-170 orthologue Bik1 [45] and bovine serum albumin (BSA) [46]. These proteins represent three distinct mechanisms of condensation. First, the low-complexity domain of FUS condenses through interactions of multivalent aromatic ‘stickers,’ connected by flexible glycine-rich spacers [23]. Second, Bik1 is a coiled-coil mediated dimer [47], which consists of several folded domains separated by flexible linker regions and condenses through a pocket–ligand interaction [48] between its N-terminal cytoskeleton-associated protein glycine-rich (CAP-Gly) domain and a C-terminal EEY/F-like (QQFF) motif [45, 49, 50]. This specific interaction promotes condensation of Bik1 [51]. Third, BSA is a fully folded protein that undergoes segregative condensation driven by depletion interactions [52] with polyethylene glycol (PEG) 4k as the crowding agent. Although not exhaustive, these mechanisms span both associative (FUS and Bik1) and segregative (BSA/PEG) phase separation [22, 53].

Measured phase diagrams of these model systems are shown in Fig. 1a–c. The systems phase-separate along dashed tie lines that connect the compositions of the two coexisting phases. For FUS and Bik1, the solutes used to scan the phase diagrams originate from components introduced during purification to maintain protein stability. In these two systems, increasing solute concentration progressively reduces the contrast between dilute and dense phases, leading to complete mixing above a threshold concentration. In the BSA/PEG system, the mixture phase-separates into PEG-rich (dilute) and BSA-rich phases. In all systems, solutes such as urea, salts, or crowding agents shift the boundary separating the one- and two-phase regions, thereby altering the dilute-phase protein concentration at coexistence.

**FIG. 1.**
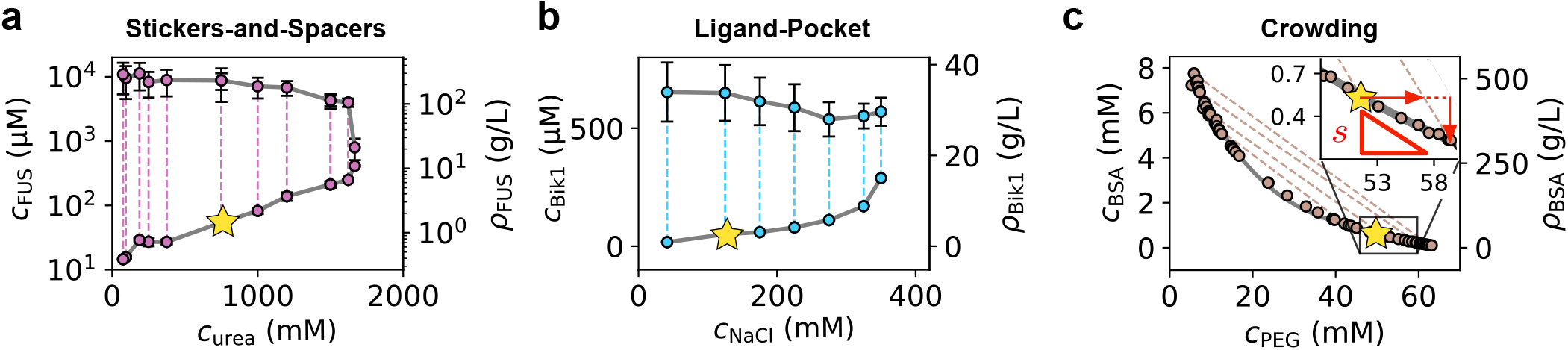
Phase diagrams of model condensates and the definition of susceptibility. **(a)** stickers-and-spacers (FUS–urea) **(b)** ligand–pocket interactions (Bik1–NaCl) **(c)** crowding (BSA–PEG). Dashed lines represent tie-lines. The addition of solutes shifts the phase boundaries, changing the dilute-phase protein concentration 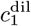. Susceptibility 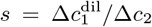 quantifies this change per unit change in the total solute concentration *c*_2_. The inset in (c) illustrates the definition of susceptibility, which is distinct from the slope of the phase boundary when solute either partitions into or is excluded from the condensate. Condensation disruptors raise 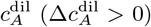, whereas promoters lower it 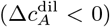. Star signs in the panels (a-c) indicate the locations in the phase space where susceptibilities are measured for Fig. 2a.

To quantify how condensates respond to solute perturbations, we define a susceptibility *s* as the change in the dilute-phase concentration of the condensing protein, 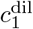, per unit change in the total concentration of a solute *c*_2_:

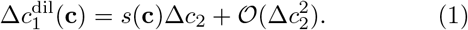

Susceptibility enables direct comparison of solute responses across condensate systems. This quantity describes how the dilute-phase concentration is altered in response to solute titration, as shown by red triangle in the inset of Fig. 1c. It is related to the slope of the phase boundary itself (see SI §IV D1 for the exact geometric relation and its connection to dominance analysis [41]). Positive (negative) values of *s* indicate that the solute promotes dissolution (condensation).

Susceptibility connects directly to the shift in the underlying free energy [43]. A small compositional perturbation Δ*c*_2_ modifies the protein’s chemical potential as

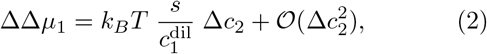

where we assume an ideal dilute phase where activities reduce to concentrations. We adopt the standard double-Δ notation to indicate the change in chemical potential between perturbed and reference states (see SI §IV A for discussion).

There is an abundance of metabolites in the cytosol [55, 56]. As a first step toward exploration of the biochemical diversity of the cellular milieu, we measured susceptibilities for diverse amino acids and nucleotides. To isolate intrinsic solute effects from pH variations, all samples are prepared in buffers at a fixed pH value. For each condition, we measure dilute-phase protein concentrations using UV–Vis spectroscopy or Bradford assays, from which susceptibilities are obtained by linear fits to at least five data points. These experiments are performed at the reference composition, labeled by stars in Fig. 1a-c. These compositions lie near the dilute-arm of the phase boundary, where condensates occupy only a small fraction of system volume. Sample preparation is automated using a liquid-handling robot to ensure reproducibility (see SI §I C). The screen includes 13 solutes (6 amino acids, 5 nucleotides, 1 crowding agent (PEG4k), and 1 dissolving agent (1,6-hexanediol)), comprising over 800 prepared samples in total.

The resulting data, shown in Fig. 2a, shows that metabolites *generically* modulate phase equilibria of biomolecular condensates. Nearly all tested solutes induce measurable shifts in the dilute phase; however, the concentrations required to produce comparable effects vary substantially across solutes and condensates. In BSA condensates, most metabolites promoted condensation, with arginine as a notable exception. In contrast, the same metabolites generally inhibit condensation in the case of FUS. This variability in susceptibility reflects the distinct molecular forces that stabilize each condensate. To dissect these interaction-specific effects, we next focus on amino acids, a major class of cellular metabolites as their chemical diversity mirrors the interaction types that drive condensation.

**FIG. 2.**
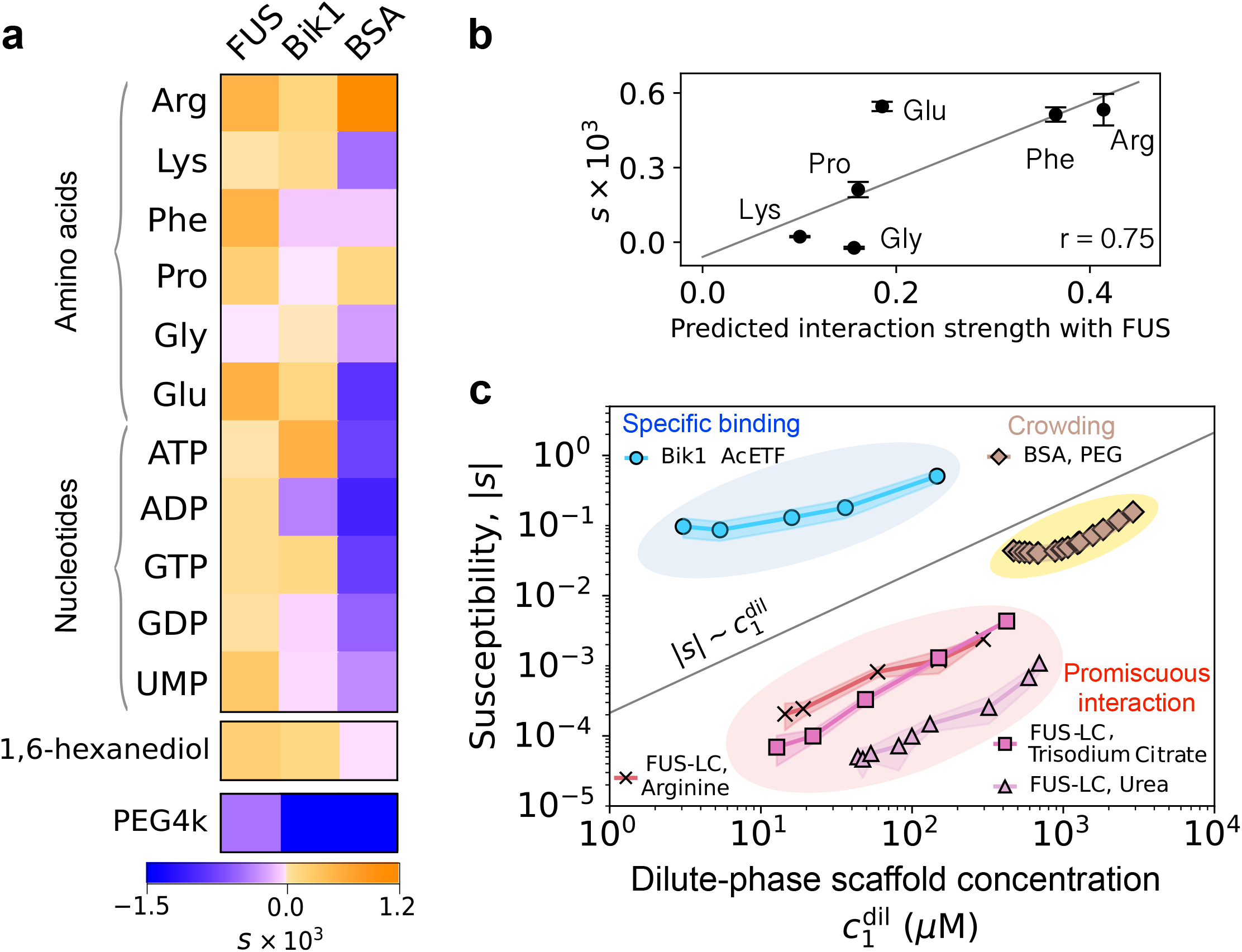
Metabolites shift phase equilibria of biomolecular condensates. **(a)** Susceptibility of model condensates (FUS, Bik1, and BSA) across metabolites reveal diverse solute responses, reflecting their underlying molecular interactions. 1,6-hexanediol, a commonly used condensation disruptor, is shown as a reference. **(b)** Susceptibility of FUS-LC to amino acids positively correlates with the predicted interaction strength computed from the Mpipi model for intrinsically disordered proteins [54]. **(c)** Susceptibility magnitude increases with scaffold concentration in the dilute phase 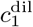 across protein-solute interaction types. Promiscuous interactions cluster at |*s*| ≤ 10^−3^, crowding at |*s*| ~ 10^−1^, and specific binding at |*s*| ~ 1. Across all interaction types, |*s*| increases with dilute-phase protein concentration 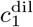, consistent with both Flory–Huggins and polyphasic linkage models.

### Metabolites regulate condensate stability

Amino acids are abundant cellular metabolites, present at tens to hundreds of millimolar concentrations, yet their role in condensate regulation is only beginning to be understood [23, 24, 26, 27, 57]. The susceptibility values of amino acids differ widely across our three model condensates (Fig. 2a). Even identical solutes exhibit opposite effects to condensates. For example, glutamic acid dissolves FUS condensates but stabilizes BSA condensates. These results demonstrate that amino-acid regulation of condensation is not universal but instead reflects the molecular grammar by which condensates are stabilized. *π*-rich or sp^2^-containing amino acids, such as phenylalanine and arginine, modulate the stability of FUS condensates, while their influence on Bik1 and BSA is minimal. Arginine is the only amino acid that has the same sign of effect on all three systems (Fig. 2a).

The importance of *π*-mediated interactions in condensation is well established [58–60]. Mpipi [54] is a coarse-grained model that simulates condensation of intrinsically disordered proteins from amino-acid sequence using an interaction matrix parameterized by *π*–*π* contact frequencies, asymmetric cation–*π* interactions, and electrostatics. We hypothesize that free amino acids interact with intrinsically disordered proteins in a manner analogous to residue–residue interactions within the sequence. To test this, we compute the average interaction strength of each free amino acid along the FUS sequence using the Mpipi interaction matrix. As shown in Fig. 2b, susceptibility values correlate with this estimated interaction strength (Pearson’s coefficient *r* = 0.75). The susceptibilities capture both the strong efficacy of aromatic amino acids and the asymmetry between arginine and lysine. Consistent with Mpipi predictions that arginine contributes more strongly to condensation than lysine despite identical charge [61], our measurements show lysine to be largely inert while arginine acts as a relatively strong dissolver.

Nucleotides are another major class of metabolites implicated in condensation processes. Cellular energy state is tightly coupled to condensate assembly, with stress granules forming under diverse stress conditions including ATP depletion [19, 62–64]. Extending these observations, our susceptibility measurements show that nucleotides generate a selective yet distinct pattern of condensate regulation. ATP, ADP, GTP, GDP, and UMP each modulate phase equilibria with magnitudes comparable to amino acids, but their effects differ markedly across systems (Fig. 2a). All these nucleotides shift FUS toward dissolution. In contrast, these same metabolites promote phase separation in BSA condensates, while Bik1 exhibits a mixed response. As with amino acids, nucleotide regulation is therefore nonuniform across condensates and depends on the molecular grammar specific to each scaffold.

Across all pairs of the solutes and condensates, the magnitude of susceptibility remains remarkably constrained within |*s*|≈10^−5^ − 10^−3^ despite large differences in molecular structure and interaction mechanisms (Fig. 2a). Even 1,6-hexanediol, often assumed to be a ‘universal’ condensate dissolver [15, 65], falls within this range, and several metabolites exhibit susceptibilities that are comparable or even higher. On the other hand, PEG, a widely used polymer to induce condensation [66], produces some of the strongest susceptibilities observed, with |*s*|≈ 10^−2^ − 10^−1^. This raises a central question: what determines the scale of susceptibility?

### Promiscuous interactions generically modulate phase equilibria

The solutes that we have considered so far interact weakly and non-specifically with scaffold proteins. For such *promiscuous* interactions, a mean-field model, such as Flory–Huggins theory, provides a natural starting point. In this framework, interactions between species are encoded by *χ*_*ij*_ parameters that quantify the relative energetic penalty or gain of mixing. Starting from a phase-separating binary mixture, with dilute-phase composition 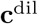, we apply the Flory-Huggins theory to determine how the dilute-phase concentration of condensing protein, 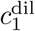, is altered when a small amount of an additional species (solute 2) is introduced while enforcing phase equilibrium. Assuming an ideal dilute phase, we find

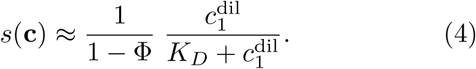

where *Φ* is the volume fraction of the dense phase, *v*_*i*_ *≡ N*_*i*_*v*_*i*,monomer_ denotes the molecular volume of component *i, v*_0_ is the molecular volume of the solvent, and *N*_*A*_ is Avogadro’s number. Here, *χ*^Δ^ = *χ*_12_ − *χ*_13_ − *χ*_23_, and *h* accounts for solute partitioning and is proportional to (*k*_2_ − 1), where 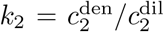 is the solute partition coefficient. This term vanishes when the solute does not partition. See SI §IV E for derivation.

The key insight from Eq. 3 is that susceptibility can be decomposed into three independent factors. First, the prefactor *v*_2_/*v*_0_ reflects the increased frequency of solute–protein encounters for larger solutes. Second, the factor 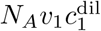 corresponds to the protein volume fraction in the dilute phase, implying that susceptibility increases as more protein is available in the dilute phase, consistent with the measurements (Fig. 2c). Third, the term (1 + *χ*^Δ^ + *h*) encodes the relative favorability of solute–protein interactions and determines the overall sign of *s*. The parameter *χ*^Δ^ may be positive or negative; positive values correspond to segregative solute–protein interactions, where solutes preferentially remain solvated rather than associating with proteins [40]. The term *h* can likewise take either sign, with partitioning (*k*_2_ > 1) favoring condensation. Even in the absence of energetic gain or solute partitioning (*χ*^Δ^ = 0 and *h* = 0), the addition of solute lowers chemical potential, favoring condensation. Thus, *χ*^Δ^ must be sufficiently negative (*χ*^Δ^ < −1 *− h*) to promote dissolution.

With typical parameters 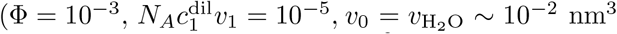, solute volumes *v*_2_ = 10^−1^− 10^0^ nm^3^), the perturbative Flory-Huggins theory typically predicts *s* between 10^−4^− 10^−3^ when solute–protein interaction strength is ~ *k*_*B*_*T*. This estimate naturally explains the order of susceptibilities observed in our experiments using amino acids, nucleotides, and 1,6-hexanediol. Susceptibility to crowding agents such as PEG4k falls between 10^−2^ and 10^−1^, approximately 100 times higher than the values observed for metabolites. This enhancement primarily reflects the much larger volume of PEG4k [67], *v*_PEG4k_ = *N*_2_*v*_monomer_ ≈ 20 nm^3^, in comparison with the volume of the metabolites (e.g. *v*_ATP_ = 1.4 nm^3^ [68]). Indeed, varying the PEG chain length shows that susceptibility systematically increases with solute volume across the model condensates, provided that the chemical interaction parameter *χ* remains unchanged (see SI § III B). However, we find a volume dependence 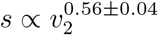, which deviates from the Flory–Huggins prediction *s ∝ v*_2_. Polyuridine RNA of different lengths likewise shows a monotonic increase in susceptibility to FUS condensates with increasing number of monomers, with 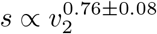.

Our extension of Flory–Huggins theory considers how added solutes modify phase equilibria through changes in bulk free energy. However, it does not explicitly resolve how local reorganization of biomolecules, solvent, and solutes stabilize biomolecules. Such local distribution of solutes and solvent molecules around biomolecules is known to be crucial in protein denaturation [69] and specific ion effects [70] in homogeneous protein solutions. These effects are more naturally described through the concept of preferential binding in Kirkwood–Buff (KB) theory [71, 72], which relates intermolecular spatial correlations to thermodynamic response functions. Interpreting susceptibility in light of KB theory, we find that solutes favor dissolution when they are enriched around proteins in the dilute phase, and favor condensation when they have repulsive interactions (see SI §IV C). Next, we consider a distinct regime in which solutes bind strongly to biomolecules.

### Specific interactions amplify susceptibility

Solutes can disrupt condensation far more efficiently when they bind tightly to specific sites responsible for condensation. Bik1 is a clear candidate for this effect as its phase separation is driven by a specific pocket–ligand interaction. Each Bik1 dimer carries two CAP-Gly domains and two EEY/F-like motifs, which form one-to-one bonds to assemble a liquid network [51]. Disrupting this interaction directly yields large susceptibility; The N-acetylated peptides ETF and EETF (AcETF and AcEETF), which mimic the C-terminal EEY/F motif of the Bik1 binding partner Bim1 (PMID: 29576319), make the susceptibility three orders of magnitude higher than promiscuous solutes such as amino acids and nucleotides (Fig. 3a). The corresponding susceptibilities are *s*_ETF_ = 0.50 *±*0.03 and *s*_EETF_ = 1.2 *±* 0.1. In terms of stoichiometry, it translates to roughly one or two peptides per Bik1 molecule, consistent with a one-to-one binding stoichiometry. This drastic dissolution of condensates is visible using a gradient chamber, in which Bik1 droplets are initially confined between two hydrogels loaded with different peptide concentrations. As the peptide (AcEETF) diffuses into the chamber, Bik1 droplets gradually swell and ultimately dissolve (Fig. 3b and SI Movie 1).

**FIG. 3.**
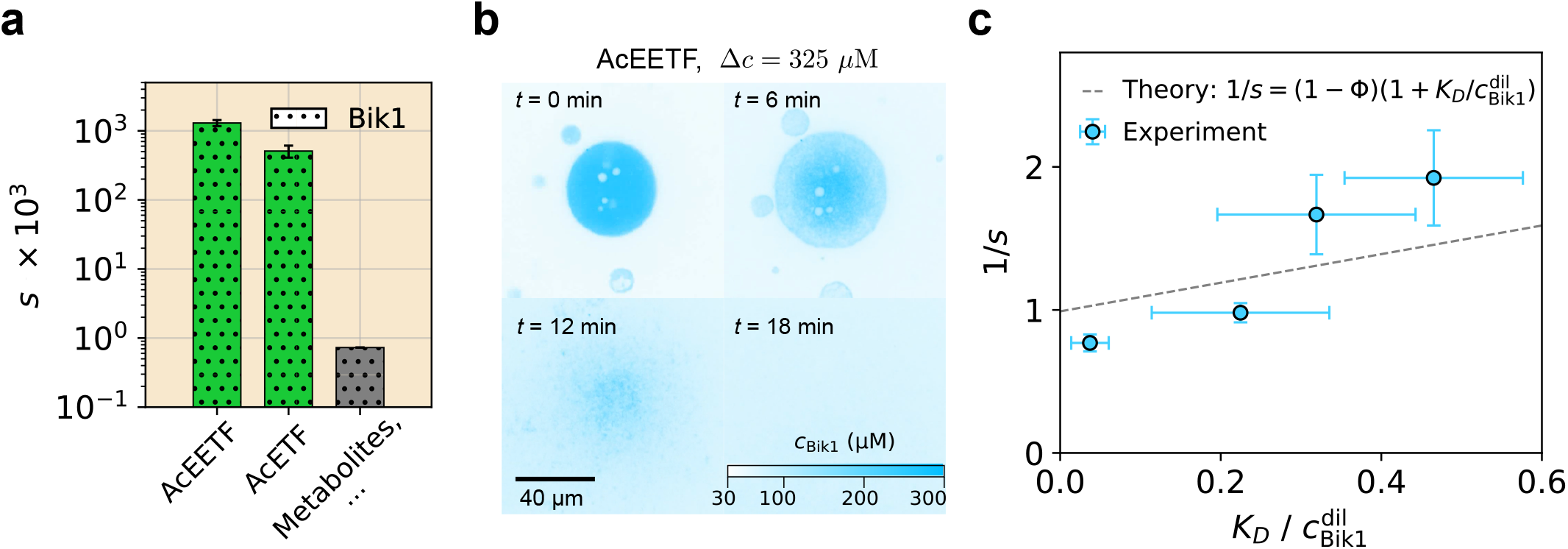
Susceptibility is amplified for solutes that strongly bind to protein pockets involved in condensate formation. **(a)** Susceptibilities *s* of Bik1 to peptides (N-acetylated EETF and N-acetylated ETF) that specifically bind to the active sites, responsible for condensation, are three orders of magnitude larger than those of typical metabolites. **(b)** Time-lapse micrographs show dissolution of Bik1 condensates at the center of the chamber as the peptide (AcEETF) concentration gradually increases. **(c)** Susceptibility to strongly binding solutes is inversely proportional to 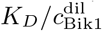, the ratio of binding affinity *K*_*D*_ and dilute-phase Bik1 concentration 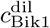, consistent with the polyphasic linkage theory prediction 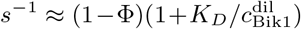. The x-error bars denote 90% confidence intervals obtained via a parametric bootstrap of the independent binding model, and the y-error bars indicate the standard deviation of the 1/*s* values (*n* = 3).

The binding of AcEETF to Bik1 violates assumptions of the Flory-Huggins theory where the solutes interact with proteins in a promiscuous manner. To achieve a susceptibility as high as observed for AcEETF to Bik1, the Flory-Huggins theory would require the unphysical *χ*^Δ^ value as high as 1000. When solutes bind tightly to scaffold proteins, the polyphasic linkage theory is a natural candidate to replace the Flory-Huggins theory [48, 73, 74]. Here the macromolecule *M* and ligand *X* are in equilibrium *M* +*X* ⇌ *MX* in both dilute and dense phases, with dissociation constants *K*_*D*_ = (*c*_*M*_ *c*_*X*_)/*c*_*MX*_ defined separately in each phase. Unlike Flory-Huggins, which encodes interactions *via χ*, this framework focuses on the solute-protein interaction *via* the binding free energy as Δ*G*_binding_ = *k*_*B*_*T* ln *K*_*D*_. In the limit where free ligands are scarce 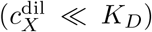, we find that the susceptibility takes the functional form (see SI §IV G for derivation):

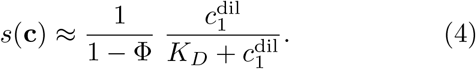

Equation 4 links susceptibility to binding affinity; the stronger a ligand binds to the site driving condensation, the more disruptive it is. In the strong binding limit *K*_*D*_→ 0, *s* approaches 1/(1− Φ) which is of order one in most experimental conditions. To test this prediction, we measured the binding affinity of AcETF and AcEETF using isothermal titration calorimetry (ITC) [75]. We performed ITC under conditions where Bik1 remained homogeneously mixed, and repeated the measurements at elevated salt concentration to obtain additional data points (see SI §III C). Fig. 3c shows that *s*^−1^ scales linearly with 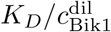, consistent with Eq. 4 with no fitting parameters.

### Susceptibility increases with dilute-phase scaffold concentration

Having examined how both promiscuous and strongly binding solutes modulate condensation, we now turn to how susceptibility varies across phase space. Figure 2c summarizes susceptibility against the dilute-phase scaffold concentration 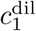 for multiple condensate–solute pairs. For promiscuous solutes, including metabolites and macromolecular crowders, susceptibility shows a linear dependence on 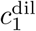. Notably, this scaling holds despite large differences in molecular size and interaction chemistry, indicating that the trend is governed primarily by the dilute-phase scaffold concentration rather than solute-specific details. A similar trend is observed for the strongly binding solutes. In the Bik1 system, the susceptibility to the peptide AcETF exhibits an approximately linear dependence on 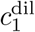, particularly at large 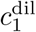.

This empirical scaling is consistent with both Flory–Huggins and polyphasic linkage models (Eqs. 3 and 4), which predict 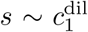, provided that 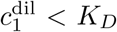 in the polyphasic linkage model. This shared scaling behavior across distinct interaction models indicates a common physical origin. As 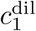 increases, the dilute and dense phases become progressively more similar in composition, with proteins more weakly held together within the condensates. As a result, solutes more efficiently perturb phase equilibrium, and condensates become increasingly susceptible to compositional changes. This linear dependence on *c*^dil^ is a robust and general feature of susceptibility, supported by thermodynamic arguments (see SI §IV B).

### Extension to multi-solute systems

Up to this point, our analysis has focused on susceptibilities to individual solutes. However, in physiological settings, multiple solutes can change concentration simultaneously. Such multi-solute variations commonly arise in processes such as energy metabolism, signal transduction, osmotic regulation, post-translational modification [10, 20, 76–83].

The susceptibility framework is readily applied to such multi-solute changes. When solutes are non-interacting, we expect that susceptibilities add linearly, and the dilute-phase response is given by

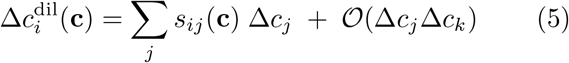

where 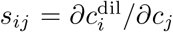 is the generalized susceptibility of component *i* to solute *j*. Our measurements on binary metabolite mixtures support this hypothesis (See SI §III D). Deviations from linearity can be interpreted as the result of interactions between solutes, which may act cooperatively or antagonistically to modulate 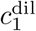.

Applying this framework to chemical reactions, the modulation of condensation can be predicted from a linear sum of susceptibilities, weighted by the reaction stoichiometry. By assigning each substrate and product a measured susceptibility, we define a reaction susceptibility,

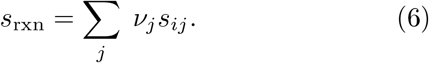

where *v*_*j*_ encodes the stoichiometry of the reaction (*v*_*j*_ < 0 for substrates and *v*_*j*_ > 0 for products). If *s*_rxn_ < 0, the reaction drives condensation; if *s*_rxn_ > 0, it favors dissolution.

### Modulation of Condensate Phase Behavior

With this principle in hand, we can pick an enzymatic reaction with the desired sign of *s*_rxn_. Equation 6 indicates that reactions exhibiting a larger difference in susceptibilities between substrates and products are more effective at modulating the phase behaviors of condensates. Our screen shows that ATP suppresses condensation of Bik1 while ADP promotes it, making ATPase a natural pick to promote condensation (*s*_rxn_ < 0; Fig. 4a). ATP hydrolysis proceeds through two sequential steps, ATP → ADP → AMP, and whether the reaction terminates after the first or proceeds to the second depends on the enzyme. Here, we use apyrase, a ubiquitous ATPase present in eukaryotes and some prokaryotes, which catalyzes both steps sequentially to yield AMP (Fig. 4b). Under our conditions, ATP hydrolysis is the dominant reaction, as it proceeds approximately seven times more efficiently than ADP hydrolysis (see SI §III E2). Guided by this prediction, we form Bik1 condensates in the presence of apyrase, ATP, and calcium ions (cofactor of the enzyme). Apyrase partitions into Bik1 condensates (Fig. 4c) and retains enzymatic activity in the experimental buffer (5 mM CaCl_2_, 20 mM Tris, 150–375 mM NaCl, pH 6.8), as confirmed by ITC (see SI §III E1 and 2).

**FIG. 4.**
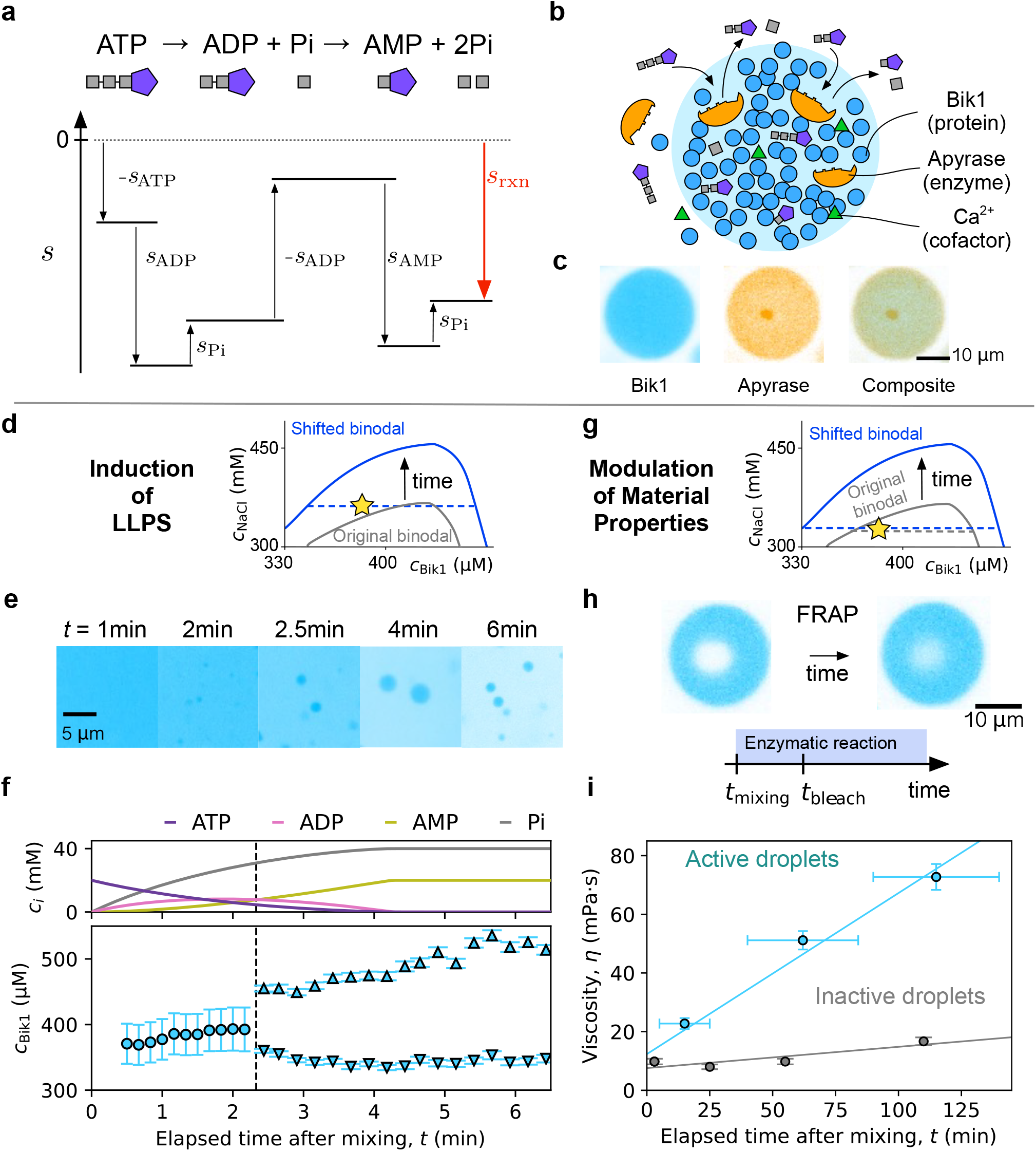
Susceptibility to enzymatic substrates and products enables control of condensate stability and material properties. **(a) Susceptibility measurements show that ATP promotes Bik1 dissolution (***s* > 0**), while ADP and AMP promote phase separation (***s <* 0**). The reaction susceptibility**, *s*_rxn_, **defined as the stoichiometry-weighted sum of substrate and product susceptibilities, predicts whether the reaction promotes or inhibits Schematic of apyrase-catalyzed ATP hydrolysis inside Bik1 condensates. (c) Apyrase partitions into Bik1 condensates with a partition coefficient of** 1.60 *±* 0.20, visualized by composite fluorescence images. **(d–f)** Enzyme-induced condensation. (d) Schematic: ATP hydrolysis drives an upward binodal shift. Star marks experimental composition. (370 *µ*MBik1, 375 mM NaCl, 20 mM ATP, 5 mM CaCl_2_, 20 mM Tris, pH 6.8) (e) Time-lapse micrographs show that droplets emerge from a homogeneous phase, indicating condensation onset as hydrolysis progresses. (f) Top: Simulated concentrations of ATP, ADP, AMP, and inorganic phosphate from Michaelis–Menten kinetics using measured parameters. Bottom: Bik1 concentration extracted from micrographs shows a bifurcation at *t* = 2.2 min, marking the condensation onset. The vertical error bars represent STD. (*n* ≈ 70 droplets) **(g–i)** Enzymatic activity alters condensate rheology. (g) Schematic: Star marks the initial composition, which lies within the two-phase region at the start of the experiment. (325 mM NaCl; all other conditions are the same as in (d)) (h) FRAP experiments are performed while enzymatic reaction proceeds. (i) Viscosity increases over time for apyrase-loaded droplets, in contrast to inactive droplets containing apyrase but lacking cofactor. Horizontal error bars indicate a half the time for each FRAP experiment; vertical bars represent STD (*n* = 7 droplets).

Starting from homogeneous solutions containing Bik1, ATP, and calcium chloride, we observe that condensates nucleate and grow as ATP is consumed (Fig. 4d–e and SI Movie 2). Following the onset of condensation, Bik1 continues to be recruited into droplets, increasing the contrast between the dense and dilute phases (Fig. 4e). This enrichment is quantified in Fig. 4f, which shows the widening gap between dilute- and dense-phase concentrations as the reaction proceeds. Substrate and product concentrations are based on the Michaelis–Menten kinetics with competing substrates for ATP and ADP, obtained from the ITC experiments [84] (see SI §III E2).

By contrast, when droplets are present, ATP hydrolysis further promotes condensation, driving the system deeper into the two-phase region of the phase diagram (Fig. 4g). We measure Bik1 diffusivity within condensates using fluorescence recovery after photobleaching (FRAP) as the enzymatic reaction progresses (Fig. 4h). Because the reaction continues during measurement, the diffusivity of Bik1 decreases over time. Nevertheless, FRAP recovery curves are described by a single exponential, indicating that diffusion equilibrates faster than the enzymatic reaction timescale (see SI §III E3). Converting the apparent diffusivity to viscosity using the Einstein–Stokes relation shows that active droplets thicken linearly over time, whereas inactive controls (lacking the cofactor) maintain essentially constant viscosities. These results demonstrate that enzymatic activity not only promotes condensation but also progressively alters the mechanical properties of condensates as the reaction proceeds.

### Concluding remarks

We have shown that biomolecular condensation is generically sensitive to solute perturbations, and that this sensitivity can be quantitatively captured by the susceptibility across condensate–solute pairs. Susceptibility directly quantifies how phase equilibria shift under compositional perturbations and is tied to the underlying thermodynamics and molecular interactions. Across a wide range of systems, experimentally measured susceptibilities span a vast range from 10^−5^ to 10^0^, consistent with standard descriptions of condensation, including Flory–Huggins [85, 86], and polyphasic linkage theories [74, 87]. Across the diversity of molecular interactions, susceptibility increases with the dilute-phase protein concentration 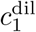. While susceptibility is most easily quantified by modulating individual solutes, these data can be applied to multicomponent perturbations, including shifts in phase equilibria driven by chemical reactions. Evaluating the reaction susceptibility enables the identification of new biochemical routes for regulating condensation and material properties.

In living cells, thousands of metabolites and other small molecules coexist in the cytosol, making it infeasible to understand condensate phase behavior using full phase diagrams. Susceptibility overcomes this limitation by shifting the focus from global phase behavior to local, experimentally accessible responses. Care must be taken when interpreting susceptibility values, as they depend strongly on the location in the phase diagram. Comparisons of susceptibilities across species are meaningful only when the condensates are in the same state. For example, FUS condensates are 10 times more susceptible to arginine as the 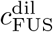 increases by a factor of 10, as shown in Fig. 2c. Because *in vivo* condensates are known to have low partition coefficients of scaffold proteins [88], even weakly interacting solutes can exhibit large susceptibilities to shift their phase equilbria.

The broad range of susceptibility values has important implications for cellular physiology and therapeutics. For solutes with small susceptibility, large concentration changes are required to substantially modulate condensate phase behavior. In physiological settings, such large variations occur for metabolites that are abundant in the cytosol, including ATP (1 − 10 mM) [56, 89, 90], amino acids (on the order of 10 − 100 mM) [26, 91], and other osmolytes such as trehalose (on the order of 100 – 1000 mM) [92]. Large concentration shifts can compensate for small susceptibility values. By contrast, solutes with large susceptibility are natural candidates for condensate-targeted drugs, as small doses can influence condensation while minimizing off-target effects. These include small molecules that strongly bind to target proteins, as well as molecules that engage in promiscuous interactions whose effects are amplified by large solute volumes and partitioning. For the latter, examining the role of partitioning in drug discovery is a natural next step [93].

Our findings support previous studies that have shown that condensates are not passive phase-separated droplets but dynamically regulated biochemical assemblies whose onset and material properties can be tuned in a spatiotemporal manner by biological processes such as metabolism [20, 77, 94], signaling pathways [76, 79–83], and post-translational modifications [77, 95–97]. What distinguishes our work is the role of metabolites as generic modulators of condensate stability. Previous studies have primarily focused on altering condensate behavior through direct modification of scaffold proteins [23, 83, 95, 98, 99]. In contrast, our results show that even weakly interacting solutes can exert substantial effects when present at sufficient concentrations, within the reach of physiological settings. From this perspective, we show that enzymatic reactions can regulate condensates indirectly by producing and consuming small molecules, even without directly modifying proteins. This work paves the way to program condensates with desired phase and material properties by identifying the relevant solutes, biochemical reactions, and compositional conditions.

## ACKNOWLEDGMENTS

The authors thank Nathaniel Hess, Francesco Stellacci, Madhurima Choudhury, and members of the Dufresne lab for helpful discussions and comments on the manuscript. We also thank Allison Mae Dew for assistance with the ITC measurements. This work was partially supported by the Swiss National Science Foundation through the NCCR Bio-Inspired Materials (Grant No. 205603) and Sinergia program (Grant No. 189940). TM acknowledges support from Schmidt Science Fellows, in partnership with Rhodes Trust, and CL acknowledges funding from the Deutsche Forschungsgemeinschaft (Proj. Nr. 523842861, LO 3088/1-1). We thank the Cornell Center for Materials Research at Cornell University for access to its shared experimental facilities (US NSF grant DMR 1719875) and the BRC Imaging Core Facility (RRID SCR 021741) at the Cornell Institute of Biotechnology for enabling FRAP experiments.

## AUTHOR CONTRIBUTIONS

T.M. and E.R.D. designed the project, analyzed, and interpreted the results, and wrote the manuscript with inputs from K.V., T.B., J.B., R.W.S., T.G., and M.O.S. T.M., K.V., and T.B. purified proteins with assistance from T.G., C.L., and M.O.S. T.M. and K.V. automated liquid handling for sample preparation. T.M., K.V., T.B., and D.M. performed experiments to determine phase diagrams. T.M. and K.V. measured susceptibility values. T.M. performed and analyzed calorimetry and confocal microscopy experiments. K.L. developed the protocol for the gradient chamber experiments. T.M., J.B., R.W.S. and E.R.D. developed the theoretical models. E.R.D. directed the research.

## DATA AVAILABILITY

The data underlying the plots in this article are available from the corresponding author upon reasonable request. See Supplementary Information §III A for the exact values of susceptibility shown in Fig. 2a.

## CODE AVAILABILITY

The code used to operate the liquid-handling robot is available via the author’s GitHub repository (https://github.com/Laboratory-of-Soft-and-Living-Materials/2026_Matsuzawa_Susceptibility).

