## Supplementary Information for "Susceptibility and Regulation of Biomolecular Condensates by Solutes"

### Susceptibility and Regulation of Biomolecular Condensates by Solutes Supplementary Information

Jonathan Bauermann

*Department of Physics, Harvard University, Cambridge, MA 02138, USA*

Dana Matthias and Robert W. Style

*Department of Materials, ETH Zürich, 8093 Zurich, Switzerland*

Tarik Grubić

*PSI Center for Life Sciences, Villigen PSI, Switzerland*

Michel O. Steinmetz

*PSI Center for Life Sciences, Villigen PSI, Switzerland and*

*University of Basel, Biozentrum, Basel, Switzerland*

Eric R. Dufresne

*Department of Physics, Cornell University, Ithaca, NY 14853, USA and*

*Department of Materials Science and Engineering, Cornell University, Ithaca, NY 14853, USA*

(Dated: May 18, 2026)

**CONTENTS**


I. Materials and methods 4
A. Proteins and solutes 4
1. FUS 4
2. Bik1 4
3. BSA 4
4. Apyrase 4
5. Solutes 5
B. Confocal microscopy and PEGMA-acrylamide glass coating 5
C. Sample preparation 6
1. Automated sample preparation using a liquid handler 6
2. pH variation in susceptibility measurements 6
D. Gradient Chamber Experiment (Bik1 × Peptides) 8
E. Isothermal Titration Calorimetry (ITC) 9
F. Fluorescence recovery after photobleaching (FRAP) 9

II. Protein, condensation Mechanism, and chemical properties 10

III. Experiments 11
A. Susceptibility values in Figure 2A 11
B. Dependence of susceptibility on solute volume 13
C. ITC experiment: AcEETF and AcETF bind to Bik1 14
D. Additivity of susceptibility 15
E. Modulation of Bik1 droplets using apyrase 17
1. Quantification of Partitioning Coefficient of Apyrase to Bik1 droplets 17
2. Kinetics of Apyrase-mediated ATP hydrolysis 19
3. FRAP experiments show that activity of apyrase thickens Bik1 condensates. 21

IV. Theory 22
A. Derivation of $\Delta\Delta\mu_i = k_B T \sum_j \frac{s_{ij}}{c_i^{\mathrm{dil}}} \Delta c_j$ 22
B. Dependence of susceptibility on $c_1^{\mathrm{dil}}$ originates from entropy 23
C. Relationship between susceptibility and Kirkwood–Buff theory 26
1. Results at fixed volume $V$ and temperature $T$ 26
2. Results at fixed pressure $P$ and temperature $T$ 28
D. Geometrical interpretation of susceptibility $s$ 31
1. Relationship among susceptibility $s$, binodal gradient $r$, and tie-line slope $t$ in two-component systems 31
2. Extension to $N$-component systems that separate into two phases 34
E. Susceptibility to promiscuous (non-specific) interactions 35
1. Estimation of $s$ using the Flory Huggins theory without solute partitioning 35
2. Estimation of $s$ using the Flory Huggins theory with solute partitioning 36
3. Summary 39
F. Susceptibility to depletion interactions (crowding): BSA-PEG 40
1. Summary 41
G. Susceptibility to ligand-pocket interactions: Bik1-Peptide 43
1. Chemical equilibrium between coexisting liquid phases 43
2. Definition of susceptibility for strongly binding solutes 44
3. Derivation of Eq. 4 in the main text 45

4. Summary 48
H. Summary of susceptibility models 49

V. Captions for Supplementary Movies 49

A. Relation between the hessians of Helmholtz and Gibbs free energies 50

References 52

### I. MATERIALS AND METHODS

#### A. Proteins and solutes

##### 1. *FUS*

Histidine-and Gb1-tagged FUS (residues 1-270) was overexpressed in *E. coli* strain BL21 (DE3) (New England Biolabs, Ipswich, MA USA; C2527H) at 20°C overnight. A low speed spin (5 kg, 22C, 25 m) after initial lysis was used to capture exclusion bodies. The pellet was re-suspended under strong denaturing conditions (8 M urea, 50 mM HEPES, 500 mM NaCl, pH 7.5) and dounced to destroy inclusion bodies before performing a typical clarification spin (16 kg, 22C, 25 m). The protein was purified under denaturing conditions (1 M urea, 150 mM NaCl, 50 mM HEPES, pH 7.5) on a nickel affinity chromatography column, followed by enzymatic cleavage of the histidine tags overnight with histadine tagged TEV protease (in-house, pET29b-10xHis-Super TEV [1]). The protein was concomitantly dialyzed to remove imidazole (1 M urea, 150 mM NaCl, 50 mM HEPES, 1 mM DTT, pH 7.5). Next, the protein was dialyzed to stronger denaturing conditions (6 M urea, 50 mM HEPES, 150 mM NaCl, pH 7.5) to prevent phase separation during subsequent concentration. Additional nickel affinity chromatography removed the protease and cleaved products. Finally, the protein was concentrated up to 2 mM (6 M urea, 50 mM HEPES, 150 mM NaCl, pH 7.5), snap frozen and stored at -80°C. To initiate phase separation, the stock protein solution is diluted to 50 $\mu$M in buffer (50 mM HEPES and 150 mM NaCl, pH 7.5).

##### 2. *Bik1*

N-terminally tagged hexahistidine-thrombin cleavage site-*S. cerevisiae* full length Bik1 (H6-TCS-Bik1) was overexpressed in *E. coli* strain BL21-CodonPlus (DE3)-LOBSTR (Agilent, Santa Clara, CA USA; 230280) at 20°C overnight. Protein samples were purified by nickel affinity and size exclusion chromatography (500 mM NaCl, 20 mM Tris, 10% Glycerol, 1 mM DTT, pH 7.5). Bik1 protein samples were concentrated to 1-2 mM (500 mM NaCl, 20 mM Tris, 10% Glycerol, 1 mM DTT, pH 7.5). To induce phase separation, the stock protein solution is diluted in buffer (150 mM NaCl, 20 mM Tris, pH 7.5) to a final concentration of 30 - 100 $\mu$M. To image the droplets Alexa Fluor 546-NHS Ester (Succinimidyl Ester) (ThermoFisher Scientific, Waltham, MA USA; A20002) was de-activated in 1 M Tris and mixed into the droplet solution, to a final concentration of 2 $\mu$M, where it partitioned into the condensed phase with a partition coefficient $k \approx 4$.

##### 3. *BSA*

Bovine serum albumin in lyophilized powder, essentially globulin free, $\geq$99% was purchased from Millipore Sigma (A7638), and kept at 4°C.

##### 4. *Apyrase*

Apyrase from potatoes was purchased from Millipore Sigma (A7640-500UN) as lyophilized powder. Its ATPase activity listed on the label is 500 units per milligram. The protein was first suspended in 20 mM Tris, pH 6.8, and desalted using a desalting column to remove impurities. It was then concentrated to 1 mg/mL (22 $\mu$M for a molecular weight of 49 kDa), aliquoted in 20 mM Tris, pH 6.8, and stored at $-20$°C.

To measure its partition coefficient in Bik1 droplets, apyrase was dissolved in 20 mM bicarbonate buffer, pH 8.3, to 1 mg/mL and labeled with Alexa 647 NHS ester (ThermoFisher Scientific, A20006) by incubating it for 2 hours at room temperature with continuous stirring. After labeling, the solution was buffer-exchanged into 20 mM Tris, pH 6.8, before use in partitioning experiments, and concentrated back to 1 mg/mL.

###### 5. *Solutes*

All solutes except for oligopeptides (N-acetylated EETF and N-acetylated ETF) were purchased from Millipore Sigma and prepared in buffer conditions specific to each model condensate (FUS: 1 M urea, 150 mM NaCl, 50 mM HEPES, 1 mM DTT, pH 7.5; Bik1: 150 mM NaCl, 20 mM Tris, pH 7.5; BSA/PEG: 200 mM NaCl, 100 mM potassium phosphate, pH 7.5). The solutes and their corresponding catalog numbers are listed in Table S1. Stock solutions were prepared near the solubility limit of each solute by dissolving powders directly into the appropriate buffer (or mixing liquids with buffer solutions), followed by pH adjustment using HCl or NaOH as needed. These stock solutions were kept at 4°C when not used.

TABLE S1. Solutes used in this study and their Millipore Sigma catalog numbers.

| **Solute** | **Catalog Number (Millipore Sigma)** |
|---|---|
| L-Arginine | A5006 |
| L-Lysine | 62840 |
| Glycine | G7126 |
| L-Proline | P0380 |
| L-Glutamic acid | G1251 |
| ATP | A1852 |
| ADP | A2754 |
| AMP | A2252 |
| GTP | G8877 |
| GDP | G7127 |
| UMP | U1752 |
| $KH_2PO_4$ | P0062 |
| $K_2HPO_4$ | P3786 |
| Sodiunm chloride | S9888 |
| Trisodium citrate dihydrate | 1064480500 |
| Urea | U5378 |
| 1,6-hexanediol | 240117 |
| PEG 400 | 06855 |
| PEG 1000 | 8.07488 |
| PEG 4000 | 95904 |
| PEG 10000 | 8.21881 |

All oligopeptides samples (EETF and ETF) were purchased from GenScript Biotech (Piscataway, NJ, USA) in an N-acetylated form with purity $\geq 95\%$ after exchanging each each buffer for FUS, Bik1, and BSA to remove trifluoroacetic acid in the samples. The pH was adjusted by adding a small amount of 1M NaOH.

##### B. Confocal microscopy and PEGMA-acrylamide glass coating

All imaging was performed on a Nikon Ti2 Eclipse microscope equipped with a Yokogawa CSU W1 spinning disk confocal unit, a Teledyne Kinetix 22 camera, and a Nikon F-LUN laser stack. A

60x water immersion objective (NA: 1.3) was used for high-resolution imaging. The microscope was configured for single-camera acquisition with triggering and a multi-pass filter. The minimum PSF was measured as 243.5 ± 11.5 nm in the xy plane by 44 nm fluorescent polystyrene microsopheres with an emission wavelength of 525 nm, using a 100X Oil objective with an NA of 1.49.

Imaging were performed on the glass slides with PEGMA-Acrylamide-glass slides coating. Slides and slips were first cleaned using Helmanex III (Sigma-Aldrich, Inc., St. Louis, MO, USA; Z805939) in boiling water, followed by sonication and thorough rinsing. Ethanol and 100 mM potassium hydroxide sonication steps etched the glass surface to expose hydroxyl groups. Silanization was performed with a solution containing trimethoxysilyl-propylmethacrylate (TOPA; Sigma-Aldrich, Inc., St. Louis, MO, USA; M6514), mixed immediately prior to use, to bind silane groups to the glass. Finally, slides and slips were coated with a 1.5% PEGMA (poly(ethylene glycol) methyl ether acrylate; Sigma-Aldrich, Inc.; 454990) and 0.5% acrylamide (Sigma-Aldrich, Inc.; A8887) solution polymerized with ammonium persulfate (APS; Sigma-Aldrich, Inc.; A3678) and tetramethylethylenediamine (TEMED; Sigma-Aldrich, Inc.; T9281) to create a crosslinked surface. Prepared slides and slips were stored in the coating solution and rinsed with DI water before use.

##### C. Sample preparation

###### *1. Automated sample preparation using a liquid handler*

To measure the susceptibility of a solute to a specific condensate, we collected 5–10 data points per condition ($n = 3$). With 20 solutes and 3 condensates, this resulted in more than 1,300 samples. To ensure reproducibility and enable efficient sample preparation with minimal human intervention, samples preparation was semi-automated using an Opentrons OT-2 liquid-handling robot in a 96-well plate. Each sample is defined by (i) the stock solutions, (ii) the target concentrations, and (iii) an ordered mixing scheme. The mixing scheme is critical as we always induce phase separation at the last step of mixing. For BSA/PEG, it is crucial to add PEG at the last step as PEG is the crowding agent. For FUS and Bik1, we mix all the ingredients except the proteins first, then add the mixture into pre-loaded protein solutions.

The mixing order is encoded as a nested list, allowing the robot to generate intermediate mixtures before assembling the final sample. For example, `[A, B, C]` instructs the robot to add A, then B, then C to the same well. A more complex example, `[A, [B, C, [D, E], F], H, [I, J]]`, creates mixtures of D+E, then B+C+(D+E)+F, then I+J, and finally combines everything in the order A $\rightarrow$ (mixture) $\rightarrow$ H $\rightarrow$ (mixture).

The OT-2 automatically assigns wells for intermediate mixtures and executes the mixing steps in the specified order, ensuring reproducible preparation of multi-component samples.

###### *2. pH variation in susceptibility measurements*

To isolate the effects of solutes from the pH changes, we prepare the concentrated solution with the target solute in the buffer. The buffers for FUS and Bik1 reflect the protein purification conditions and were selected for their stability. For the condensate in this study, we used phosphate (pH 7.0), HEPES (pH 7.5), and Tris (pH 7.5) buffer. Phosphate buffer is tripronic buffer with pKa values equal to 2.12, 7.21, and 12.44 at 20°C. The pKa of HEPES and Tris are 7.55 and 8.3, respectively. The pH of the Tris buffer is most sensitive to temperature variation among these buffers. The buffer capacity $\beta$ quantifies the amount of equivalent $H^+$ or $OH^-$ required to change the pH by one unit. It is defined as the required molarity of strong base to change unit pH, and is given by

$$\beta = \frac{dc_b}{d\text{pH}} = \ln 10 \frac{K_a[\text{H}^+]}{(K_a + [\text{H}^+])^2} \cdot c \tag{1}$$

where $c_b$ is the molarity of strong base added to the solution, $c$ is the molarity of single monoprotic buffer with the acid dissociation constant $K_a$.

In this study, the buffer capacities were 54.6 mM/pH for 100 mM phosphate (pH 7.0, BSA/PEG), 86.1 mM/pH for 150 mM HEPES (pH 7.5, FUS), and 5.44 mM/pH for 20 mM Tris (pH 7.5, Bik1) [2]. To minimize pH changes upon mixing, we adjusted the pH of stock solutions containing additional solutes prior to mixing. For example, dissolving arginine powder in buffer makes the solution slightly acidic; therefore, a small amount of NaOH was added to adjust the pH prior to mixing with the protein solution.

##### D. Gradient Chamber Experiment (Bik1 × Peptides)

As shown in the main Fig. 3, we developed a gradient-chamber assay to observe dissolution of Bik1 droplets as Bik1 strongly binds to peptides (AcEETF or AcETF). The setup consists of a fluid channel on a PEGMA-coated glass slide, hydrogel pieces in a confined chamber, and the photograph and the schematic of the assembled gradient chamber are shown in the Fig. S1a-b.

To prepare the hydrogel, we mixed agarose powder in water (3% w/v agarose), heated it in a microwave without boiling, poured the solution into a Petri dish to a thickness of approximately 5 mm, and cooled it to room temperature. From this hydrogel, we cut one thin gel strip (1 mm × 15 mm × 5 mm) and two gel blocks (5 mm × 5 mm × 5 mm). The gel blocks were then equilibrated separately in either buffer alone (20 mM Tris, 400 mM NaCl, pH 7.5) or buffer containing 325 $\mu$M AcEETF.

A phase-separated Bik1 sample (366 $\mu$M Bik1, 20 mM Tris, 400 mM NaCl, 2.5% glycerol, pH 7.5) containing fluorescent beads was loaded into a chamber assembled on a PEGMA-coated glass slide using a parafilm spacer. The two pre-equilibrated gel blocks were then placed on opposite sides of the gel strip, forming reservoirs that establish a stable diffusion gradient once the chamber is sealed to minimize evaporation.

As illustrated in Fig. S1c, AcEETF diffuses through the gel strip. The experiment reported in main Fig. 3b was performed before a steady linear gradient was established. When the local peptide concentration exceeds a threshold, Bik1 droplets expand and then gradually dissolve (Fig. S1d).

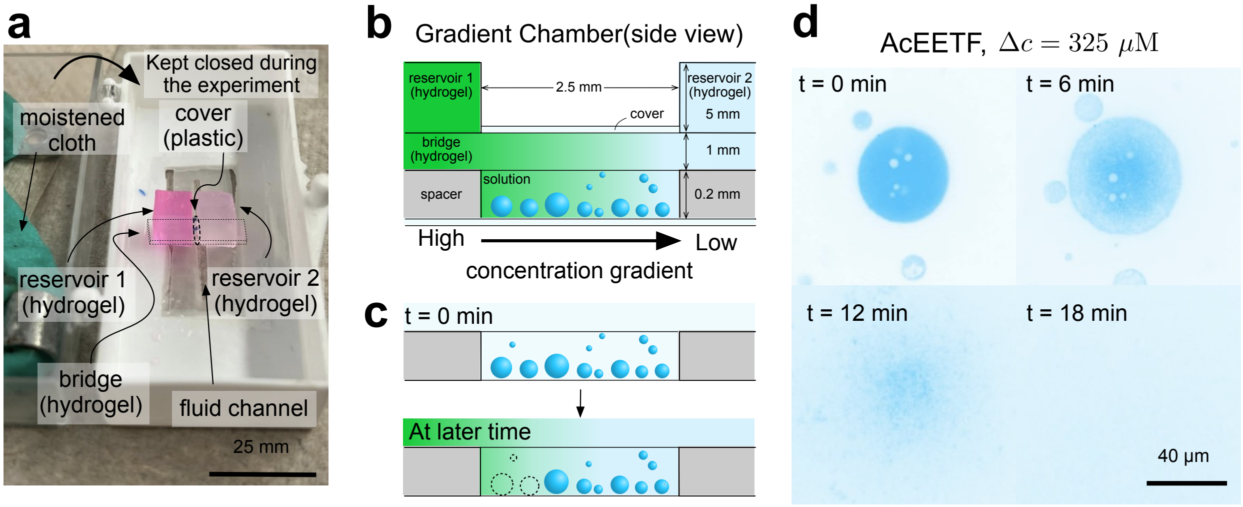


FIG. S1. **Dissolution of Bik1 condensates in a peptide (AcEETF) gradient.** **(a)** A photograph of the gradient chamber is shown. The hydrogel of the reservoir 1 contains rhodamine for the visualization purpose. In the actual experiment, it is immersed with the peptide solution.

**(b)** Schematic (side view) of the gradient chamber. Two agarose hydrogel reservoirs—one containing buffer and the other containing 325 $\mu$M AcEETF—are placed on opposite sides of a hydrogel strip to generate a linear concentration gradient across the chamber holding Bik1 droplets. **(c)** AcEETF diffuses through the hydrogel and it reaches a steady state at $t = 90$ min. Once the local peptide concentration exceeds the dissolution threshold, Bik1 droplets swell and subsequently dissolve from the high-concentration side. **(d)** Time series of Bik1 droplets as AcEETF concentration increases. Droplets swell (t = 6–12 min) and dissolve by ~18 min. Scale bar: 40 $\mu$m. (c) was replicated from the Figure 3.

##### E. Isothermal Titration Calorimetry (ITC)

Peptides (N-acetylated EETF and N-acetylated ETF) were dissolved in ITC buffer (250 or 375 mM NaCl, 20 mM Tris, 10% glycerol, pH 7.5). The pH was adjusted to 7.5 with NaOH to compensate for residual trifluoroacetic acid (TFA) from peptide synthesis. All protein and peptide samples were then buffer-exchanged into the ITC buffer using Zeba Spin Desalting Columns (7K MWCO, Thermo Fisher Scientific). ITC experiments were performed at 20 °C with a stirring speed of 75 rpm using an Affinity ITC instrument (TA Instruments). The Bik1 solution (100 $\mu$M monomer equivalents in ITC buffer) was loaded into the sample cell, and the peptide solution (900 $\mu$M) into the syringe. Bik1 solutions at both 375 mM and 250 mM NaCl remained homogeneously mixed (not phase-separated), as confirmed by UV–Vis concentration measurements performed prior to the experiments. Thirty injections of 2.5 $\mu$L were performed. Binding isotherms were fitted by nonlinear least-squares minimization using NanoAnalyze software (TA Instruments). Control experiments were performed by injecting buffer alone. Reported errors represent the 90% confidence interval, estimated using a bootstrapping method. See §III C for results.

##### F. Fluorescence recovery after photobleaching (FRAP)

FRAP experiments on Bik1 droplets with apyrase were performed using a Zeiss LSM880 laser scanning confocal microscope (Carl Zeiss Microscopy, GmbH, Jena, Germany) with a 100,× oil immersion objective (N.A. 1.4). Solutions with phase-separated droplets were prepared 1–2 minutes before imaging. The composition of the "active Bik1 droplets" was 20 mM ATP, 5 mM $CaCl_2$, 10 $\mu$g/mL apyrase, 298 $\mu$M Bik1, 292 mM NaCl, 20 mM Tris, 2% glycerol, pH 6.8. The composition of the "inactive Bik1 droplets" was the same as the active droplets except that $CaCl_2$ was omitted. For the FRAP experiments, we used Bik1 labeled with Alexa 647 such that the labeling percentage of the solution was 1%. The sample was imaged on a PEGMA-coated glass slide. A circular area (diameter = 4 $\mu$m) near the center of the droplet was photobleached to at least 50% of its initial fluorescence intensity. Recovery of fluorescence was monitored every 10 s using ZEN Microscopy Software (Carl Zeiss Microscopy, GmbH, Jena, Germany).

#### II. PROTEIN, CONDENSATION MECHANISM, AND CHEMICAL PROPERTIES

Table II summarizes the properties and driving interactions of LLPS. The net charge of FUS and Bik1 are inferred from the sequence.

TABLE S2. Properties of proteins used in this study

| Protein | Type | Driving interaction of condensation | $N$ | MW (kDa) | pI | Net charge (e) | $R_g$ (nm) |
|---|---|---|---|---|---|---|---|
| FUS | Intrinsically disordered | Multivalent ($\pi$–sp$^2$, cation–$\pi$, hydrophobic interaction) | 267 | 26.3 | 8.86 [3] | +2.15 at pH 7.5 | $3.5 \pm 0.1$ |
| Bik1 | Multidomain | Pocket–ligand interaction (CAP-Gly : EEY/F motif [4]) | 439 | 51.0 | 5.62 [5] | −14.3 at pH 7.5 | $9.2 \pm 0.5$ [6] |
| BSA | Globular | Depletion interaction induced by PEG 4k | 583 | 66.5 | 5.1-5.5 [7] | −18 at pH 7.0 [7] | 2.9 [8] |

#### III. EXPERIMENTS

##### A. Susceptibility values in Figure 2A

Table S3 reports the susceptibility values of small molecules (metabolites, 1,6-hexanediol, and PEG4k) in the model condensates.

TABLE S3. Susceptibility values $s \times 10^3$ in Figure 2A and 3A reported as mean ± fit error.

| | FUS-LC | Bik1 | BSA | Concentration Range for Linear Fit |
|---|---|---|---|---|
| Arg | $0.51 \pm 0.03$ | $0.16 \pm 0.02$ | $1.57 \pm 0.08$ | 0 – 100 mM |
| Lys | $0.02 \pm 0.00$ | $0.09 \pm 0.01$ | $-0.78 \pm 0.08$ | 0 – 100 mM |
| Phe | $0.53 \pm 0.06$ | $-0.13 \pm 0.13$ | $-0.14 \pm 0.17$ | 0 – 40 mM |
| Pro | $0.21 \pm 0.03$ | $-0.02 \pm 0.00$ | $0.13 \pm 0.01$ | 0 – 100 mM |
| Gly | $-0.02 \pm 0.00$ | $-0.00 \pm 0.00$ | $-0.41 \pm 0.02$ | 0 – 100 mM |
| Glu | $0.55 \pm 0.02$ | $0.13 \pm 0.04$ | $-1.46 \pm 0.09$ | 0 – 40 mM |
| ATP | $0.10 \pm 0.01$ | $0.54 \pm 0.16$ | $-1.27 \pm 0.17$ | 0 – 50 mM |
| ADP | $0.06 \pm 0.02$ | $-0.62 \pm 0.04$ | $-1.69 \pm 0.09$ | 0 – 50 mM |
| AMP | n/a | $-0.52 \pm 0.10$ | n/a | 0 – 40 mM |
| GTP | $0.07 \pm 0.01$ | $0.12 \pm 0.06$ | $-1.30 \pm 0.07$ | 0 – 50 mM |
| GDP | $0.04 \pm 0.01$ | $-0.07 \pm 0.06$ | $-0.90 \pm 0.05$ | 0 – 50 mM |
| UMP | $0.30 \pm 0.04$ | $-0.06 \pm 0.00$ | $-0.55 \pm 0.03$ | 0 – 50 mM |
| 1,6-hexanediol | $0.21 \pm 0.01$ | $0.12 \pm 0.02$ | $-0.05 \pm 0.02$ | 0 – 100 mM |
| PEG4k | $-0.73 \pm 0.01$ | $-4.70 \pm 0.13$ | $-12.00 \pm 1.55$ | 0 – 10 mM |
| AcEETF | n/a | $1.2 \pm 0.1$ | n/a | 0 - 100 $\mu$M |
| AcETF | n/a | $0.50 \pm 0.03$ | n/a | 0 - 1 mM |

Measurements were performed under the following conditions:

1. FUS-LC condensates: 51.8 $\mu$M FUS-LC (1.34 mg/mL), 150 mM urea, 150 mM NaCl, 50 mM HEPES, pH 7.5.

2. Bik1 condensates: 39.8 $\mu$M Bik1 (2.07 mg/mL), 150 mM NaCl, 20 mM Tris, 34 mM glycerol, pH 7.5.

3. BSA condensates: 700 $\mu$M BSA (46.7 mg/mL), 52.5 mM PEG4k, 200 mM KCl, 100 mM phosphate buffer, pH 7.0.

Fig. 1a–c illustrates the locations in phase space at which the experiments were performed. Figure S2 shows representative confocal microscopy images of the condensates obtained without added solutes. In all measurements, the condensates exhibited liquid-like behavior such as droplet coalescence.

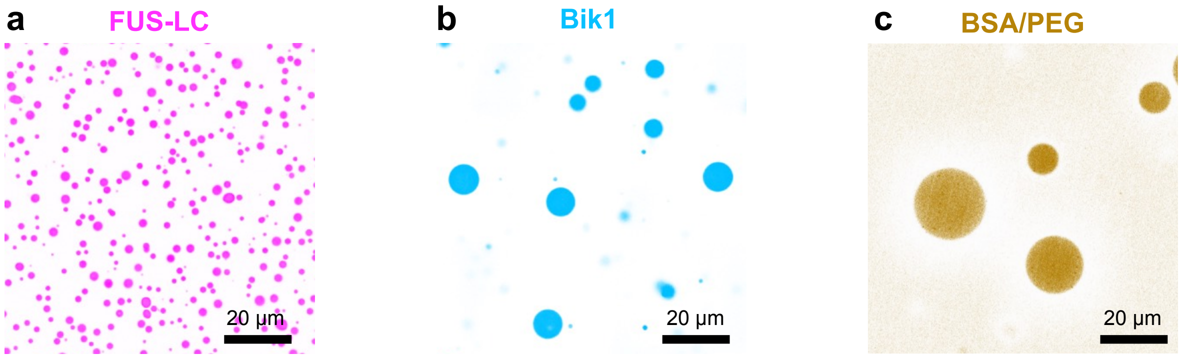


FIG. S2. **Representative confocal microscopy images of model condensates in the absence of added solutes.**
**(a)** FUS-LC, **(b)** FUS-LC, **(c)** BSA/PEG condensates. Scale bars, 20 $\mu$m.

#### B. Dependence of susceptibility on solute volume

This section supplements the claim in the manuscript that susceptibility increases with solute molecular volume when the underlying molecular interactions are held constant ($\chi^{\Delta} = \text{const.}$). For promiscuous interactions, Flory–Huggins theory predicts that susceptibility is proportional to the solute volume $v_2$:

$$s(\mathbf{c}) \approx -\frac{1}{1-\Phi} \underbrace{\left(\frac{v_2}{v_0}\right)}_{\text{relative solute volume}} \left(N_A v_1 c_1^{\text{dil}}(\mathbf{c})\right)(1 + \chi^{\Delta} + h).$$

To test this volume dependence, we measured the susceptibility of polyethylene glycol (PEG), a commonly used crowder in biological assays [9], under identical buffer conditions while varying its molecular weight from 400 to 20,000 Da. Square symbols in Fig. S3a and b show that PEG susceptibility increases with molecular weight.

The definition of the solute volume $v_2$ may be ambiguous. In Flory–Huggins theory, $v_2$ is defined as the molecular volume, $v_2 = N_2\tilde{v}_2$ An alternative definition is the steric volume, $v_2 = \frac{4\pi}{3}R_g^3$, where $R_g$ is the radius of gyration. Because Flory–Huggins theory applies to polymers without folded structure, the use of molecular volume is appropriate for capturing protein–solute contacts. However, this assumption may not hold for general solutes, or even for long polymers, which can adopt compact configurations. For this reason, we examine the volume dependence by plotting susceptibility against the monomer volume (Fig. S3a) and the steric volume (Fig. S3b).

The susceptibilities of PEG with different molecular weights exhibits susceptibilities over 10 times larger than the metabolites in our screen. Although susceptibility increases monotonically with PEG molecular weight, the observed scaling $v_2^{0.60\pm0.01}$ deviates from the Flory–Huggins prediction $|s| \propto v_2$. This trend also persists for nucleotides: FUS condensates show greater susceptibility to longer-chain polyuridine RNA , as shown by the blue points in Fig. S3. For poly(rU) in FUS condensates, we observe a sublinear dependence on volume, $s \sim v_2^{0.72\pm0.08}$.

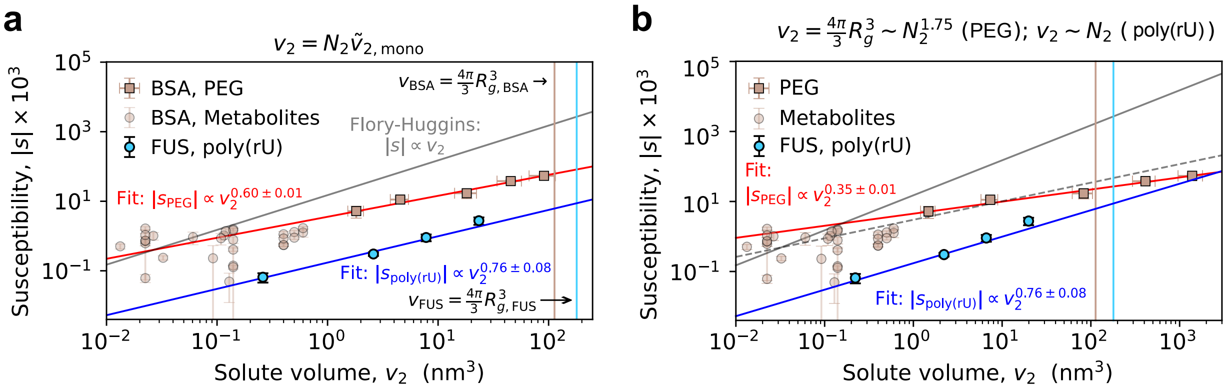


FIG. S3. **Susceptibility increases with solute volume. (a–b)** The magnitude of the susceptibility $|s|$ is shown as a function of solute molecular volume $v_2$. Brown square symbols represent polyethylene glycol (PEG) with molecular weights of 400, 1000, 2000, 4000, and 8000 g/mol, and the brown circular symbols represent the metabolites presented in Figure 2a. Blue circle symbols, from left to right, represent uridine monophosphate (UMP) and polyuridine RNAs of different lengths (rU10, rU30, and rU90).
(a) Chemical volume: $v_2 = N_2 v_{\text{mono}}; v_{\text{mono}} = 0.06\ \text{nm}^3$ for PEG [10] and 0.26 nm$^3$ for poly(rU). (b) Steric volume: $v_2 = (4\pi/3)R_g^3; R_g = 0.0215M^{0.583} \sim N_2^{1.75}$ for PEG [11] and 0.55 $N^{1/3}$ nm for RNAs [12]. Using the chemical volume, both PEG and poly(rU) with different chain lengths $N_2$ exhibit a sublinear scaling than the Flory–Huggins expectation $|s| \propto v_2$.

##### C. ITC experiment: AcEETF and AcETF bind to Bik1

This section supplements the observed binding of N-acetylated EETF (AcEETF) and N-acetylated ETF (AcETF) to Bik1. The presented data are used in Fig. 3c. Thermograms from the ITC experiments (see §I E for experimental protocols) and the corresponding binding isotherms confirm one-to-one binding of both AcEETF and AcETF to Bik1 (Fig. S4). The dissociation constant, $K_D$, obtained from fits to an independent binding model, is summarized in Table S4.

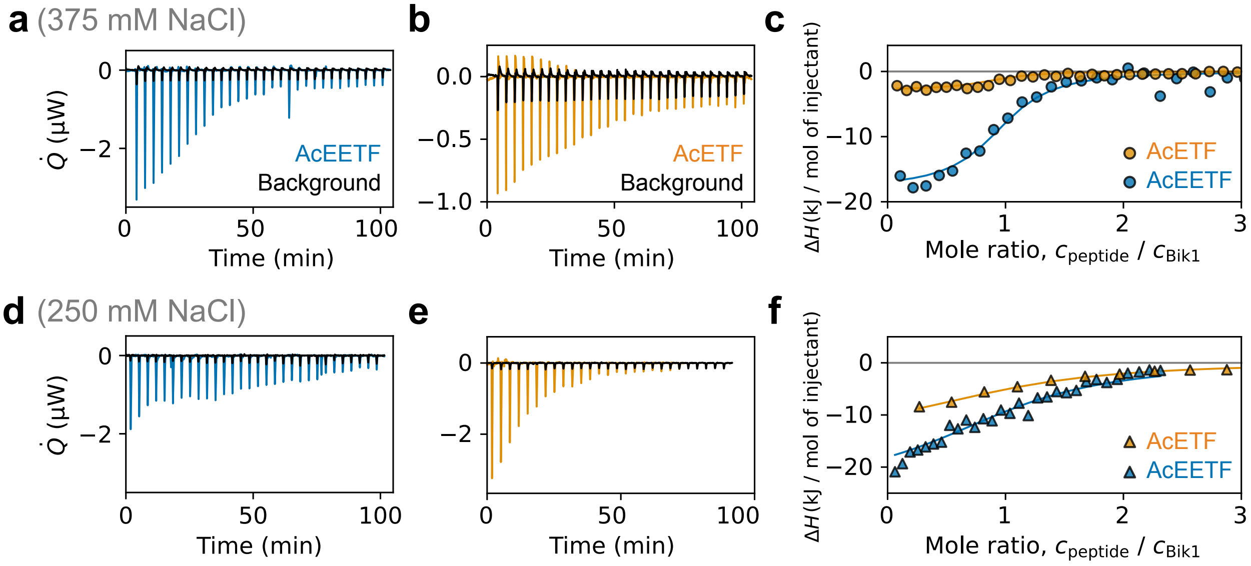


FIG. S4. **One-to-one binding between Bik1 monomers and N-acetylated peptides AcEETF and AcETF measured by ITC. (a, b)** Representative ITC thermograms for titration of Bik1 with AcEETF (blue) and AcETF (orange), respectively, under high-salt conditions. Black traces show control injections of peptide into buffer. **(c)** Corresponding binding isotherms derived from panels (a,b) and fit to an independent binding model are shown. **(d, e)** ITC thermograms for titration of Bik1 with AcEETF (blue) and AcETF (orange), respectively, under low-salt conditions. Black traces show control injections of peptide into buffer. **(f)** Binding isotherms corresponding to panels (d, e). High-salt conditions correspond to 375 mM NaCl, 20 mM Tris, pH 7.5, and low-salt conditions to 250 mM NaCl, 20 mM Tris, pH 7.5. All experiments were performed at 20 °C.

TABLE S4. **ITC measurements of Bik1 binding to AcEETF and AcETF peptides at high (375 mM) and low (250 mM) NaCl concentrations.** The dissociation constant $K_D$, stoichiometry $n$, and enthalpy change $\Delta H$ were obtained from single-site fits to the thermograms. The entropy contribution $-T\Delta S$ and free energy change $\Delta G$ were calculated from $\Delta G = RT \ln K_D = \Delta H - T\Delta S$. Uncertainties represent 90% confidence interval, estimated by a bootstrapping method.

| Name | [NaCl] (mM) | $K_D$ (μM) | $n$ | $\Delta H$ (kJ/mol) | $-T\Delta S$ (kJ/mol) | $\Delta G$ (kJ/mol) |
|---|---|---|---|---|---|---|
| Bik1 : AcEETF | 375 | $7.1 \pm 4.5$ | $0.99 \pm 0.07$ | $-15.5 \pm 1.6$ | −29 | −45 |
| Bik1 : AcETF | 375 | $43 \pm 21$ | $0.96 \pm 0.12$ | $-3.4 \pm 0.7$ | −22 | −25 |
| Bik1 : AcEETF | 250 | $32 \pm 12$ | $1.19 \pm 0.07$ | $-32.0 \pm 3.8$ | 6.6 | −26 |
| Bik1 : AcETF | 250 | $47 \pm 11$ | $0.99 \pm 0.12$ | $-13.9 \pm 2.2$ | −11 | −25 |

#### D. Additivity of susceptibility

We examined whether solute susceptibility is additive in multi-solute mixtures. For a buffer-matched mixture containing $N$ solutes that act independently, the expected susceptibility is

$$s = \sum_{j=1}^{N} x_j \, s_{ij},$$

where $x_j$ is the mole fraction of solute $j$.

To test this prediction, we titrated binary mixtures while varying the total concentration $c_A + c_B$. Figure S5a–c shows results for three solute pairs applied to BSA condensates: arginine–proline, ammonia–sodium bicarbonate, and arginine–glutamic acid.

For the arginine–proline and ammonia–bicarbonate mixtures, the measured susceptibility varies linearly with the mixing ratio, consistent with additivity. For the arginine–glutamic acid pair, nonlinear behavior emerges above a total solute concentration of 20 mM. Because these solutes carry opposite charges in addition to hydron bonding [13] and can form ion pairs, solute–solute interactions become significant at higher concentrations, producing deviations from linearity. In the dilute limit, where such interactions are weak, the additivity relation is well satisfied.

To assess linearity in binary solute mixtures, we plotted the normalized susceptibility against the mole fraction $x = c_A/(c_A + c_B)$, as shown in Fig. S6. The line $y = x$ represents perfect additivity, and deviations above or below this line indicate cooperative or antagonistic behavior, respectively. Arginine–glutamic acid (an ion-pairing combination) and ammonia–bicarbonate (an acid–base pair) show mild antagonistic deviations at higher mole fractions. These deviations are small, indicating that susceptibility remains a robust and informative metric even in the presence of modest solute–solute interactions.

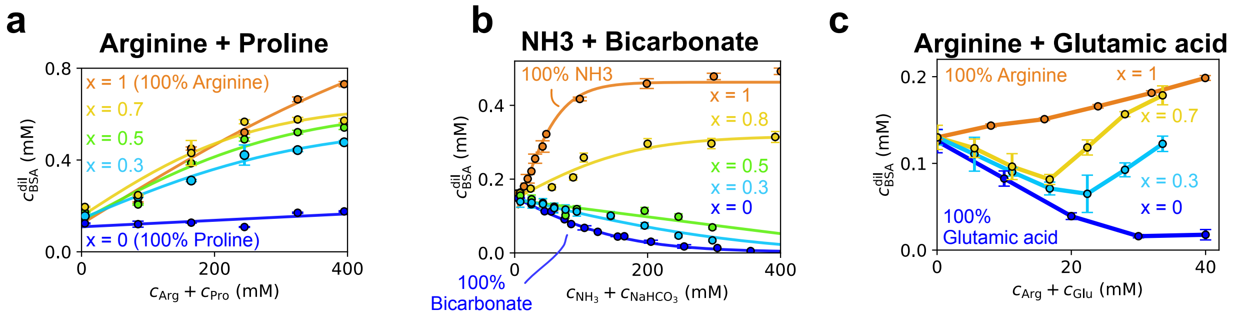


FIG. S5. **Additivity of susceptibility in binary solute mixtures.** **(a)** Arginine–proline: susceptibility varies linearly with the mixing ratio, consistent with independent solute contributions. **(b)** Ammonia–bicarbonate: susceptibility remains additive across mixing ratios. **(c)** Arginine–glutamic acid: nonlinear behavior appears above 20 mM total solute, consistent with attractive interactions between oppositely charged solutes.

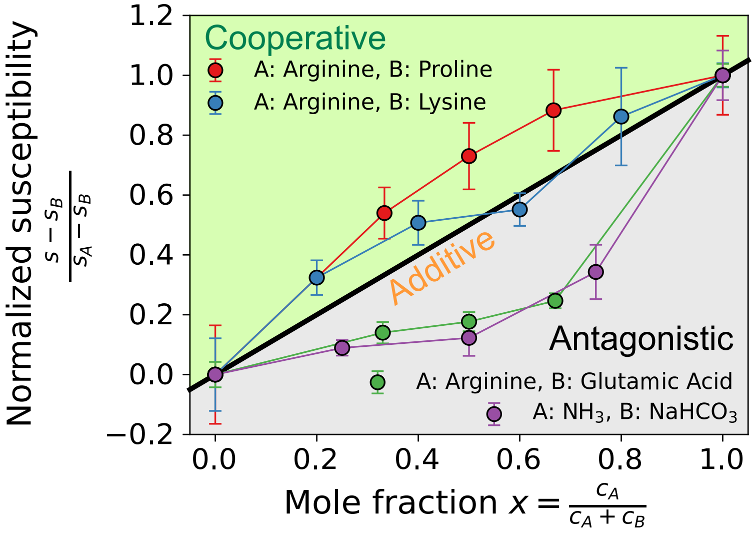


FIG. S6. **Cooperative, additive, and antagonistic effects of mixed solutes.** Normalized susceptibility curves for binary mixtures of arginine–proline, arginine–lysine, arginine–glutamic acid, and ammonia–bicarbonate. Susceptibility increases above the additive prediction in cooperative pairs, follows linear additivity for non-interacting pairs, and falls below the additive line for antagonistic pairs where solute–solute interactions suppress the response.

##### E. Modulation of Bik1 droplets using apyrase

In this section, we supplement our measurements on apyrase-loaded Bik1 droplets.

###### 1. *Quantification of Partitioning Coefficient of Apyrase to Bik1 droplets*

This section supplements the claim in main Fig. 4c that apyrase partitions into Bik1 droplets. To quantify this, we report the partition coefficient of apyrase, $k_{\mathrm{Apy}} = c_{\mathrm{Apy}}^{\mathrm{den}}/c_{\mathrm{Apy}}^{\mathrm{dil}}$, by fluorescence imaging of apyrase labeled with Alexa Fluor 488 NHS ester (succinimidyl ester). In this experiment, Bik1 was also labeled with Alexa Fluor 647 NHS ester to image droplets.

For main Fig. 4c, imaging was performed on a Nikon Ti2 Eclipse microscope equipped with a Yokogawa CSU W1 spinning-disk confocal unit, a Teledyne Kinetix 22 camera, and a Nikon F-LUN laser stack, using a 60× water-immersion objective (NA 1.3). All images were corrected for spatial inhomogeneity in illumination by dividing by a vignette intensity profile. This profile was obtained by averaging 10 independent images after Gaussian filtering with $\sigma_x = \sigma_y = 100$ px. Droplets were then identified using a custom segmentation algorithm. For each droplet, an azimuthally averaged intensity profile $I(r)$ about the centroid was computed to identify the bulk, interface, and dilute regions. The bulk and dilute regions were defined from the plateau regions of $I(r)$, and the bulk intensity was taken as the dense-phase composition to avoid interfacial effects. To further mitigate vignetting effects, we computed the partition coefficient for each droplet as $k_i = I_i^{\mathrm{den}}/I_i^{\mathrm{dil}}$, where $i$ indexes the droplets.

Figure S7 shows the measured partition coefficients of Bik1 and apyrase under conditions equivalent to those of main Fig. 4c: 20 $\mu$M Bik1, 20 mM Tris, 150 mM NaCl, pH 6.8. We find an apyrase partition coefficient of $1.7 \pm 0.1$ in the absence of cofactor and $1.6 \pm 0.2$ in its presence (5 mM). By contrast, the free dye (Alexa Fluor 488) does not partition into Bik1 droplets. We therefore conclude that apyrase weakly partitions into Bik1 droplets regardless of the presence of cofactor.

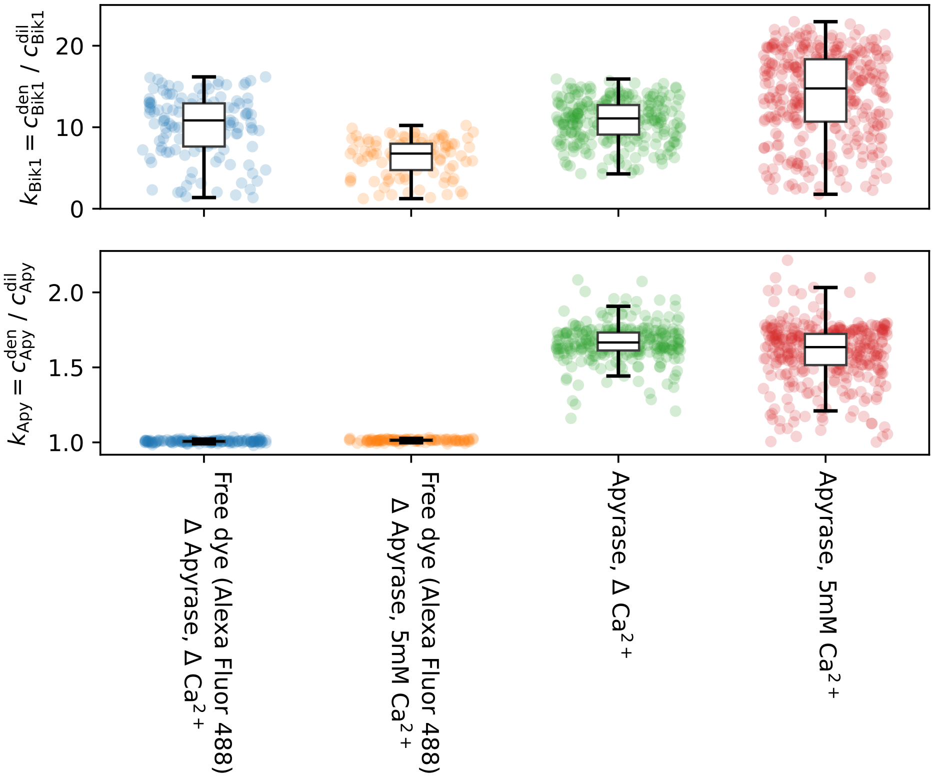


FIG. S7. **Apyrase weakly partitions into Bik1 droplets.** (Top) Partition coefficient of Bik1 under the indicated conditions. (Bottom) Partition coefficient of apyrase measured from fluorescence intensity. Free dye (Alexa Fluor 488) shows no partitioning, whereas apyrase exhibits weak enrichment in droplets, with comparable partition coefficients in the presence and absence of $Ca^{2+}$. Data points represent individual droplets ($n > 70$ for each condition); boxes indicate median and 1.5 times interquartile range (IQR).

*2. Kinetics of Apyrase-mediated ATP hydrolysis*

This section supplements Fig. 4f by describing how the substrate concentrations (ATP and ADP) were simulated. To this end, we measured the Michaelis–Menten parameters of apyrase activity.

We consider a single enzyme population with total enzyme concentration $c_E$. Step-specific kinetic parameters $(k_{\mathrm{cat},1}, K_{M,1})$ and $(k_{\mathrm{cat},2}, K_{M,2})$ are defined for the reactions ATP → ADP and ADP → AMP, respectively. Because the same active site binds both ATP and ADP, the substrates compete for the enzyme. The resulting coupled ordinary differential equations are given by

$$\frac{dc_{\mathrm{ATP}}}{dt} = -v_1, \qquad \frac{dc_{\mathrm{ADP}}}{dt} = v_1 - v_2, \qquad \frac{dc_{\mathrm{AMP}}}{dt} = v_2,$$

with the following initial conditions

$$c_{\mathrm{ATP}}(0) = c_0, \qquad c_{\mathrm{ADP}}(0) = 0, \qquad c_{\mathrm{AMP}}(0) = 0.$$

Under the rapid-equilibrium (Briggs–Haldane) assumption, the rates are given by [14]

$$v_1 = \frac{\left(\frac{k_{\mathrm{cat},1}}{K_{M,1}}\right) c_E\, c_{\mathrm{ATP}}}{1 + \frac{c_{\mathrm{ATP}}}{K_{M,1}} + \frac{c_{\mathrm{ADP}}}{K_{M,2}}}, \qquad v_2 = \frac{\left(\frac{k_{\mathrm{cat},2}}{K_{M,2}}\right) c_E\, c_{\mathrm{ADP}}}{1 + \frac{c_{\mathrm{ATP}}}{K_{M,1}} + \frac{c_{\mathrm{ADP}}}{K_{M,2}}}. \tag{2}$$

Figure S8a shows Michaelis–Menten curves for apyrase, obtained by isothermal titration calorimetry (Affinity ITC, TA Instruments, Delaware). In this assay, 20 $\mu$M ATP or ADP was titrated 30 times in 1-$\mu$L injections into a buffer-matched solution containing 20 $\mu$M apyrase (20 mM Tris, 150 mM NaCl, 5 mM $CaCl_2$, pH 6.8). The data show that ATP hydrolysis is the dominant reaction ($k_{\mathrm{cat},1} = 40 \pm 1\ \mathrm{s}^{-1}$, $K_{M,1} = 200 \pm 30\ \mu$M), whereas ADP hydrolysis is slower ($k_{\mathrm{cat},2} = 6.1 \pm 0.3\ \mathrm{s}^{-1}$, $K_{M,2} = 150 \pm 37\ \mu$M).

The experiments reported in the manuscript were performed at mildly acidic pH (6.8) because apyrase activity declines sharply with increasing pH and is nearly abolished at pH 7.5 (Fig. S8b). The extracted kinetic parameters are summarized in Table S5.

TABLE S5. **Michaelis-Menten parameters of apyrase in buffer (20 mM Tris, 150 mM NaCl, 5 mM $CaCl_2$)**

| pH | $k_{cat,1}$ ($s^{-1}$) | $K_{M,1}$ ($\mu$M) | $k_{cat,2}$ ($s^{-1}$) | $K_{M,2}$ ($\mu$M) |
|---|---|---|---|---|
| 6.8 | $40 \pm 1$ | $200 \pm 30$ | $6.1 \pm 0.3$ | $150 \pm 37$ |
| 7.2 | $17 \pm 1$ | $41 \pm 18$ | n/a | n/a |
| 7.5 | $0.70 \pm 0.10$ | $21 \pm 34$ | n/a | n/a |

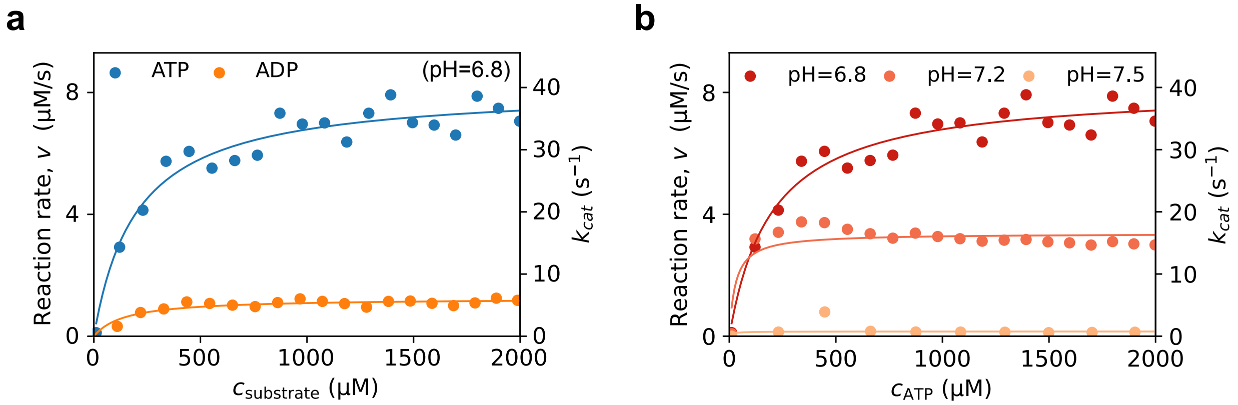


FIG. S8. **Michaelis–Menten kinetics of ATP hydrolysis by apyrase at various pH values. (a)** Reaction velocity $v$ as a function of substrate concentration (ATP or ADP) in the buffer solution (20 mM Tris, 150 mM NaCl, 5 mM $CaCl_2$pH=6.8 The curves show that ATP hydrolysis is the dominant reaction catalyzed by apyrase, whereas ADP hydrolysis is markedly slower. **(b)** Michaelis–Menten curves for ATP hydrolysis at pH 6.8, 7.2, and 7.5. In both panels, solid lines represent fits to the Michaelis–Menten model, from which the catalytic rate constant $k_{cat}$ and $K_M$ were obtained.

*3. FRAP experiments show that activity of apyrase thickens Bik1 condensates.*

This section supplements Fig. 4i by presenting the results of the FRAP experiments as shown in Fig. III E 3. See Section I F for details of the experimental procedure.

The normalized intensity profiles were fitted to Eq. 3:

$$I(t) = I(\infty) + \left[I(0^+) - I(\infty)\right] \exp\left(-\frac{t}{\tau}\right), \tag{3}$$

where $I(0^+)$ is the fluorescence intensity immediately after bleaching, $I(\infty)$ is the intensity after full recovery, and $\tau$ is the relaxation time. The half-recovery time is given by $t_{1/2} = (\ln 2)\tau$, corresponding to the time required for half of the reduced fluorescence signal to recover.

The apparent diffusion coefficients were extracted using the Soumpasis equation [15]:

$$D_{\text{app}} = 0.224\,\frac{R^2}{t_{1/2}}, \tag{4}$$

where $R$ is the radius of the circular bleached region. The viscosity was then obtained using the Stokes–Einstein relation:

$$\eta = \frac{k_B T}{6\pi D_{\text{app}} R_{\text{Bik1}}}, \tag{5}$$

with $T = 20°\text{C} = 293.15$ K and $R_{\text{Bik1}} = 9.2 \pm 0.5$ nm [6], the radius of gyration obtained by SAXS measurements.

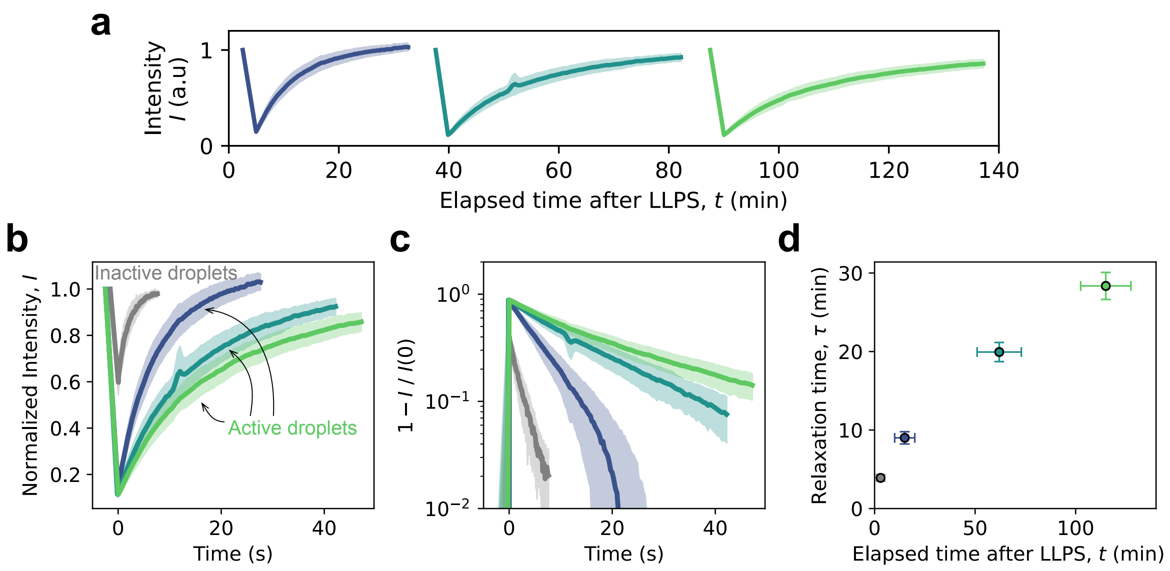


FIG. S9. **Apyrase activity progressively slows molecular mobility in Bik1 condensates. (a)** FRAP recovery curves measured at three distinct times after LLPS, showing a systematic slowing of the relaxation dynamics as the enzymatic reaction proceeds. Curves are averaged over multiple droplets ($n = 7$); shaded regions represent the standard deviation. **(b)** Normalized FRAP curves are shown, corresponding to data in (a). The gray curve corresponds to inactive control droplets containing apyrase but lacking cofactor. **(c)** The semi-log plot shows that the FRAP curves are well described by a single-exponential form, $1 - I/I(0^+) = A\exp(-t/\tau)$. **(d)** Relaxation time $\tau$ is plotted against the eelapsed time after LLPS, indicating a progressive slowdown of molecular mobility within the condensates.

#### IV. THEORY

##### A. Derivation of $\Delta\Delta\mu_i = k_B T \sum_j \frac{s_{ij}}{c_i^{\mathrm{dil}}} \Delta c_j$

Consider a solution of macromolecules (e.g. proteins) in a solvent. We compare two systems: one without solutes, and another containing an additional solute (component $j$). We are interested in how the chemical potential of component $i$ (typically a scaffold protein) in the dilute phase changes upon adding solute. To distinguish compositions in each phase from the total composition, we use superscripts (e.g., $c_i^{\mathrm{dil}}$) to denote the dilute phase.

The chemical potential of component $i$ in the dilute phase is given by

$$\mu_i^{\mathrm{dil}} = \mu_i^{\circ} + k_B T \ln a_i^{\mathrm{dil}},$$

where $a_i^{\mathrm{dil}}$ is the activity of species $i$ in the dilute phase, and $\mu_i^{\circ}$ is the standard chemical potential, defined in the Henry's law reference state (infinite dilution).

In the presence of solutes, the chemical potential in the dilute phase is given by

$$\mu_i'^{\,\mathrm{dil}} = \mu_i^{\circ} + k_B T \ln a_i'^{\,\mathrm{dil}}.$$

The change in the chemical potential $\Delta\Delta\mu$ is then

$$\Delta\Delta\mu \equiv \mu_i'^{\,\mathrm{dil}} - \mu^{\mathrm{dil}} = k_B T \ln \frac{a_i'^{\,\mathrm{dil}}}{a_i^{\mathrm{dil}}}.$$

Assuming ideal behavior, the ratio between the activities reduces to the ratio of concentrations, yielding

$$\Delta\Delta\mu = k_B T \ln \frac{c_i'^{\,\mathrm{dil}}}{c_i^{\mathrm{dil}}}.$$

We now Taylor-expand $c_i'^{\,\mathrm{dil}}$ around $c_i^{\mathrm{dil}}$ to linear order:

$$c_i'^{\,\mathrm{dil}} = c_i^{\mathrm{dil}} + \sum_j s_{ij}(\mathbf{c}^{\mathrm{dil}}) \Delta c_j + O(\Delta c_j^2),$$

where $s_{ij}(\mathbf{c}^{\mathrm{dil}}) \equiv \left(\partial c_i^{\mathrm{dil}} / \partial c_j\right)\big|_{\mathbf{c}^{\mathrm{dil}}}$ is the susceptibility of component $i$ in the dilute phase to solute $j$. Substituting into $\Delta\Delta\mu_i$ yields

$$\Delta\Delta\mu_i = k_B T \sum_j \frac{s_{ij}(\mathbf{c}^{\mathrm{dil}})}{c_i^{\mathrm{dil}}} \Delta c_j + O(\Delta c_j^2). \tag{6}$$

In the phase-separated systems, the chemical potential must be equal across coexisting phases, $\mu_i^{\mathrm{dil}} = \mu_i^{\mathrm{den}}$. Therefore, the change in dense-phase chemical potential due to solute addition, $\Delta\Delta\mu_i^{\mathrm{den}}$, is identical to the dilute-phase expression in Eq. 6. Thus, solute-induced changes in protein–protein interactions within the dense phase can be inferred from the susceptibility $s_{ij}$.

Positive $s_{ij}$ indicates that the solute suppresses LLPS (raising the dilute-phase concentration of scaffold $i$), whereas negative $s_{ij}$ corresponds to solutes that promote LLPS by lowering it.

##### B. Dependence of susceptibility on $c_1^{\mathrm{dil}}$ originates from entropy

This section supplements Fig. 2c and clarifies the thermodynamic origin of the scaling $s \sim c_i^{\mathrm{dil}}$.

In our experiments, phase separation occurs at constant pressure and temperature. Therefore, phase equilibrium is determined by minimizing the Gibbs free energy $G$ rather than the Helmholtz free energy $F$. At constant volume and temperature, however, it is more natural to work with $F$. To avoid confusion, we note that these two approaches are equivalent in our experiments because all samples are prepared at the same total volume: solvent is replaced by added solutes so that the volume is fixed. Consequently, $G = F + PV$ differs from $F$ only by a constant $PV$, and we may equivalently use $F$ to discuss the relation between susceptibility and the underlying free energy.

Suppose that a mixture phase-separates into a dilute phase $\boldsymbol{\phi}^{\mathrm{dil}} = (\phi_1^{\mathrm{dil}}, \phi_2^{\mathrm{dil}}, \ldots, \phi_M^{\mathrm{dil}})$ and a dense phase $\boldsymbol{\phi}^{\mathrm{den}} = (\phi_1^{\mathrm{den}}, \phi_2^{\mathrm{den}}, \ldots, \phi_M^{\mathrm{den}})$. Throughout this paper, we treat the $M$-th component as the solvent. At equilibrium, the chemical potentials of each solute component ($\mu_i \equiv \partial F/\partial N_i$) must be equal between the two phases:

$$\mu_i^{\mathrm{dil}}(\boldsymbol{N}^{\mathrm{dil}}) = \mu_i^{\mathrm{den}}(\boldsymbol{N}^{\mathrm{den}}), \qquad (i = 1, 2, \ldots, M-1), \tag{7}$$

and the osmotic pressure $\Pi$ must also be equal.

These equilibrium conditions are more conveniently expressed in terms of the exchange chemical potential $\hat{\mu}_i \equiv v_i\, \partial f/\partial \phi_i$, where $f \equiv F/V$ and $v_i$ is the molecular volume of component $i$. In terms of $\hat{\mu}_i$, phase equilibrium requires matching the exchange chemical potential

$$\hat{\mu}_i^{\mathrm{dil}}(\boldsymbol{\phi}^{\mathrm{dil}}) = \hat{\mu}_i^{\mathrm{den}}(\boldsymbol{\phi}^{\mathrm{den}}), \qquad (i = 1, 2, \ldots, M-1), \tag{8}$$

and the osmotic pressure

$$\left( f(\boldsymbol{\phi}) - \sum_{j=1}^{M-1} \phi_j \frac{\partial f(\boldsymbol{\phi})}{\partial \phi_j} \right)\Bigg|_{\boldsymbol{\phi}^{\mathrm{dil}}} = \left( f(\boldsymbol{\phi}) - \sum_{j=1}^{M-1} \phi_j \frac{\partial f(\boldsymbol{\phi})}{\partial \phi_j} \right)\Bigg|_{\boldsymbol{\phi}^{\mathrm{den}}}. \tag{9}$$

Now consider infinitesimal shifts of the coexisting compositions,

$$\boldsymbol{\phi}^{\mathrm{dil}} \to \boldsymbol{\phi}^{\mathrm{dil}} + \Delta\boldsymbol{\phi}^{\mathrm{dil}}, \qquad \boldsymbol{\phi}^{\mathrm{den}} \to \boldsymbol{\phi}^{\mathrm{den}} + \Delta\boldsymbol{\phi}^{\mathrm{den}}. \tag{10}$$

Taylor-expanding Eq. 8 to linear order gives

$$\frac{\partial f}{\partial \phi_i}\bigg|_{\boldsymbol{\phi}^{\mathrm{dil}}} + \sum_j \frac{\partial^2 f}{\partial \phi_i \partial \phi_j}\bigg|_{\boldsymbol{\phi}^{\mathrm{dil}}} \Delta\phi_j^{\mathrm{dil}} = \frac{\partial f}{\partial \phi_i}\bigg|_{\boldsymbol{\phi}^{\mathrm{den}}} + \sum_j \frac{\partial^2 f}{\partial \phi_i \partial \phi_j}\bigg|_{\boldsymbol{\phi}^{\mathrm{den}}} \Delta\phi_j^{\mathrm{den}}. \tag{11}$$

The zeroth-order terms cancel due to equilibrium, yielding

$$\sum_j \frac{\partial^2 f}{\partial \phi_i \partial \phi_j}\bigg|_{\boldsymbol{\phi}^{\mathrm{dil}}} \Delta\phi_j^{\mathrm{dil}} = \sum_j \frac{\partial^2 f}{\partial \phi_i \partial \phi_j}\bigg|_{\boldsymbol{\phi}^{\mathrm{den}}} \Delta\phi_j^{\mathrm{den}}. \tag{12}$$

Defining the hessian matrix $\hat{H}_{ij} \equiv \partial^2 f/\partial\phi_i \partial\phi_j$, this condition becomes

$$\hat{H}_{ij}(\boldsymbol{\phi}^{\mathrm{dil}})\Delta\phi_j^{\mathrm{dil}} = \hat{H}_{ij}(\boldsymbol{\phi}^{\mathrm{den}})\Delta\phi_j^{\mathrm{den}}. \tag{13}$$

Now, we apply the same procedure to the osmotic pressure equilibrium condition. Taylor-expanding Eq. 9 to linear order yields

$$\begin{aligned}
&\left[ f + \sum_j \frac{\partial f}{\partial \phi_j} \Delta\phi_j^{\mathrm{dil}} - \sum_j \phi_j \frac{\partial f}{\partial \phi_j} - \sum_j \Delta\phi_j^{\mathrm{dil}} \frac{\partial f}{\partial \phi_j} - \sum_{j,k} \phi_j \frac{\partial^2 f}{\partial \phi_j \partial \phi_k} \Delta\phi_k^{\mathrm{dil}} \right]_{\boldsymbol{\phi}^{\mathrm{dil}}} \\
&= \left[ f + \sum_j \frac{\partial f}{\partial \phi_j} \Delta\phi_j^{\mathrm{den}} - \sum_j \phi_j \frac{\partial f}{\partial \phi_j} - \sum_j \Delta\phi_j^{\mathrm{den}} \frac{\partial f}{\partial \phi_j} - \sum_{j,k} \phi_j \frac{\partial^2 f}{\partial \phi_j \partial \phi_k} \Delta\phi_k^{\mathrm{den}} \right]_{\boldsymbol{\phi}^{\mathrm{den}}}.
\end{aligned} \tag{14}$$

Using Eq. 9 to eliminate the zeroth-order terms, we obtain

$$\sum_{j,k} \phi_j \frac{\partial^2 f}{\partial \phi_j \partial \phi_k} \Delta\phi_k^{\mathrm{dil}} \bigg|_{\boldsymbol{\phi}^{\mathrm{dil}}} = \sum_{j,k} \phi_j \frac{\partial^2 f}{\partial \phi_j \partial \phi_k} \Delta\phi_k^{\mathrm{den}} \bigg|_{\boldsymbol{\phi}^{\mathrm{den}}} . \tag{15}$$

Equivalently, this can be written as

$$\sum_{j,k} \phi_j^{\mathrm{dil}} \hat{H}_{jk}(\boldsymbol{\phi}^{\mathrm{dil}}) \Delta\phi_k^{\mathrm{dil}} = \sum_{j,k} \phi_j^{\mathrm{den}} \hat{H}_{jk}(\boldsymbol{\phi}^{\mathrm{den}}) \Delta\phi_k^{\mathrm{den}}. \tag{16}$$

Combining this result with Eq. 13 to eliminate $\Delta\boldsymbol{\phi}^{\mathrm{den}}$, we arrive at

$$\sum_{j,k} (\phi_j^{\mathrm{den}} - \phi_j^{\mathrm{dil}}) \hat{H}_{jk}(\boldsymbol{\phi}^{\mathrm{dil}}) \Delta\phi_k^{\mathrm{dil}} = 0. \tag{17}$$

Equation 17 must be satisfied for any infinitesimal perturbation of the phase compositions.

To relate $\Delta\boldsymbol{\phi}^{\mathrm{dil}}$ to the change in the total composition $\Delta\boldsymbol{\phi}$, consider a phase-separating system characterized by the overall composition $\boldsymbol{\phi}$ and the coexisting compositions $\boldsymbol{\phi}^{\mathrm{dil}}$ and $\boldsymbol{\phi}^{\mathrm{den}}$. When the total composition is perturbed as $\boldsymbol{\phi} \to \boldsymbol{\phi} + \Delta\boldsymbol{\phi}$, the new equilibrium compositions move along the tie line, defined by the unit vector

$$\hat{\boldsymbol{t}} \equiv \frac{\boldsymbol{\phi}^{\mathrm{den}} - \boldsymbol{\phi}^{\mathrm{dil}}}{|\boldsymbol{\phi}^{\mathrm{den}} - \boldsymbol{\phi}^{\mathrm{dil}}|},$$

until the new dilute-phase composition is reached. This implies

$$\Delta\phi_j^{\mathrm{dil}} = \Delta\phi_j + \alpha \hat{t}_j. \tag{18}$$

Substituting Eq. 18 into Eq. 17 gives

$$\sum_{j,k} \hat{t}_j \hat{H}_{jk} \left( \Delta\phi_k + \alpha \hat{t}_k \right) = 0,$$

which yields

$$\alpha = -\frac{\sum_{j,k} \hat{t}_j \hat{H}_{jk} \Delta\phi_k}{\sum_{j,k} \hat{t}_j \hat{H}_{jk} \hat{t}_k}. \tag{19}$$

Substituting this expression for $\alpha$ back into Eq. 18, we obtain

$$\Delta\phi_i^{\mathrm{dil}} = \Delta\phi_i - \frac{\sum_{j,k} \hat{t}_j \hat{H}_{jk} \Delta\phi_k}{\sum_{j,k} \hat{t}_j \hat{H}_{jk} \hat{t}_k} \hat{t}_i. \tag{20}$$

To relate Eq. 20 to the susceptibility $s_{ij}$, we note that

$$s_{ij} \equiv \frac{\Delta c_i^{\mathrm{dil}}}{\Delta c_j} = \frac{v_j}{v_i} \frac{\Delta\phi_i^{\mathrm{dil}}}{\Delta\phi_j}, \tag{21}$$

where we have used $c_i = N_i/(N_A V) = \phi_i/(N_A v_i)$; $N_A$ is the Avogadro's number.

We now consider how the protein concentration (component 1) in the dilute phase responds to the addition of a solute (component 2), corresponding to a perturbation $\Delta\boldsymbol{\phi} = (0, \Delta\phi_2, 0, 0, \ldots)$. In this case, Eq. 21 gives

$$\boxed{s_{12}(\boldsymbol{\phi}^{\mathrm{dil}}) = -\frac{v_2}{v_1} \frac{\sum_j \hat{t}_j \hat{H}_{j2}(\boldsymbol{\phi}^{\mathrm{dil}}) \hat{t}_1}{\sum_{j,k} \hat{t}_j \hat{H}_{jk}(\boldsymbol{\phi}^{\mathrm{dil}}) \hat{t}_k}.} \quad \text{(general case)} \tag{22}$$

If the protein (component 1) is the only species that partitions between the two phases, so that $t_i = 0$ for $i = 2, 3, \ldots, M$, this expression simplifies to

$$\boxed{s_{12}(\boldsymbol{\phi}^{\text{dil}}) = -\frac{v_2}{v_1}\frac{\hat{H}_{12}(\boldsymbol{\phi}^{\text{dil}})}{\hat{H}_{11}(\boldsymbol{\phi}^{\text{dil}})}.} \quad \text{(no solute partitioning)} \tag{23}$$

It is instructive to evaluate $H_{11} = \partial^2 f/\partial\phi_1^2$ in the dilute limit of component 1. In this case, the chemical potential of component 1 can be approximated as

$$\mu_1 = \mu_1^\circ + k_B T \ln \frac{c_1}{c_0}.$$

Equivalently, the exchange chemical potential can be written as

$$\hat{\mu}_1 \equiv v_1 \frac{\partial f}{\partial \phi_1} = \hat{\mu}_1^\circ + k_B T \ln \phi_1.$$

It follows that

$$\begin{aligned} \hat{H}_{11}(\boldsymbol{\phi}^{\text{dil}}) &= \frac{1}{v_1}\left.\frac{\partial^2 f}{\partial \phi_1^2}\right|_{\boldsymbol{\phi}^{\text{dil}}} = \frac{k_B T}{v_1}\frac{1}{\phi_1^{\text{dil}}} \\ &= \frac{k_B T}{N_A v_1^2}\frac{1}{c_1^{\text{dil}}} \propto \frac{1}{c_1^{\text{dil}}} \qquad \text{(dilute limit)} \end{aligned} \tag{24}$$

Substituting this result into Eq. 23, we obtain

$$\boxed{s_{12} = -\left(\frac{v_2}{v_0}\right) \cdot (N_A v_1 c_1^{\text{dil}}) \cdot \left(\frac{v_0 \hat{H}_{12}(\boldsymbol{c}^{\text{dil}})}{k_B T}\right)} \quad \text{(no solute partitioning, } \phi_1^{\text{dil}} \ll 1) \tag{25}$$

where $v_0$ is the reference volume. We therefore conclude that the entropic contribution alone gives rise to the scaling

$$s = s_{12} \sim c_1^{\text{dil}},$$

and that deviations from this linear dependence originate from departures from the dilute limit and from protein–solute interactions, as encoded in $\hat{H}_{12}$.

##### C. Relationship between susceptibility and Kirkwood–Buff theory

###### 1. *Results at fixed volume $V$ and temperature $T$*

The effects of solutes on biomolecules have historically been discussed in terms of preferential binding. Kirkwood–Buff (KB) theory relates intermolecular spatial distributions in homogeneous mixtures to derivatives of chemical potentials. A useful review is given in Ref. [16]. In particular, KB theory provides a framework for describing solute effects in homogeneous protein solutions, including salting-in and salting-out effects, by connecting changes in chemical potentials to changes in local molecular ordering.

For an open system with the grand canonical ensemble, KB theory [16, 17] gives

$$k_B T \left( \frac{\partial c_i}{\partial \mu_j} \right)_{\{\mu_{i \neq j}\}, V, T} = c_i \delta_{ij} + c_i c_j \tilde{G}_{ij}, \tag{26}$$

where $c_i$ is the molarity of species $i$, $\tilde{G}_{ij} \equiv N_A G_{ij}$, and $G_{ij}$ is the KB integral

$$G_{ij} = G_{ji} = 4\pi \int_0^\infty \left[ g_{ij}(r) - 1 \right] r^2 \, dr.$$

Here $g_{ij}(r)$ is the radial distribution function of species $j$ around species $i$. The KB integral measures the excess of species $j$ around species $i$ relative to a random distribution.

In matrix form, Eq. 26 can be written as

$$H = k_B T \, \mathbf{A}^{-1}, \tag{27}$$

with

$$H_{ij} \equiv \left( \frac{\partial \mu_i}{\partial c_j} \right)_{\{c_{i \neq j}\}, V, T}, \qquad A_{ij} \equiv c_i \delta_{ij} + c_i c_j \tilde{G}_{ij}.$$

We now consider a ternary mixture consisting of protein (1), solute (2), and solvent (3), and derive the susceptibility for the case that the solute does not partition between phases, $k_2 = 1$ using Eq. 23. For the matrix $\mathbf{A}$, the relevant hessian elements are

$$\frac{H_{11}}{k_B T} = \frac{A_{22} A_{33} - A_{23}^2}{D} = \frac{(c_2 + c_2^2 \tilde{G}_{22})(c_3 + c_3^2 \tilde{G}_{33}) - (c_2 c_3 \tilde{G}_{23})^2}{D},$$
$$\frac{H_{12}}{k_B T} = \frac{A_{13} A_{23} - A_{12} A_{33}}{D} = \frac{(c_1 c_3 \tilde{G}_{13})(c_2 c_3 \tilde{G}_{23}) - (c_1 c_2 \tilde{G}_{12})(c_3 + c_3^2 \tilde{G}_{33})}{D},$$

where $D = \det \mathbf{A}$.

To the order up to $c_1^2 c_2$, the determinant is

$$\begin{aligned} D &= A_{11}(A_{22} A_{33} - A_{23}^2) - A_{12}(A_{12} A_{33} - A_{13} A_{23}) + A_{13}(A_{12} A_{23} - A_{13} A_{22}) \\ &\approx c_1 c_2 c_3 (1 + c_3 \tilde{G}_{33}) \left[ 1 + c_1 \left( \tilde{G}_{11} - \frac{c_3 \tilde{G}_{13}^2}{1 + c_3 \tilde{G}_{33}} \right) \right]. \end{aligned}$$

Therefore,

$$\frac{H_{11}}{k_B T} \approx \frac{1}{c_1} - \tilde{G}_{11} + \frac{c_3 \tilde{G}_{13}^2}{1 + c_3 \tilde{G}_{33}}$$
$$\frac{H_{12}}{k_B T} \approx -\tilde{G}_{12} + \frac{c_3 \tilde{G}_{13} \tilde{G}_{23}}{1 + c_3 \tilde{G}_{33}}$$

Plugging these expressions into Eq. 23 yields

$$s_{12} = -\frac{v_2}{v_1}\frac{\hat{H}_{12}(\boldsymbol{\phi}^{\text{dil}})}{\hat{H}_{11}(\boldsymbol{\phi}^{\text{dil}})} = -\frac{v_2}{v_1}\cdot\frac{v_1^2}{v_1 v_2}\frac{H_{12}(\mathbf{c}^{\text{dil}})}{H_{11}(\mathbf{c}^{\text{dil}})} = -\frac{H_{12}(\mathbf{c}^{\text{dil}})}{H_{11}(\mathbf{c}^{\text{dil}})}$$
$$\approx \frac{\tilde{G}_{12}^{\text{dil}} - \dfrac{c_3^{\text{dil}}\tilde{G}_{13}^{\text{dil}}\tilde{G}_{23}^{\text{dil}}}{1+c_3^{\text{dil}}\tilde{G}_{33}^{\text{dil}}}}{\dfrac{1}{c_1^{\text{dil}}} - \tilde{G}_{11}^{\text{dil}} + \dfrac{c_3^{\text{dil}}(\tilde{G}_{13}^{\text{dil}})^2}{1+c_3^{\text{dil}}\tilde{G}_{33}^{\text{dil}}}}. \tag{28}$$

Here, $\hat{H}$ and $H$ describe the same hessian at fixed $V$ and $T$, up to the change of variables between volume fraction and concentration.

According to KB theory [16, 17], the solvent denominator is related to the isothermal compressibility of the solvent,

$$1 + c_3^{\text{dil}}\tilde{G}_{33}^{\text{dil}} = c_3^{\text{dil}} RT\kappa_T. \tag{29}$$

In the dilute limit, the solute–solvent KB integral $G_{i3}$ is related to the infinite-dilution partial molecular volume $v_i^\infty$ by

$$G_{i3} = k_B T\kappa_T - v_i^\infty. \tag{30}$$

Here, $v_i^\infty$ is defined as the volume increment per molecule of species $i$ upon adding it to the solvent. For proteins and small molecules, we approximate $v_i^\infty$ by the molecular volume $v_i$. Since $k_B T\kappa_T = 1.9\times10^{-3}$ nm$^3$ for water at 20°C, this term is small compared with typical molecular volumes. For example, $v_{\text{BSA}} \approx 100$ nm$^3$ and $v_{\text{ATP}} \approx 0.5$ nm$^3$. Thus, $G_{13}^{\text{dil}} \approx -v_1$ and $G_{23}^{\text{dil}} \approx -v_2$.

Using Eqs. 29 and 30, together with $\tilde{G}_{ij} = N_A G_{ij}$, Eq. 28 becomes

$$s_{12} \approx \frac{N_A c_1^{\text{dil}}\left(G_{12}^{\text{dil}} - \dfrac{v_1 v_2}{k_B T\kappa_T}\right)}{1 - N_A c_1^{\text{dil}}\left(G_{11}^{\text{dil}} - \dfrac{v_1^2}{k_B T\kappa_T}\right)} = \frac{\phi_1^{\text{dil}}\left(\dfrac{G_{12}^{\text{dil}}}{v_1} - \dfrac{v_2}{k_B T\kappa_T}\right)}{1 - \phi_1^{\text{dil}}\left(\dfrac{G_{11}^{\text{dil}}}{v_1} - \dfrac{v_1}{k_B T\kappa_T}\right)}. \tag{31}$$

Here, $\phi_1^{\text{dil}} = N_A c_1^{\text{dil}} v_1$.

The terms $v_i/(k_B T\kappa_T)$ are large because water has a small compressibility. For BSA, $v_1/(k_B T\kappa_T) \approx 5\times10^4$, while for ATP, $v_2/(k_B T\kappa_T) \approx 260$. These terms reflect solvent compression as solutes are added while the system volume is held fixed. A simple mechanical analogy is that the work required to compress a solvent volume $V$ by a volume $v$ scales as $(1/2)v^2/(\kappa_T V)$.

When $v_i/(k_B T\kappa_T) \gg 1$ and $\phi_1^{\text{dil}} v_1/(k_B T\kappa_T) \gg 1$, Eq. 31 reduces to

$$\boxed{s_{12} \approx -\frac{v_2}{v_1} + \frac{k_B T\kappa_T}{v_1}\frac{G_{12}^{\text{dil}}}{v_1} \qquad \text{at fixed } V, T.} \tag{32}$$

In this fixed-volume, susceptibility $s_{12}$ is essentially equal to $v_2/v_1$ as $k_B T\kappa/v_1$ is small.

It is pedagogical to consider the case where protein (1) and solute (2) interact as hard spheres. The pair radial distribution function is

$$g_{ij}(r) = \begin{cases} 0, & r < R_i + R_j, \\ 1, & r > R_i + R_j. \end{cases}$$

Therefore,

$$\begin{aligned} G_{ij} &= 4\pi \int_0^\infty (g_{ij}(r) - 1)\, r^2\, dr \\ &= 4\pi \int_0^{R_i+R_j} (-1) r^2\, dr \\ &= -\frac{4\pi}{3}(R_i + R_j)^3. \end{aligned}$$

Using $v_1 = (4\pi/3)R_1^3$, this gives $G_{12}/v_1 = -(1+R_2/R_1)^3$. Therefore, in this fixed-volume, solvent-compression-dominated regime, hard spheres give

$$\boxed{s_{12} \approx -\left[\frac{v_2}{v_1} + \frac{k_B T \kappa_T}{v_1}\left(1 + \left(\frac{v_2}{v_1}\right)^{\frac{1}{3}}\right)^3\right] \qquad \text{at fixed } V, T \text{ (hard spheres).}} \tag{33}$$

*2. Results at fixed pressure P and temperature T*

In the fixed-volume formulation, solvent compression can dominate the susceptibility and eliminate the explicit dependence on $c_1^{\mathrm{dil}}$ or $\phi_1^{\mathrm{dil}}$ (see Eq. 32). We now apply Kirkwood–Buff theory to derive the corresponding result at fixed pressure and temperature, which is the more relevant condition for experiments.

The susceptibility expressions in Eqs. 22 and 23 remain valid at fixed $P, T$, provided that the hessian is evaluated under the same thermodynamic constraint. This is because that Eqs. 22 and 23 are derived by equating the chemical potentials between two phases, and the chemical potentials in the $NVT$ and $NPT$ ensembles are identical.

$$\left(\frac{\partial F}{\partial N_i}\right)_{V,T,\{N_{j\neq i}\}} = \left(\frac{\partial G}{\partial N_i}\right)_{P,T,\{N_{j\neq i}\}} = \mu_i.$$

However, its derivative with respect to composition depends on the constraint. We denote the fixed-$V, T$ hessian by

$$H_{ij} \equiv \left(\frac{\partial \mu_i}{\partial c_j}\right)_{V,T},$$

and the fixed-$P, T$ hessian by

$$\mathcal{H}_{ij} \equiv \left(\frac{\partial \mu_i}{\partial c_j}\right)_{P,T}.$$

As derived in Appendix A, these hessians are related by

$$\mathcal{H} = H - \frac{(H\mathbf{c})(H\mathbf{c})^{\mathsf{T}}}{\mathbf{c}^{\mathsf{T}} H \mathbf{c}}. \tag{34}$$

The correction term in Eq. 34 for the $(1, 2)$ element is

$$\frac{1}{k_B T}\frac{\left[(H\mathbf{c})(H\mathbf{c})^{\mathsf{T}}\right]_{12}}{\mathbf{c}^{\mathsf{T}} H \mathbf{c}} \approx \frac{\left(1 + c_3\tilde{G}_{33} - c_3\tilde{G}_{13}\right)\left(1 + c_3\tilde{G}_{33} - c_3\tilde{G}_{23}\right)}{c_3\left(1 + c_3\tilde{G}_{33}\right)}.$$

Therefore,

$$\frac{\mathcal{H}_{12}}{k_BT} \approx -\tilde{G}_{12} + \frac{c_3\tilde{G}_{13}\tilde{G}_{23}}{1+c_3\tilde{G}_{33}} - \frac{\left(1+c_3\tilde{G}_{33}-c_3\tilde{G}_{13}\right)\left(1+c_3\tilde{G}_{33}-c_3\tilde{G}_{23}\right)}{c_3\left(1+c_3\tilde{G}_{33}\right)}.$$

Expanding the last term, $c_3\tilde{G}_{13}\tilde{G}_{23}/(1+c_3\tilde{G}_{33})$ cancels exactly, giving

$$\frac{\mathcal{H}_{12}}{k_BT} \approx -\tilde{G}_{12} + \tilde{G}_{13} + \tilde{G}_{23} - \tilde{G}_{33} - \frac{1}{c_3}.$$

Using Eq. 29, we obtain

$$\frac{\mathcal{H}_{12}}{k_BT} \approx -\tilde{G}_{12} + \tilde{G}_{13} + \tilde{G}_{23} - RT\kappa_T.$$

For the $(1,1)$ element,

$$\mathcal{H}_{11} = H_{11} - \frac{\left(H_{11}c_1 + H_{12}c_2 + H_{13}c_3\right)^2}{\sum_{kl} c_k H_{kl} c_l}. \tag{35}$$

In the dilute limit, $c_1, c_2 \to 0$ while $c_3$ remains finite. Keeping the finite terms in $H_{11}$ that are of the same order as the pressure correction, we find

$$\frac{\mathcal{H}_{11}}{k_BT} \approx \left(\frac{1}{c_1} - \tilde{G}_{11} + \frac{c_3\tilde{G}_{13}^2}{1+c_3\tilde{G}_{33}}\right) - \frac{\left(1+c_3\tilde{G}_{33}-c_3\tilde{G}_{13}\right)^2}{c_3\left(1+c_3\tilde{G}_{33}\right)} \tag{36}$$

$$= \frac{1}{c_1} - \tilde{G}_{11} + 2\tilde{G}_{13} - \frac{1+c_3\tilde{G}_{33}}{c_3}. \tag{37}$$

Using Eq. 29, this becomes

$$\frac{\mathcal{H}_{11}}{k_BT} \approx \frac{1}{c_1} - \tilde{G}_{11} + 2\tilde{G}_{13} - RT\kappa_T. \tag{38}$$

Thus, under fixed pressure and temperature, the susceptibility without solute partitioning is given by

$$s_{12} = -\frac{\mathcal{H}_{12}}{\mathcal{H}_{11}} \tag{39}$$

$$\approx -\frac{-\tilde{G}_{12} + \tilde{G}_{13} + \tilde{G}_{23} - RT\kappa_T}{\frac{1}{c_1} - \tilde{G}_{11} + 2\tilde{G}_{13} - RT\kappa_T}. \tag{40}$$

In the dilute protein limit, the leading term in the denominator is $1/c_1$, yielding

$$s_{12} \approx c_1^{\text{dil}}\left(\tilde{G}_{12}^{\text{dil}} - \tilde{G}_{13}^{\text{dil}} - \tilde{G}_{23}^{\text{dil}} + RT\kappa_T\right) \tag{41}$$

$$= N_A c_1^{\text{dil}}\left(G_{12}^{\text{dil}} - G_{13}^{\text{dil}} - G_{23}^{\text{dil}} + k_BT\kappa_T\right). \tag{42}$$

Using Eq. 30 and assuming $v_i^\infty \approx v_i$, we obtain

$$\boxed{\begin{aligned} s_{12} &\approx N_A c_1^{\text{dil}}\left(G_{12}^{\text{dil}} + v_1 + v_2 - k_BT\kappa_T\right) \\ &= \phi_1^{\text{dil}}\left(1 + \frac{G_{12}^{\text{dil}}}{v_1} + \frac{v_2}{v_1} - \frac{k_BT\kappa_T}{v_1}\right) \qquad \text{at fixed } P,T. \end{aligned}} \tag{43}$$

If the protein (1) and solute (2) are treated as hard spheres, then

$$\frac{G_{12}}{v_1} = -\left(1 + \frac{R_2}{R_1}\right)^3 .$$

Using $v_2/v_1 = (R_2/R_1)^3$, the bracket in Eq. 43 becomes

$$\begin{aligned} 1 + \frac{G_{12}^{\mathrm{dil}}}{v_1} + \frac{v_2}{v_1} - \frac{k_B T \kappa_T}{v_1} &= 1 - \left(1 + \frac{R_2}{R_1}\right)^3 + \left(\frac{R_2}{R_1}\right)^3 - \frac{k_B T \kappa_T}{v_1} \\ &= -3\frac{R_2}{R_1} - 3\left(\frac{R_2}{R_1}\right)^2 - \frac{k_B T \kappa_T}{v_1}. \end{aligned}$$

Therefore,

$$\boxed{s_{12} \approx -\phi_1^{\mathrm{dil}} \left[3\left(\frac{v_2}{v_1}\right)^{1/3} + 3\left(\frac{v_2}{v_1}\right)^{2/3} + \frac{k_B T \kappa_T}{v_1}\right] < 0 \qquad \text{at fixed } P, T \text{ (hard spheres).}} \tag{44}$$

In the hard-sphere model, the susceptibility is always negative. Adding solute favors condensation because clustered proteins exclude less total volume to the solute than dispersed proteins. For example, $v_1 = v_{\mathrm{BSA}} \approx 100\ \mathrm{nm}^3$, $v_2 = v_{\mathrm{ATP}} = 0.5\ \mathrm{nm}^3$, and $k_B T \kappa_T = 1.9 \times 10^{-3}\ \mathrm{nm}^3$ for water at 20°C. This makes the last term $\mathcal{O}(10^{-5})$ is negligible. For ATP and BSA, $v_2/v_1 \simeq 4.9 \times 10^{-3}$, leading to $s_{12} \approx -0.6\phi_1^{\mathrm{dil}}$. The coefficient in Eq. 44 without the solvent-compression term is shown in Fig. S10.

To summarize, solutes must be sufficiently enriched around proteins to promote condensation by overcoming the excluded-volume effect.

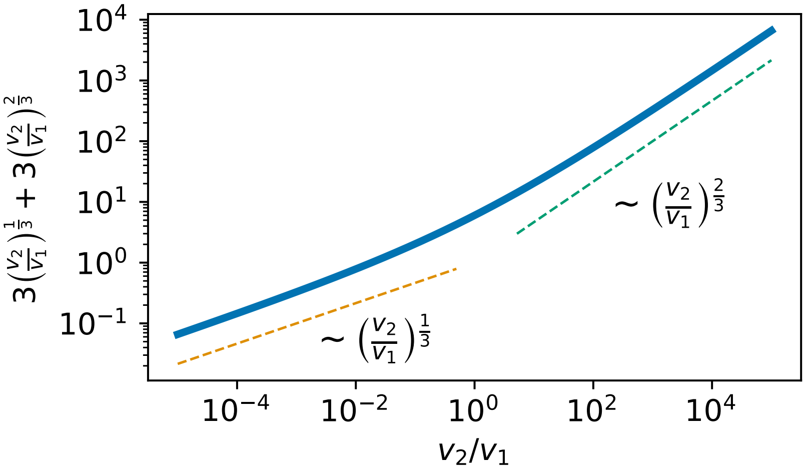


FIG. S10. **Volume dependence of susceptibility from Kirkwood–Buff theory with hard-sphere interactions.** The dimensionless coefficient of the susceptibility in Eq. 44 is plotted. In this model, proteins (1) and solutes (2) interact as hard spheres. For small solutes, the coefficient scales as $(v_2/v_1)^{1/3}$, while for large solutes it scales as $(v_2/v_1)^{2/3}$. The crossover occurs around $v_2/v_1 = 10^{-1}$–$10^1$.

##### D. Geometrical interpretation of susceptibility $s$

From a geometrical perspective, adding solute moves a thermodynamic state along the binodal manifold $\mathcal{B}$ in composition space. For a ternary mixture, the free-energy landscape spans three dimensions and the binodal is a two-dimensional manifold $\mathcal{B} \subset \mathbb{R}^3$ that separates one-phase and two-phase regions. The projection of $\mathcal{B}$ onto the $c_1$–$c_2$ plane shifts the binodal curve as $c_3$ increases; these shifts are encoded in $s_{ij}$.

###### 1. *Relationship among susceptibility $s$, binodal gradient $r$, and tie-line slope $t$ in two-component systems*

We consider a binary system $(c_1, c_2)$ near the binodal dilute branch. Let the dense-phase volume fraction be $\Phi = V^{\text{den}}/V$ with $\Phi \ll 1$. The mass balance reads

$$c_1^{\text{tot}} = c_1^{\text{dil}} + \Phi(c_1^{\text{den}} - c_1^{\text{dil}}) = c_1^{\text{dil}} + \Phi t_1, \tag{45}$$

$$c_2^{\text{tot}} = c_2^{\text{dil}} + \Phi(c_2^{\text{den}} - c_2^{\text{dil}}) = c_2^{\text{dil}} + \Phi t_2, \tag{46}$$

with $t_1 = c_1^{\text{den}} - c_1^{\text{dil}}$, $t_2 = c_2^{\text{den}} - c_2^{\text{dil}}$, and $t = t_1/t_2$ (tie-line slope). The binodal gradient is defined as

$$r \equiv \left.\frac{\partial c_1}{\partial c_2}\right|_{\mathbf{c}\in\mathcal{B}},$$

and the susceptibility as

$$s \equiv \left(\frac{\partial c_1^{\text{dil}}}{\partial c_2^{\text{tot}}}\right)_{c_1^{\text{tot}}}. \tag{47}$$

The total differential of $c_1^{\text{tot}}$ is given by

$$dc_1^{\text{tot}} = dc_1^{\text{dil}} + t_1\, d\Phi + \Phi\, dt_1. \tag{48}$$

We consider variations at fixed total composition ($dc_1^{\text{tot}} = 0$) in the vicinity of the dilute-phase composition (where $\Phi \ll 1$). Keeping only leading-order terms in $\Phi$, the contribution $\Phi dt_1$ can be neglected, yielding

$$0 = dc_1^{\text{dil}} + t_1\, d\Phi \quad \Rightarrow \quad dc_1^{\text{dil}} = -t_1\, d\Phi. \tag{49}$$

From Eq. 46, we similarly obtain

$$dc_2^{\text{tot}} = dc_2^{\text{dil}} + t_2\, d\Phi. \tag{50}$$

On the binodal, we note that

$$dc_1^{\text{dil}} = r\, dc_2^{\text{dil}} \quad \Leftrightarrow \quad dc_2^{\text{dil}} = \frac{1}{r}\, dc_1^{\text{dil}}. \tag{51}$$

Eliminating $d\Phi$ using Eq. 49 and substituting into Eq. 50 with Eq. 51, we obtain

$$dc_2^{\text{tot}} = dc_1^{\text{dil}} \left(\frac{1}{r} - \frac{1}{t}\right). \tag{52}$$

At fixed $c_1^{\rm tot}$ and in the limit $\Phi \to 0$, the susceptibility reduces to $s = dc_1^{\rm dil}/dc_2^{\rm tot}$, yielding the exact relation

$$\frac{1}{r} = \frac{1}{s} + \frac{1}{t}. \tag{53}$$

This relation holds only in the vicinity of the binodal, where $\Phi \ll 1$. This identity links the susceptibility $s$ to the binodal gradient $r$ and the tie-line slope $t$. This relation is equivalent to Eq. 27 in the Supplementary Information of Ref. [18] and Eq. 6 in [19], where the authors use the normal to the phase boundary rather than the tangent. A refined form appears in Eq. 3 of Ref. [20]. In that work, the authors use the equation to define a quantity called dominance,

$$\mathcal{D} \equiv \frac{t}{t-r} = \frac{s}{r} \quad \because \text{Eq. 53},$$

which describes the relative energetic contribution of the protein-protein interactions to the overall free-energy gain upon phase separation. The dominance can be given by the ratio of the susceptibility $s$ to the binodal gradient $r$. Because determining the tie-line slope requires measurements in both the dilute and dense phases, this ratio provides the simplest way to quantify the dominance.

An important implication is that this relation enables the indirect inference of partition coefficient of component 2 (solute), $k_2 = \phi_2^{\rm den}/\phi_2^{\rm dil}$, given by

$$k_2 = \frac{c_2^{\rm den}}{c_2^{\rm dil}} = 1 + \left(\frac{1}{r} - \frac{1}{s}\right)(k-1) = 1 + \frac{(\mathcal{D}-1)(k-1)}{s},$$

where $k = c_1^{\rm den}/c_1^{\rm dil}$ is the partition coefficient of component 1. The primary difficulty in quantifying $k_2$ lies in measuring the dilute-phase solute concentration $c_2^{\rm dil}$, which is required to determine $r$.

This geometric relation between $r$, $s$, and $t$ holds for any binary system undergoing phase separation. Figure S11a illustrates a representative example: segregative LLPS driven by macromolecular crowding, computed using a free-volume theory with a macromolecule–depletant size ratio of 2 (see SI §IV F). All three quantities vary along the binodal and can be parameterized by the partition coefficient of component 1 (protein), as shown in Fig. S11b. Segregative LLPS is characterized by $r$, $s$, and $t$ all being negative. For associative LLPS, other sign combinations are possible as long as Eq. 53 is satisfied. Figure S11c confirms the relation $1/r = 1/s + 1/t$.

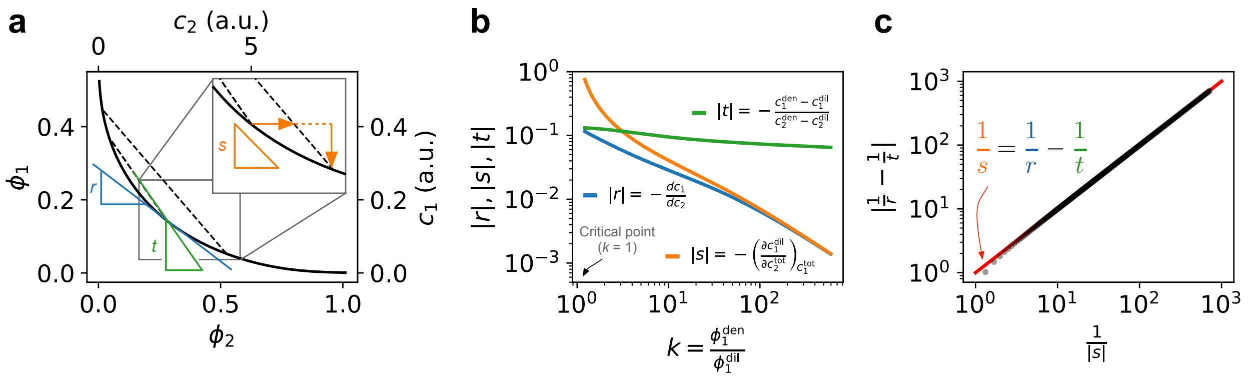


FIG. S11. **Geometric relationship between the binodal gradient $r$, the dilute-phase susceptibility $s$, and the tie-line slope $t$.** **(a)** Phase diagram of a binary mixture undergoing segregative phase separation, simulated using scaled particle theory with depletion interactions with the size ratio $R_2/R_1$ of 0.5. The dashed lines indicate tie-lines; their slopes define $t$. Three slopes are highlighted: the binodal gradient $r = \mathrm{d}c_1/\mathrm{d}c_2$, the tie-line slope $t = (c_1^{\mathrm{den}} - c_1^{\mathrm{dil}})/(c_2^{\mathrm{den}} - c_2^{\mathrm{dil}})$, and the dilute-phase susceptibility $s = (\partial c_1^{\mathrm{dil}}/\partial c_2^{\mathrm{tot}})_{c_1^{\mathrm{tot}}}$, which quantifies how the dilute-phase composition responds to added solute. **(b)** Magnitudes of $r$, $s$, and $t$ plotted against the partition coefficient $k = \phi_1^{\mathrm{den}}/\phi_1^{\mathrm{dil}}$. **(c)** Plot of $|1/s|$ versus $|1/r - 1/t|$. The data collapse onto the line $y = x$ (red), confirming the relation $1/s = 1/r - 1/t$ predicted by mass balance and tie-line geometry.

*2. Extension to N-component systems that separate into two phases*

The relation between the susceptibility $s$, binodal gradient $r$, and tie-line slope $t$ holds for multi-component systems that phase separates into two phases. To generalize from the binary case, it is useful to first clarify the geometry of the concentration space. Let $\mathbf{c} = (c_1, \ldots, c_N) \in \mathbb{R}^N$ denote the molar concentrations of the $N$ components. Note that the free energy density $f = f(\mathbf{c})$ is an $N$-dimensional manifold, and the binodal manifold $\mathcal{B}$ is an $(N-1)$-dimensional surface in $\mathbb{R}^N$ separating single- and two-phase regions. At a given point on $\mathcal{B}$, the tangent vector $\mathbf{r} \in T_{\mathbf{c}}\mathcal{B}$ specifies how the dilute-phase composition changes along the phase coexistence boundary. In contrast, the coexisting dilute and dense phases are connected by a single tie-line, whose direction is given by the tie-line vector

$$\mathbf{t} = \mathbf{c}^{\mathrm{den}} - \mathbf{c}^{\mathrm{dil}} \in \mathbb{R}^N.$$

The susceptibility is generalized from the scalar $s$ of the binary case to the Jacobian matrix $S$:

$$s_{ij} = \left(\frac{\partial c_i^{\mathrm{dil}}}{\partial c_j^{\mathrm{tot}}}\right)_{c_i^{\mathrm{tot}}=\mathrm{const.}, i\neq j}, \qquad S = (s_{ij}) \in \mathbb{R}^{N\times N}.$$

By the lever rule, for infinitesimal variations near the binodal (as $\Phi \to 0$), the total and dilute concentrations are related by

$$d\mathbf{c}^{\mathrm{tot}} = d\mathbf{c}^{\mathrm{dil}} + \mathbf{t}\, d\Phi, \tag{54}$$

while coexistence along the binodal enforces

$$d\mathbf{c}^{\mathrm{dil}} = \alpha \mathbf{r}. \tag{55}$$

This is because the dilute phase is restricted to remain on the binodal manifold, any infinitesimal change in its composition is tangent to that manifold, hence parallel to the binodal gradient $\mathbf{r}$. The scalar coefficient $\alpha$ is determined by the imposed change in the total concentrations $\boldsymbol{\delta} = d\mathbf{c}^{\mathrm{tot}}$, and is given explicitly by

$$\alpha = \frac{(\mathbf{t}\cdot\mathbf{t})\,(\mathbf{r}\cdot\boldsymbol{\delta}) - (\mathbf{r}\cdot\mathbf{t})\,(\mathbf{t}\cdot\boldsymbol{\delta})}{D},$$

where $D = (\mathbf{r}\cdot\mathbf{r})(\mathbf{t}\cdot\mathbf{t}) - (\mathbf{r}\cdot\mathbf{t})^2 > 0$ is the determinant of the Gram matrix of $\mathbf{r}$ and $\mathbf{t}$. Eliminating $d\Phi$ from Eqs. 54–55 yields a linear map between $d\mathbf{c}^{\mathrm{tot}}$ and $d\mathbf{c}^{\mathrm{dil}}$. Explicitly,

$$d\mathbf{c}^{\mathrm{dil}} = S\, d\mathbf{c}^{\mathrm{tot}}, \qquad S = \mathbf{r}\,\mathbf{w}^{\top},$$

with

$$\mathbf{w} = \frac{(\mathbf{t}\cdot\mathbf{t})\,\mathbf{r} - (\mathbf{r}\cdot\mathbf{t})\,\mathbf{t}}{D}.$$

Thus the Jacobian elements are

$$s_{ij} = r_i\, w_j = \frac{r_i\big[(\mathbf{t}\cdot\mathbf{t})\, r_j - (\mathbf{r}\cdot\mathbf{t})\, t_j\big]}{D}. \tag{56}$$

For $N = 2$, Eq. 56 reduces to the relation for the binary system (Eq. 53)

$$\frac{1}{r} = \frac{1}{s} + \frac{1}{t},$$

demonstrating that the Jacobian $S$ in higher dimensions is the natural generalization of the scalar response $s$.

#### E. Susceptibility to promiscuous (non-specific) interactions

This section supplements the discussion of susceptibilities arising from promiscuous (non-specific) interactions, where solutes interact weakly with both proteins and solvent. Consequently, their effects on phase equilibria are well described by a mean-field model. **Flory–Huggins theory** provides a natural starting point, capturing the generic features of such weak interactions. First, we derive the susceptibility within the Flory–Huggins framework using Eq. 23 in the case that solutes do not partition into condensates. We then develop a perturbative formulation that incorporates partitioning.

##### 1. *Estimation of s using the Flory Huggins theory without solute partitioning*

We derive the expression of the susceptibility using the Flory-Huggins free energy function in the case that solutes do not partition. In the dilute limit, recall the Eq. 23:

$$s_{12}(\boldsymbol{\phi}^{\text{dil}}) = -\frac{v_2}{v_1}\frac{\hat{H}_{12}(\boldsymbol{\phi}^{\text{dil}})}{\hat{H}_{11}(\boldsymbol{\phi}^{\text{dil}})}$$

The Flory–Huggins free energy density at fixed temperature $T$ is given by

$$f(\boldsymbol{\phi}) = \frac{k_B T}{v_0}\left(\sum_i \frac{v_0}{\tilde{v}_i}\frac{\phi_i}{N_i}\ln\phi_i + \frac{1}{2}\sum_{i,j}\chi_{ij}\phi_i\phi_j\right), \tag{57}$$

where $\phi_i$ is volume fraction of component $i$ with $\sum_i \phi_i = 1$, $v_0$ is a reference volume, and $N_i$ is the number of segments per molecule. The quantity $\tilde{v}_i$ denotes the segment (monomer) volume.

For a $M$-component system, its hessian is given by

$$\hat{H}_{ij}(\boldsymbol{\phi}) = \frac{\partial f(\boldsymbol{\phi})}{\partial\phi_i\partial\phi_j} = \frac{k_B T}{v_0}\left(\frac{v_0}{\tilde{v}_i N_i}\frac{\delta_{ij}}{\phi_i} + \frac{v_0}{\tilde{v}_M N_M}\frac{1}{\phi_M} + \chi_{ij} - \chi_{iM} - \chi_{Mj} + \chi_{MM}\right), \tag{58}$$

where $\delta_{ij}$ is the Kronecker delta, and the $M$-th component is the solvent such that $\phi_M = 1 - \sum_{i=1}^{M-1}\phi_i$. In practice, the prefactor $v_0/(\tilde{v}_M N_M)$ is typically set to unity, and the diagonal elements of the $\chi$ maxtrix is set to zero ($\chi_{MM} = 0$).

Note that

$$\hat{H}_{11} = \frac{k_B T}{v_0}\left(\frac{v_0}{\tilde{v}_1 N_1}\frac{1}{\phi_1} + \frac{1}{\phi_M}\right); \qquad \hat{H}_{12} = \frac{k_B T}{v_0}\left(\frac{1}{\phi_M} + \chi_{12} - \chi_{1M} - \chi_{M2}\right)$$

We obtain

$$s_{12}(\boldsymbol{\phi}^{\text{dil}}) = -\frac{v_2}{v_1}\frac{\frac{1}{\phi_M} + \chi_{12} - \chi_{1M} - \chi_{M2}}{\frac{v_0}{\tilde{v}_1 N_1}\frac{1}{\phi_1^{\text{dil}}} + \frac{1}{\phi_M}}.$$

In the dilute limit, the first term in $\hat{H}_{11}$ significantly larger than the second term, so we can drop the second term $1/\phi_M$. In the ternary system, we shall consider the effect of the solute (component 2) to protein (component 1). The susceptibility is then

$$\begin{aligned} s_{12} &\approx -\left(\frac{v_2}{v_0}\right)\left(\frac{\tilde{v}_1 N_1}{v_1}\right)\phi_1^{\text{dil}}\Big(\underbrace{\frac{1}{1-\phi_1-\phi_2}}_{\approx 1 \text{ (dilute solution)}} + \underbrace{\chi_{12}}_{\text{protein–solute}} - \underbrace{\chi_{13}}_{\text{protein–solvent}} - \underbrace{\chi_{32}}_{\text{solute–solvent}}\Big) \\ &\approx -\left(\frac{v_2}{v_0}\right)\left(N_A v_1 c_1^{\text{dil}}(\mathbf{c})\right)(1+\chi^{\Delta}). \end{aligned} \tag{59}$$

where $\chi^{\Delta} \equiv \chi_{12} - \chi_{13} - \chi_{23}$, using the symmetry $\chi_{23} = \chi_{32}$. In the last step, $v_1 = \tilde{v}_1 N_1$ is assumed. That is, the effective volume of molecule matches the simple multiple of segment volume. The second factor in Eq. 59 corresponds to the volume fraction of component 1 in the dilute phase, $\phi_1^{\text{dil}}(\mathbf{c}) = N_A v_1 c_1^{\text{dil}}(\mathbf{c})$. A larger $\chi^{\Delta}$ indicates more repulsive (less favorable) protein–solute interactions. Consequently, when protein–solute interactions are more repulsive than the combined protein–solvent and solute–solvent interactions, solutes promote phase separation. Conversely, when solutes preferentially interact with proteins relative to the solvent, they favor dissolution.

*2. Estimation of s using the Flory Huggins theory with solute partitioning*

Eq. 59 neglects the effect of solute partitioning. In this section, we derive $s$ when solute partitions into condensates. We consider an incompressible ternary mixture consisting of components 1 (scaffold), 2 (solute), and 3 (solvent). Its Helmholtz free-energy density at temperature $T$ is given by

$$f(\boldsymbol{\phi} = (\phi_1, \phi_2)) = \frac{k_B T}{v_0} \Big[ \frac{v_0}{\tilde{v}_1} \frac{1}{N_1} \phi_1 \ln \phi_1 + \frac{v_0}{\tilde{v}_2} \frac{1}{N_2} \phi_2 \ln \phi_2 + (1 - \phi_1 - \phi_2) \ln(1 - \phi_1 - \phi_2) + \chi_{13}\phi_1(1 - \phi_1 - \phi_2) + \chi_{23}\phi_2(1 - \phi_1 - \phi_2) + \chi_{12}\phi_1\phi_2 \Big]. \tag{60}$$

Phase coexistence between a scaffold-poor (dilute, $\alpha$) and scaffold-rich (dense, $\beta$) phase is determined by the equality of (exchange) chemical potentials, $\hat{\mu} \equiv \partial f / \partial \phi_i = \mu_i / \tilde{v}_i$,

$$\hat{\mu}_i(\boldsymbol{\phi}^{\alpha}) = \hat{\mu}_i(\boldsymbol{\phi}^{\beta}), \qquad i = 1, 2, \tag{61}$$

and of osmotic pressure $\Pi = -f(\boldsymbol{\phi}) + \hat{\boldsymbol{\mu}} \cdot \boldsymbol{\phi} + f(\boldsymbol{\phi} = 0)$,

$$f(\boldsymbol{\phi}^{\beta}) - f(\boldsymbol{\phi}^{\alpha}) = \hat{\boldsymbol{\mu}}(\boldsymbol{\phi}^{\alpha}) \cdot (\boldsymbol{\phi}^{\beta} - \boldsymbol{\phi}^{\alpha}), \tag{62}$$

The incompressibility condition gives a lever rule:

$$\phi_i = (1 - \Phi)\phi_i^{\alpha} + \Phi\phi_i^{\beta}, \qquad i = 1, 2, \tag{63}$$

where $\Phi = V^{\beta}/V$ is the volume fraction of the dense phase $\beta$. These equations are sufficient to determine the unknowns $(\phi_1^{\alpha}, \phi_2^{\alpha}, \phi_1^{\beta}, \phi_2^{\beta}, \Phi)$.

Now we consider a small perturbation $\Delta\phi_2$ at equilibrium as the system phase-separates into two phases at $\hat{\phi}_i^{\alpha}$ and $\hat{\phi}_i^{\beta}$. This small perturbation is exhibited differently in each phase $\Delta\phi_2^{\text{phase}}$.

$$\phi_1^{\text{phase}} = \hat{\phi}_1^{\text{phase}} + \sigma^{\text{phase}} \Delta\phi_2^{\text{phase}} \qquad \text{phase} : \alpha, \beta$$

Linearizing the left-hand side of Eq. 62 yields

$$\delta(f(\boldsymbol{\phi}^{\alpha}) - f(\boldsymbol{\phi}^{\alpha})) = \hat{\boldsymbol{\mu}}(\hat{\boldsymbol{\phi}}^{\beta}) \cdot (\Delta\hat{\boldsymbol{\phi}}^{\beta} - \Delta\hat{\boldsymbol{\phi}}^{\alpha}). \tag{64}$$

Linearizing the right-hand side at $\boldsymbol{\phi} = \boldsymbol{\phi}^{\alpha}$ yields

$$\delta \left[ \hat{\boldsymbol{\mu}}(\boldsymbol{\phi}^{\alpha}) \cdot (\boldsymbol{\phi}^{\beta} - \boldsymbol{\phi}^{\alpha}) \right] = \delta\hat{\boldsymbol{\mu}}(\hat{\boldsymbol{\phi}}^{\alpha}) \cdot (\hat{\boldsymbol{\phi}}^{\beta} - \hat{\boldsymbol{\phi}}^{\alpha}) + \boldsymbol{\mu}(\hat{\boldsymbol{\phi}}^{\alpha}) \cdot (\Delta\hat{\boldsymbol{\phi}}^{\beta} - \Delta\hat{\boldsymbol{\phi}}^{\alpha}). \tag{65}$$

Here $\delta\hat{\mu}_i^{\text{phase}}$ is the linear change in the chemical potential, given by

$$\delta\hat{\mu}_i^{\text{phase}} = \sum_{j=1}^{2} \hat{H}_{ij}(\hat{\boldsymbol{\phi}}^{\text{phase}}) \Delta\phi_j^{\text{phase}}.$$

where $\hat{H}_{ij}(\boldsymbol{\phi}) = \frac{\partial \hat{\mu}_i}{\partial \phi_j} = \frac{\partial^2 f}{\partial \phi_i \partial \phi_j}$ is the hessian of free energy density. Equating Eqs. 64 and 65 yields

$$\left(\hat{H}_{11}(\hat{\boldsymbol{\phi}}^\alpha)\Delta\phi_1^\alpha + \hat{H}_{12}(\hat{\boldsymbol{\phi}}^\alpha)\Delta\phi_2^\alpha\right)\left(\hat{\phi}_1^\beta - \hat{\phi}_1^\alpha\right) + \left(\hat{H}_{21}(\hat{\boldsymbol{\phi}}^\alpha)\Delta\phi_1^\alpha + \hat{H}_{22}(\hat{\boldsymbol{\phi}}^\alpha)\Delta\phi_2^\alpha\right)\left(\hat{\phi}_2^\beta - \hat{\phi}_2^\alpha\right) = 0.$$

With $\Delta\phi_1^\alpha = \sigma^\alpha \Delta\phi_2^\alpha$, we obtain

$$\sigma^\alpha = \frac{\Delta\phi_1^\alpha}{\Delta\phi_2^\alpha} = -\frac{\hat{H}_{12}(\hat{\boldsymbol{\phi}}^\alpha)(\hat{\phi}_1^\beta - \hat{\phi}_1^\alpha) + \hat{H}_{22}(\hat{\boldsymbol{\phi}}^\alpha)(\hat{\phi}_2^\beta - \hat{\phi}_2^\alpha)}{\hat{H}_{11}(\hat{\boldsymbol{\phi}}^\alpha)(\hat{\phi}_1^\beta - \hat{\phi}_1^\alpha) + \hat{H}_{21}(\hat{\boldsymbol{\phi}}^\alpha)(\hat{\phi}_2^\beta - \hat{\phi}_2^\alpha)}. \tag{66}$$

This ratio $\sigma^\alpha = \Delta\phi_1^\alpha/\Delta\phi_2^\alpha$ is essentially the susceptibility of component 1 to component 2 in the dilute phase $\alpha$. This expression appears to diverge in the limit $\phi_2 \to 0^+$ because $\hat{H}_{22}(\boldsymbol{\phi}^\alpha) \sim 1/\phi_2^\alpha \to \infty$; however, the product $\hat{H}_{22}(\boldsymbol{\phi}^\alpha)(\phi_2^\beta - \phi_2^\alpha)$ remains finite. To regularize this, we assume that the solute partition coefficient $k_2$ is constant throughout the phase space:

$$k_2 \equiv \frac{\phi_2^\beta}{\phi_2^\alpha}.$$

To obtain the solute partition coefficient $k_2$, we solve $\hat{\mu}_2(\boldsymbol{\phi}^\alpha) = \hat{\mu}_2(\boldsymbol{\phi}^\beta)$. The solute chemical potential $\hat{\mu}_2(\boldsymbol{\phi})$ is given by

$$\hat{\mu}_2(\boldsymbol{\phi}) = \frac{\partial f}{\partial \phi_2} = \log\phi_2 - \log(1-\phi_1-\phi_2) + (\chi_{12}-\chi_{13}-\chi_{23})\phi_1 - 2\chi_{23}\phi_2 + \chi_{23}.$$

Thus, $\hat{\mu}_2(\boldsymbol{\phi}^\alpha) = \hat{\mu}_2(\boldsymbol{\phi}^\beta)$ yields

$$k_2 = \frac{1-\phi_1^\alpha-\phi_2^\alpha}{1-\phi_1^\beta-\phi_2^\beta}\exp\left[(\chi_{12}-\chi_{13}-\chi_{23})(\phi_1^\beta-\phi_1^\alpha) - 2\chi_{23}(\phi_2^\beta-\phi_2^\alpha)\right]. \tag{67}$$

Expressing Eq. 67 using $k_2$ and the total volume fraction of component 2 $\phi_2$ makes it clear that this equation is transcendental about $k_2$:

$$k_2 = \frac{1-\phi_1^\alpha - c^\alpha(k_2)\phi_2}{1-\phi_1^\beta - c^\beta(k_2)\phi_2}\exp\left[(\chi_{12}-\chi_{13}-\chi_{23})(\phi_1^\beta-\phi_1^\alpha) - 2\chi_{23}\left[\, c^\beta(k_2) - c^\alpha(k_2)\,\right]\phi_2\right] \tag{68}$$

where $c^\alpha = 1/(1-\Phi+\Phi k_2)$ and $c^\beta = k_2/(1-\Phi+\Phi k_2)$. Here, $\Phi$ is the volume fraction of the phase $\beta$. In the limit $\phi_2 \to 0$, $k_2$ becomes

$$\lim_{\phi_2\to 0^+} k_2 = \frac{1-\phi_1^\alpha}{1-\phi_1^\beta}\exp\left[(\chi_{12}-\chi_{13}-\chi_{23})(\phi_1^\beta-\phi_1^\alpha)\right]. \tag{69}$$

We consider Eq. 66 in the limit $\phi_2 \to 0^+$ and assume a standard solvent. That is, $\tilde{v}_3 = v_0$ and $N_3 = 1$. Then, each term in the numerator and denominator is given by

$$\lim_{\phi_2\to 0^+} \hat{H}_{12}(\hat{\boldsymbol{\phi}}^\alpha)(\hat{\phi}_1^\beta - \hat{\phi}_1^\alpha) = \frac{k_BT}{v_0}\left(\frac{1}{1-\hat{\phi}_1^\alpha} + \chi_{12} - \chi_{13} - \chi_{23} + \chi_{33}\right)(\hat{\phi}_1^\beta - \hat{\phi}_1^\alpha)$$

$$\lim_{\hat{\phi}_2\to 0} \hat{H}_{22}(\hat{\boldsymbol{\phi}}^\alpha)(\hat{\phi}_2^\beta - \hat{\phi}_2^\alpha) = \frac{k_BT}{v_0}\frac{v_0}{\tilde{v}_2 N_2}\frac{k_2-1}{(1-\Phi)+\Phi k_2}$$

$$\lim_{\hat{\phi}_2\to 0} \hat{H}_{11}(\hat{\boldsymbol{\phi}}^\alpha)(\hat{\phi}_1^\beta - \hat{\phi}_1^\alpha) = \frac{k_BT}{v_0}\left(\frac{v_0}{\tilde{v}_1 N_1}\frac{1}{\hat{\phi}_1^\alpha} + \frac{1}{1-\hat{\phi}_1^\alpha} + \chi_{11} + \chi_{33} - 2\chi_{13}\right)(\hat{\phi}_1^\beta - \hat{\phi}_1^\alpha)$$

$$\lim_{\hat{\phi}_2\to 0} \hat{H}_{21}(\hat{\boldsymbol{\phi}}^\alpha)(\hat{\phi}_2^\beta - \hat{\phi}_2^\alpha) = 0.$$

Eq. 66 becomes

$$\lim_{\phi_2\to 0^+}\sigma^\alpha = \lim_{\phi_2\to 0^+}\frac{\Delta\phi_1^\alpha}{\Delta\phi_2^\alpha} = -\frac{\left[\frac{1}{1-\hat\phi_1^\alpha} + \chi_{12} - \chi_{13} - \chi_{23} + \chi_{33}\right](\hat\phi_1^\beta - \hat\phi_1^\alpha) + \frac{v_0}{\tilde v_2 N_2}\frac{k_2-1}{(1-\Phi)+\Phi k_2}}{\left(\frac{v_0}{\tilde v_1 N_1}\frac{1}{\hat\phi_1^\alpha} + \frac{1}{1-\hat\phi_1^\alpha} + \chi_{11} + \chi_{33} - 2\chi_{13}\right)(\hat\phi_1^\beta - \hat\phi_1^\alpha)}$$

$$= -\frac{\frac{1}{1-\hat\phi_1^\alpha} + (\chi_{12} - \chi_{13} - \chi_{23} + \chi_{33}) + \frac{k_2-1}{(1-\Phi)+\Phi k_2}\frac{v_0}{\tilde v_2 N_2}\frac{1}{\hat\phi_1^\beta - \hat\phi_1^\alpha}}{\frac{v_0}{\tilde v_1 N_1}\frac{1}{\hat\phi_1^\alpha} + \frac{1}{1-\hat\phi_1^\alpha} + (\chi_{11} + \chi_{33} - 2\chi_{13})}.$$

So far, the derivation is general. The only assumptions are (1) that the free-energy density follows the Flory–Huggins form, (2) that $k_2$ remains constant in phase space, and (3) that the solvent is simple ($\tilde v_3 = v_0$ and $N_3 = 1$).

Next, we adopt a two-body interaction form for the $\chi$ matrix, encoding enthalpic changes for heterotypic contacts as $\chi_{ij} = \epsilon_{ij} - \frac{1}{2}(\epsilon_{ii} + \epsilon_{jj})$ where $\epsilon_{ij}$ denotes the pairwise interaction strength. This leads to $\chi_{11} = \chi_{22} = \chi_{33} = 0$. It follows that

$$\lim_{\phi_2\to 0^+}\sigma^\alpha = \lim_{\phi_2\to 0^+}\frac{\Delta\phi_1^\alpha}{\Delta\phi_2^\alpha} = -\frac{1}{\Gamma}\frac{N_1\tilde v_1}{v_0}\hat\phi_1^\alpha\,(1+\chi^\Delta + h)$$

where

$$\begin{aligned}
\chi^\Delta &\equiv \chi_{12} - \chi_{13} - \chi_{23},\\
h &= \underbrace{\frac{k_2-1}{(1-\Phi)+\Phi k_2}}_{\text{solute partitioning}} \cdot \underbrace{\frac{v_0}{\tilde v_2 N_2}}_{\text{size dependence}} \cdot \underbrace{\frac{1}{\hat\phi_1^\beta - \hat\phi_1^\alpha}}_{\text{location dependence}},\\
&= \frac{k_2-1}{k_1-1}\,\frac{(1-\Phi)+\Phi k_1}{(1-\Phi)+\Phi k_2}\,\frac{v_0}{\tilde v_2 N_2}\,\frac{1}{\hat\phi_1},\\
\Gamma &= 1 + \frac{N_1\tilde v_1}{v_0}\hat\phi_1^\alpha(1-2\chi_{13}) + \mathcal{O}\left((\hat\phi_1^\alpha)^2\right).
\end{aligned}$$

Here, the scaffold partition coefficient $k_1 = \phi_1^\beta/\phi_1^\alpha$ is not fixed, and depends on the solute concentration $c_2$.

To convert $\sigma^\alpha$ to susceptibility $s_{12}$, we use the chain rule

$$\begin{aligned}
s_{12} &\equiv \left(\frac{\partial c_1^\alpha}{\partial c_2}\right)_{c_1} = \left(\frac{\partial c_1^\alpha}{\partial c_2^\alpha}\right)_{c_1}\left(\frac{\partial c_2^\alpha}{\partial c_2}\right)_{c_1}\\
&= \left(\frac{\partial c_1^\alpha}{\partial c_2^\alpha}\right)_{c_1}\left[(1-\Phi) + \Phi\left(\frac{\partial c_2^\beta}{\partial c_2^\alpha}\right)_{c_1} + \left(c_2^\beta - c_2^\alpha\right)\left(\frac{\partial \Phi}{\partial c_2^\alpha}\right)_{c_1}\right]^{-1}
\end{aligned}$$

since $c_2 = (1-\Phi)c_2^\alpha + \Phi c_2^\beta$. In the steady dense-phase assumption and for negligible changes in $\Phi$, this reduces to

$$s_{12} \approx \frac{1}{1-\Phi}\left(\frac{\partial c_1^\alpha}{\partial c_2^\alpha}\right)_{c_1}.$$

Noting that $(\partial c_1^\alpha/\partial c_2^\alpha)_{c_1} = (N_2\tilde v_2/N_1\tilde v_1)\cdot(\partial\phi_1^\alpha/\partial\phi_2^\alpha)_{\phi_1}$,

$$\begin{aligned}
s_{12} &\approx \frac{1}{1-\Phi}\,\frac{N_2\tilde v_2}{N_1\tilde v_1}\,\sigma^\alpha\\
&= -\frac{1}{1-\Phi}\,\frac{1}{\Gamma}\frac{v_2}{v_0}\,\hat\phi_1^\alpha\,(1+\chi^\Delta + h)
\end{aligned}$$

where $v_2 = N_2\tilde{v}_2$ denotes the molecular volume. Finally, we consider the dilute limit ($\phi_1 \ll 1$) and use the Taylor expansion $x/(1-ax) = x + \mathcal{O}(x^2)$ for $|ax| \ll 1$, leading to $\hat{\phi}_1^\alpha/\Gamma = \hat{\phi}_1^\alpha + \mathcal{O}\left((\hat{\phi}_1^\alpha)^2\right)$ Adopting the notation $s_{12} = s$ and $\hat{\phi}_1^\alpha = N_A v_1 c_1^{\text{dil}}(\mathbf{c})$, we obtain

$$s(\mathbf{c}) \approx -\frac{1}{1-\Phi}\left(\frac{v_2}{v_0}\right)\left(N_A v_1 c_1^{\text{dil}}(\mathbf{c})\right)(1+\chi^\Delta + h), \tag{70}$$

which is presented as Eq. 3 in the main text.

##### 3. *Summary*

We summarize below the expressions for the solute–scaffold susceptibility $s_{12}$ obtained from the naïve and perturbative Flory–Huggins models:

(Perturbative Flory–Huggins)

$$s(\mathbf{c}) \approx -\frac{1}{1-\Phi}\left(\frac{v_2}{v_0}\right)\left(N_A v_1 c_1^{\text{dil}}(\mathbf{c})\right)(1+\chi^\Delta + h)$$

where

$$\chi^\Delta = \chi_{12} - \chi_{13} - \chi_{23}.$$

$$h = \underbrace{\frac{k_2 - 1}{(1-\Phi)+\Phi k_2}}_{\text{solute partitioning}} \cdot \underbrace{\frac{v_0}{\tilde{v}_2 N_2}}_{\text{size dependence}} \cdot \underbrace{\frac{1}{\phi_1^{\text{den}} - \phi_1^{\text{dil}}}}_{\text{location-dependence}},$$

$$\approx \frac{k_2-1}{k-1}\frac{v_0}{v_2}\frac{1}{\phi_1} \quad \text{(Small dense-phase volume fraction: } \Phi \ll 1\text{)}.$$

Assumptions (perturbative FH)

$\phi_2 \to 0^+$ (Dilute solute limit)
$\phi_1 \ll 1$ (Small protein fraction in the dilute phase)
$\phi_1^{\text{dil}} = \hat{\phi}_1^{\text{dil}} + \sigma^{\text{dil}}\phi_2^{\text{dil}}$ (Linear response)
$k_2 = \phi_2^{\text{den}}/\phi_2^{\text{dil}} = \text{const.}$ (Constant solute partitioning)

##### F. Susceptibility to depletion interactions (crowding): BSA-PEG

In this section, we review how crowding agents adjust the onset of LLPS using the theory of hard-sphere fluids. Several theoretical frameworks describe depletion-driven phase transitions (see Section 3 in [21] and [22] for reviews). Conceptually, phase diagrams can be obtained by equating the chemical potentials and osmotic pressures of macromolecules and depletants, modeled as hard spheres. Depletants are often treated as penetrable hard spheres of radius $R_2$. A modern and quantitatively accurate formulation is the **free-volume theory (FVT)**, developed in the early 1990s [23, 24]. In FVT, the chemical potentials of depletants and macromolecules are computed directly from the (semi-)grand potential. The system is assumed to be in contact with a reservoir of depletants, meaning that the number of depletants is not fixed. Meanwhile, the number of macromolecules in the system is held constant. Under these conditions, the system is naturally described by the semi-grand potential.

For non-interacting depletants, the non-dimensionalized chemical potential and osmotic pressure of the macromolecules can be expressed as

$$\tilde{\mu}_1 = \frac{\mu_1}{k_B T} = \tilde{\mu}_1^\circ + \tilde{P}^R g(\phi_1), \tag{71}$$

$$\tilde{P} = \frac{P v_0}{k_B T} = \tilde{P}^\circ + \tilde{P}^R h(\phi_1), \tag{72}$$

where $g(\phi_1)$ and $h(\phi_1)$ describe the free-volume dependence ($V - V_{\text{excluded}}$) (See Section 3.3.4 in [21]). These functions are given by

$$\begin{aligned} g(\phi_1) &= e^{-Q(\phi_1)}[1 + (1+y)(a + 2by + 3cy^2)], \\ h(\phi_1) &= e^{-Q(\phi_1)}(1 + ay + 2by^2 + 3cy^3), \end{aligned}$$

where

$$\begin{aligned} Q &= ay + by^2 + cy^3, \\ a &= 3q + 3q^2 + q^3 = (1+q)^3 - 1, \\ b &= \tfrac{9}{2}q^2 + 3q^3, \\ c &= 3q^3, \\ y &= \frac{\phi_1}{1-\phi_1}. \end{aligned}$$

Here, $q = R_2/R_1$ is the size ratio between depletant and macromolecule.

The dimensionless pressure of the reservoir, $\tilde{P}^R$, follows from the coexistence conditions:

$$\tilde{P}^R = \frac{\tilde{\mu}^\circ(\phi_1) - \tilde{\mu}^\circ(\phi_1^{\text{dil}})}{g(\phi_1^{\text{dil}}) - g(\phi_1)} = \frac{\tilde{P}^\circ(\phi_1) - \tilde{P}^\circ(\phi_1^{\text{dil}})}{h(\phi_1^{\text{dil}}) - h(\phi_1)}. \tag{73}$$

Solving Eqs. 71–73 self-consistently yields the coexistence compositions, enabling prediction of phase diagrams. Figure S12a shows the phase diagrams predicted by free-volume theory for size ratios $q = 0.4$–$0.9$. As the depletant size becomes more comparable to that of the macromolecule, the two-phase region expands. In our BSA–PEG4k system, the radius of gyration of BSA is $3.0 \pm 0.1$ nm from X-ray scattering measurements, and that of PEG4k is 2.64 nm [11], giving a size ratio $q \simeq 0.88$. Accordingly, the blue curve in Fig. S12 is most relevant to our experiments.

The calculated phase diagrams allow computation of the binodal gradient $r$, susceptibility $s$, and tie-line slope $t$, shown in Fig. S12b, d, and f. For any value of $q$, the geometric relation

$$\frac{1}{r} = \frac{1}{s} + \frac{1}{t}$$

is satisfied. As discussed in Section IV B, in the absence of direct interactions between components 1 and 2, the susceptibility is expected to scale as

$$s \sim \left(\frac{v_2}{v_0}\right) \phi_1^{\mathrm{dil}} \sim q^3 \phi_1^{\mathrm{dil}}.$$

Figure S12e plots $s$ as a function of $q^3\phi_1^{\mathrm{dil}}$, for which this scaling is clearly observed in the dilute limit ($\phi_1^{\mathrm{dil}} \ll 1$). At higher volume fractions, the scaling no longer holds, with the crossover volume fraction depending on $q$. For the BSA/PEG4k system in this paper, the size ratio $q$ is approximately 0.9. For this case, free-volume theory predicts $s \propto (v_2/v_0)\phi_1^{\mathrm{dil}}$ for $\phi_1^{\mathrm{dil}} < 4 \times 10^{-2}$.

We note that these curves are not expected to collapse. Depletion interactions are not self-similar. For this reason, neither the binodal gradient $r$ nor the tie-line slope $t$ collapse when plotted against $(v_2/v_0)\, \phi_1^{\mathrm{dil}} \sim q^3\phi_1^{\mathrm{dil}}$.

###### 1. *Summary*

(Free-volume theory, crowding)

$$s \approx -A(q)\left(\frac{v_2}{v_0}\right) \phi_1^{\mathrm{dil}} \qquad \text{for } \phi_1^{\mathrm{dil}} < \phi_c(q)$$

where $q = R_2/R_1$. For $q = R_{\mathrm{PEG4k}}/R_{\mathrm{BSA}} = 0.88$, we numerically find $A(q = 0.88) = 5.0$ and $\phi_c = 4 \times 10^{-2}$.

Assumptions

Mixture of hard spheres (radius $R_1$) and penetrable hard spheres (radius $R_2$).
Phase coexistence for both macromolecules and depletant $\tilde{\mu}^{\mathrm{dil}} = \tilde{\mu}^{\mathrm{den}}$, $\tilde{P}^{\mathrm{dil}} = \tilde{P}^{\mathrm{den}}$.

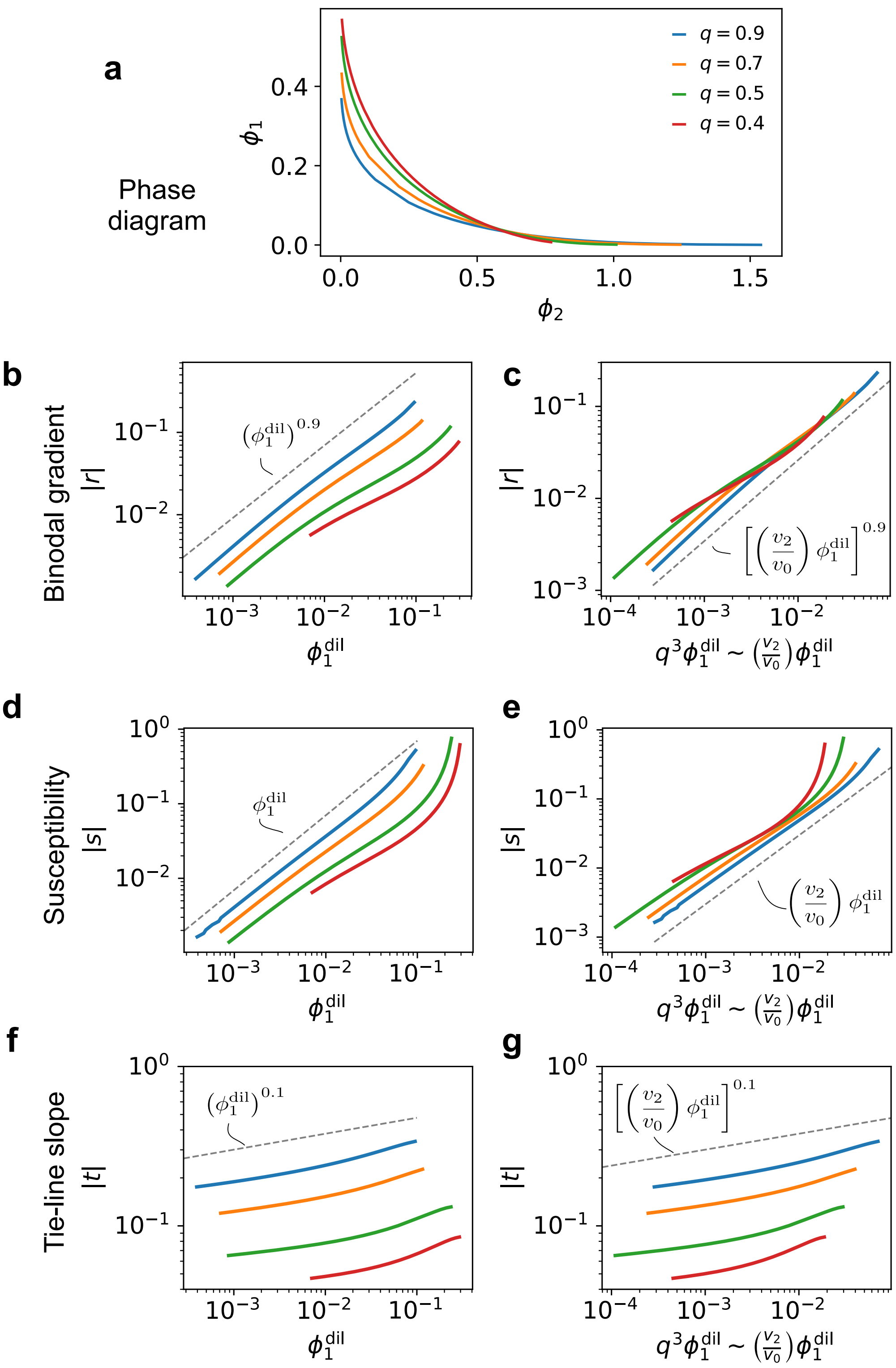


FIG. S12. **Free-volume–theory predictions of the binodal gradient, susceptibility, and tie-line slope.** **(a)** Binodal curves for macromolecule–depletant mixtures at size ratios $q = R_2/R_1$ ranging from 0.4 to 0.9. **(b)** Magnitude of binodal gradient $r \equiv dc_1/dc_2$ vs the dilute-phase volume fraction of component 1, $\phi_1^{\rm dil}$, for each $q$. **(c)** Same data as in (b) plotted against the scaled volume fraction $(v_2/v_0)\phi_1^{\rm dil}$. **(d)** Magnitude of susceptibility $s \equiv \left(\partial c_1^{\rm dil}/\partial c_2^{\rm tot}\right)_{c_1^{\rm tot}}$ vs $\phi_1^{\rm dil}$. **(e)** Same data as in (d) vs the scaled volume fraction. consistent with the dilute-limit scaling in Eq. 56. **(f)** Magnitude of tie-line slope $t \equiv (c_1^{\rm den} - c_1^{\rm dil})/(c_2^{\rm den} - c_2^{\rm dil})$ vs $\phi_1^{\rm dil}$. **(g)** Same data as in (d) vs the scaled volume fraction.

##### G. Susceptibility to ligand-pocket interactions: Bik1-Peptide

In this section, we derive the susceptibility of macromolecular condensates to solutes that bind specifically to scaffold molecules, such as N-acetylated EETF to Bik1. Such binding reactions violate the assumptions underlying Flory-Huggins theory, which treat protein-solute interactions in a mean-field manner. Instead, specific binding consumes the solute (ligand) $X$ and scaffold protein $M$ to form a biochemically distinct complex $MX$. The **polyphasic linkage theory** [25, 26] provides a more direct route by introducing measurable binding constants $K$ (or $K_D = 1/K$), linking microscopic chemical equilibria to macroscopic phase behavior.

###### 1. *Chemical equilibrium between coexisting liquid phases*

We consider chemical equilibrium between two coexisting liquid phases and derive the susceptibility to ligands. This minimal model describes the Bik1–peptide systems studied here, in which the dilute and dense phases are denoted by dil and den, respectively.

In each phase $\alpha \in \{\mathrm{dil}, \mathrm{den}\}$, we assume the binding reaction

$$\mathrm{M} + \mathrm{X} \rightleftharpoons \mathrm{MX},$$

with equilibrium constant

$$K^\alpha = \frac{a^\alpha_{\mathrm{MX}}}{a^\alpha_{\mathrm{M}} a^\alpha_{\mathrm{X}}},$$

where $a^\alpha_i = \gamma^\alpha_i c^\alpha_i / c_0$ is the activity of species $i$ in phase $\alpha$. Here, $\gamma^\alpha_i$ is the activity coefficient and $c_0$ is a reference concentration (e.g., 1 M).

We introduce the binding polynomial $P^\alpha$ as

$$P^\alpha(a^\alpha_{\mathrm{X}}) = 1 + K^\alpha a^\alpha_{\mathrm{X}},$$

which relates the *free* and *total* macromolecule activities in phase $\alpha$, $a^\alpha_{\mathrm{M}}$ and $a^\alpha_{\mathrm{MT}}$, via

$$a^\alpha_{\mathrm{MT}} = P^\alpha a^\alpha_{\mathrm{M}},$$

where $a^\alpha_{\mathrm{MT}} = a^\alpha_{\mathrm{M}} + a^\alpha_{\mathrm{MX}}$.

The chemical potential of the free macromolecule in phase $\alpha$ is given by

$$\begin{aligned} \mu^\alpha_{\mathrm{M}} &= \mu^\circ_{\mathrm{M}} + k_B T \ln a^\alpha_{\mathrm{M}} \\ &= \mu^\circ_{\mathrm{M}} + k_B T \ln a^\alpha_{\mathrm{MT}} - k_B T \ln P^\alpha. \end{aligned} \tag{74}$$

At coexistence between phases $\alpha$ and $\beta$, the free macromolecule chemical potentials are equal, $\mu^\alpha_{\mathrm{M}} = \mu^\beta_{\mathrm{M}}$. This yields

$$\begin{aligned} \mu^\circ_{\mathrm{M}} + k_B T \ln a^\alpha_{\mathrm{MT}} - k_B T \ln P^\alpha &= \mu^\circ_{\mathrm{M}} + k_B T \ln a^\beta_{\mathrm{MT}} - k_B T \ln P^\beta \\ \Leftrightarrow \quad a^\alpha_{\mathrm{MT}} &= \frac{P^\alpha}{P^\beta} a^\beta_{\mathrm{MT}} \\ \Leftrightarrow \quad c^\alpha_{\mathrm{MT}} &= \frac{P^\alpha}{P^\beta} \frac{\gamma^\beta_{\mathrm{MT}}}{\gamma^\alpha_{\mathrm{MT}}} c^\beta_{\mathrm{MT}}. \end{aligned} \tag{75}$$

Here, the activity coefficient of the total macromolecule is defined as follows:

$$\begin{aligned}\gamma_{\rm MT}^{\alpha} \equiv \frac{a_{\rm MT}^{\alpha}}{c_{\rm MT}^{\alpha}} &= \frac{a_{\rm M}^{\alpha} + a_{\rm MX}^{\alpha}}{c_{\rm M}^{\alpha} + c_{\rm MX}^{\alpha}} \\ &= \frac{\gamma_{\rm M}^{\alpha} c_{\rm M}^{\alpha} + \gamma_{\rm MX}^{\alpha} c_{\rm MX}^{\alpha}}{c_{\rm M}^{\alpha} + c_{\rm MX}^{\alpha}} \\ &= \frac{1}{P^{\alpha}(a_{\rm X}^{\alpha})}\gamma_{\rm M}^{\alpha} + \left(1 - \frac{1}{P^{\alpha}(a_{\rm X}^{\alpha})}\right)\gamma_{\rm MX}^{\alpha}\end{aligned} \tag{76}$$

##### 2. *Definition of susceptibility for strongly binding solutes*

We must first define the susceptibility to the strongly binding solutes (ligands). Because experiments measure the *total* macromolecule concentration rather than separately resolving free (M) and ligated species (MX), we define the susceptibility to binding solutes as the change in the *total* dilute-phase macromolecule concentration induced by adding a unit amount of ligand, at fixed total macromolecule concentration $c_{\rm MT}$:

$$s \equiv \left(\frac{\partial c_{\rm MT}^{\rm dil}}{\partial c_{\rm XT}}\right)_{c_{\rm MT}} . \tag{77}$$

Using the chain rule, Eq. 77 can be written as

$$s = \left(\frac{\partial c_{\rm MT}^{\rm dil}}{\partial a_{\rm X}^{\rm dil}}\right)_{c_{\rm MT}} \Bigg/ \left(\frac{\partial c_{\rm XT}}{\partial a_{\rm X}^{\rm dil}}\right)_{c_{\rm MT}} . \tag{78}$$

This is because that $c_{\rm MT}^{\rm dil}$ is a function of only a single variable. To see this, we perform a bookkeeping of the degrees of freedom relevant to $s$. For this bookkeeping, we count using concentrations instead of activities but this choice does not alter the degree of freedom in the system. There are 7 state variables:

$$c_{\rm M}^{\rm dil}, c_{\rm X}^{\rm dil}, c_{\rm MX}^{\rm dil}, c_{\rm M}^{\rm den}, c_{\rm X}^{\rm den}, c_{\rm MX}^{\rm den}, \Phi.$$

The quantities $K$, $c_{\rm MT}$, and $c_{\rm XT}$ are treated as fixed parameters; $K$ depends only on thermodynamic state variables such as pressure and temperature. By contrast, there are 6 independent constraints:

1. Equilibrium in the dilute phase: $c_{\rm MX}^{\rm dil} = K\, c_{\rm M}^{\rm dil} c_{\rm X}^{\rm dil}$.
2. Equilibrium in the dense phase:$c_{\rm MX}^{\rm den} = K\, c_{\rm M}^{\rm den} c_{\rm X}^{\rm den}$.
3. Mass balance for macromolecule $M$: $c_{\rm MT} = \Phi\,(c_{\rm M}^{\rm den} + c_{\rm MX}^{\rm den}) + (1-\Phi)\,(c_{\rm M}^{\rm dil} + c_{\rm MX}^{\rm dil})$.
4. Mass balance for solute $X$: $c_{\rm XT} = \Phi\,(c_{\rm X}^{\rm den} + c_{\rm MX}^{\rm den}) + (1-\Phi)\,(c_{\rm X}^{\rm dil} + c_{\rm MX}^{\rm dil})$.
5. Equality of macromolecule $M$ chemical potentials: $\mu_{\rm M}^{\rm dil} = \mu_{\rm M}^{\rm den}$.
6. Equality of ligand $X$ chemical potentials: $\mu_{\rm X}^{\rm dil} = \mu_{\rm X}^{\rm den}$.

The system therefore has 7 state variables and 6 independent constraints, leaving 1 internal degree of freedom. As a result, all state variables are functions of a single parameter. A convenient choice is the dilute-phase ligand activity, meaning $c_{\rm MT}^{\rm dil} = c_{\rm MT}^{\rm dil}(a_{\rm X}^{\rm dil})$ and $c_{\rm XT} = c_{\rm XT}(a_{\rm X}^{\rm dil})$. With this choice, Eq. 78 follows directly from the chain rule.

*3. Derivation of Eq. 4 in the main text*

In this section, we derive Eq. 4 in the main text from Eq. 78. Starting with its numerator, Eq. 75 can be rewritten as

$$c_{\mathrm{MT}}^{\mathrm{dil}} = \frac{1}{\gamma_{\mathrm{MT}}^{\mathrm{dil}}} A(a_{\mathrm{MT}}^{\mathrm{den}}, a_{\mathrm{X}}^{\mathrm{den}}) \; P^{\mathrm{dil}}(a_{\mathrm{X}}^{\mathrm{dil}})$$

where

$$A(a_{\mathrm{MT}}^{\mathrm{den}}, a_{\mathrm{X}}^{\mathrm{den}}) \equiv \frac{\gamma_{\mathrm{MT}}^{\mathrm{den}} \, c_{\mathrm{MT}}^{\mathrm{den}}}{P^{\mathrm{den}}(a_{\mathrm{X}}^{\mathrm{den}})} = c_0 \frac{a_{\mathrm{MT}}^{\mathrm{den}}}{P^{\mathrm{den}}(a_{\mathrm{X}}^{\mathrm{den}})}$$

Experiments indicate that the dense-phase composition is much less sensitive to ligand addition than the dilute-phase composition, so we treat $A(a_{\mathrm{MT}}^{\mathrm{den}}, a_{\mathrm{X}}^{\mathrm{den}})$ as approximately constant. Under this stationary dense-phase assumption, we obtain

$$\begin{aligned}
\left(\frac{\partial c_{\mathrm{MT}}^{\mathrm{dil}}}{\partial a_{\mathrm{X}}^{\mathrm{dil}}}\right)_{c_{\mathrm{MT}}} &\approx \frac{A}{\gamma_{\mathrm{MT}}^{\mathrm{dil}}} \left(\frac{dP^{\mathrm{dil}}(a_{\mathrm{X}}^{\mathrm{dil}})}{da_{\mathrm{X}}^{\mathrm{dil}}}\right) \\
&= \frac{A}{\gamma_{\mathrm{MT}}^{\mathrm{dil}}} K^{\mathrm{dil}} \qquad \because P^{\mathrm{dil}}(a_{\mathrm{X}}^{\mathrm{dil}}) = 1 + K^{\mathrm{dil}} a_{\mathrm{X}}^{\mathrm{dil}} \\
&= \frac{c_{\mathrm{MT}}^{\mathrm{dil}}}{P^{\mathrm{dil}}(a_{\mathrm{X}}^{\mathrm{dil}})} K^{\mathrm{dil}} \qquad \because c_{\mathrm{MT}}^{\mathrm{dil}} = A P^{\mathrm{dil}}(a_{\mathrm{X}}^{\mathrm{dil}})/\gamma_{\mathrm{MT}}^{\mathrm{dil}} \\
&= \frac{c_0}{\gamma_{\mathrm{MT}}^{\mathrm{dil}}} \frac{K^{\mathrm{dil}} a_{\mathrm{MT}}^{\mathrm{dil}}}{P^{\mathrm{dil}}(a_{\mathrm{X}}^{\mathrm{dil}})}.
\end{aligned} \tag{79}$$

We next evaluate the denominator of Eq. 78, $\left(\partial c_{\mathrm{XT}}/\partial a_{\mathrm{X}}^{\mathrm{dil}}\right)_{c_{\mathrm{MT}}}$. The total concentration of the ligands is given by

$$c_{\mathrm{XT}} = (1-\Phi) c_{\mathrm{XT}}^{\mathrm{dil}} + \Phi c_{\mathrm{XT}}^{\mathrm{den}}$$

where $\Phi$ is the volume fraction of the dense phase. Under the stationary dense-phase assumption ($c_{\mathrm{XT}}^{\mathrm{den}} = \mathrm{const.}$),

$$dc_{\mathrm{XT}} \approx (1-\Phi)\, dc_{\mathrm{XT}}^{\mathrm{dil}}.$$

and therefore

$$\left(\frac{\partial c_{\mathrm{XT}}}{\partial a_{\mathrm{X}}^{\mathrm{dil}}}\right)_{c_{\mathrm{MT}}} \approx (1-\Phi) \left(\frac{\partial c_{\mathrm{XT}}^{\mathrm{dil}}}{\partial a_{\mathrm{X}}^{\mathrm{dil}}}\right)_{c_{\mathrm{MT}}}.$$

Using the chain rule,

$$\left(\frac{\partial c_{\mathrm{XT}}^{\mathrm{dil}}}{\partial a_{\mathrm{X}}^{\mathrm{dil}}}\right)_{c_{\mathrm{MT}}} = \left(\frac{\partial c_{\mathrm{XT}}^{\mathrm{dil}}}{\partial a_{\mathrm{XT}}^{\mathrm{dil}}}\right)_{c_{\mathrm{MT}}} \left(\frac{\partial a_{\mathrm{XT}}^{\mathrm{dil}}}{\partial a_{\mathrm{X}}^{\mathrm{dil}}}\right)_{c_{\mathrm{MT}}}. \tag{80}$$

From the definition of activity, $a_{\mathrm{XT}}^{\mathrm{dil}} = \gamma_{\mathrm{XT}}^{\mathrm{dil}} c_{\mathrm{XT}}^{\mathrm{dil}}/c_0$, the first factor is

$$\left(\frac{\partial c_{\mathrm{XT}}^{\mathrm{dil}}}{\partial a_{\mathrm{XT}}^{\mathrm{dil}}}\right)_{c_{\mathrm{MT}}} = \frac{c_0}{\gamma_{\mathrm{XT}}^{\mathrm{dil}}}.$$

For the second factor, note that

$$\begin{aligned}
a_{\mathrm{XT}}^{\mathrm{dil}} &= a_{\mathrm{X}}^{\mathrm{dil}} + a_{\mathrm{MX}}^{\mathrm{dil}} \\
&= a_{\mathrm{X}}^{\mathrm{dil}} + (a_{\mathrm{MT}}^{\mathrm{dil}} - a_{\mathrm{M}}^{\mathrm{dil}}) \\
&= a_{\mathrm{X}}^{\mathrm{dil}} + a_{\mathrm{MT}}^{\mathrm{dil}} \left(1 - \frac{1}{P^{\mathrm{dil}}(a_{\mathrm{X}}^{\mathrm{dil}})}\right).
\end{aligned} \tag{81}$$

Differentiating Eq. 81 with respect to $a_{\mathrm{X}}^{\mathrm{dil}}$ yields

$$\left(\frac{\partial a_{\mathrm{XT}}^{\mathrm{dil}}}{\partial a_{\mathrm{X}}^{\mathrm{dil}}}\right)_{c_{\mathrm{MT}}} = 1 + \left(\frac{\partial a_{\mathrm{MT}}^{\mathrm{dil}}}{\partial a_{\mathrm{X}}^{\mathrm{dil}}}\right)_{c_{\mathrm{MT}}} \left(1 - \frac{1}{P^{\mathrm{dil}}}\right) + \frac{K^{\mathrm{dil}} a_{\mathrm{MT}}^{\mathrm{dil}}}{(P^{\mathrm{dil}})^2}. \tag{82}$$

To compute $\left(\frac{\partial a_{\mathrm{MT}}^{\mathrm{dil}}}{\partial a_{\mathrm{X}}^{\mathrm{dil}}}\right)_{c_{\mathrm{MT}}}$, we note that the activity of the free macromolecule must be equal in the dilute and dense phases at phase coexistence:

$$a_{\mathrm{M}}^{\mathrm{dil}} = a_{\mathrm{M}}^{\mathrm{den}} \qquad \left(\because \mu_{\mathrm{M}}^{\mathrm{dil}} = \mu_{\mathrm{M}}^{\mathrm{den}}\right).$$

Using $a_{\mathrm{MT}}^{\alpha} = P^{\alpha} a_{\mathrm{M}}^{\alpha}$, this gives

$$a_{\mathrm{MT}}^{\mathrm{dil}} = \frac{P^{\mathrm{dil}}(a_{\mathrm{X}}^{\mathrm{dil}})}{P^{\mathrm{den}}(a_{\mathrm{X}}^{\mathrm{den}})}\, a_{\mathrm{MT}}^{\mathrm{den}}. \tag{83}$$

Under the stationary dense-phase assumption, $a_{\mathrm{MT}}^{\mathrm{den}}$ and $P^{\mathrm{den}}(a_{\mathrm{X}}^{\mathrm{den}})$ are constant. Differentiating Eq. 83 with respect to $a_{\mathrm{X}}^{\mathrm{dil}}$ yields

$$\begin{aligned}
\left(\frac{\partial a_{\mathrm{MT}}^{\mathrm{dil}}}{\partial a_{\mathrm{X}}^{\mathrm{dil}}}\right)_{c_{\mathrm{MT}}} &\approx \frac{a_{\mathrm{MT}}^{\mathrm{den}}}{P^{\mathrm{den}}(a_{\mathrm{X}}^{\mathrm{den}})}\, \frac{dP^{\mathrm{dil}}(a_{\mathrm{X}}^{\mathrm{dil}})}{da_{\mathrm{X}}^{\mathrm{dil}}} \\
&= \frac{K^{\mathrm{dil}} a_{\mathrm{MT}}^{\mathrm{den}}}{P^{\mathrm{den}}(a_{\mathrm{X}}^{\mathrm{den}})} \qquad \because P^{\mathrm{dil}} = 1 + K^{\mathrm{dil}} a_{\mathrm{X}}^{\mathrm{dil}} \\
&= \frac{K^{\mathrm{dil}} a_{\mathrm{MT}}^{\mathrm{dil}}}{P^{\mathrm{dil}}(a_{\mathrm{X}}^{\mathrm{dil}})} \qquad \because \text{Eq. } 83
\end{aligned} \tag{84}$$

Substituting Eq. 84 into Eq. 82 gives

$$\begin{aligned}
\left(\frac{\partial a_{\mathrm{XT}}^{\mathrm{dil}}}{\partial a_{\mathrm{X}}^{\mathrm{dil}}}\right)_{c_{\mathrm{MT}}} &= 1 + \frac{K^{\mathrm{dil}} a_{\mathrm{MT}}^{\mathrm{dil}}}{P^{\mathrm{dil}}(a_{\mathrm{X}}^{\mathrm{dil}})} \left(1 - \frac{1}{P^{\mathrm{dil}}}\right) + \frac{K^{\mathrm{dil}} a_{\mathrm{MT}}^{\mathrm{dil}}}{(P^{\mathrm{dil}})^2} \\
&= 1 + \frac{K^{\mathrm{dil}} a_{\mathrm{MT}}^{\mathrm{dil}}}{P^{\mathrm{dil}}(a_{\mathrm{X}}^{\mathrm{dil}})}.
\end{aligned} \tag{85}$$

The denominator of Eq. 78 becomes

$$\begin{aligned}
\left(\frac{\partial c_{\mathrm{XT}}}{\partial a_{\mathrm{X}}^{\mathrm{dil}}}\right)_{c_{\mathrm{MT}}} &\approx (1 - \Phi) \left(\frac{\partial c_{\mathrm{XT}}^{\mathrm{dil}}}{\partial a_{\mathrm{X}}^{\mathrm{dil}}}\right)_{c_{\mathrm{MT}}} \\
&= (1 - \Phi) \left(\frac{\partial c_{\mathrm{XT}}^{\mathrm{dil}}}{\partial a_{\mathrm{XT}}^{\mathrm{dil}}}\right)_{c_{\mathrm{MT}}} \left(\frac{\partial a_{\mathrm{XT}}^{\mathrm{dil}}}{\partial a_{\mathrm{X}}^{\mathrm{dil}}}\right)_{c_{\mathrm{MT}}} \\
&= (1 - \Phi) \cdot \frac{c_0}{\gamma_{\mathrm{XT}}^{\mathrm{dil}}} \cdot \left(1 + \frac{K^{\mathrm{dil}} a_{\mathrm{MT}}^{\mathrm{dil}}}{P^{\mathrm{dil}}(a_{\mathrm{X}}^{\mathrm{dil}})}\right).
\end{aligned} \tag{86}$$

Finally, substituting Eqs. 79 and 86 into Eq. 78, we obtain the susceptibility $s$:

$$\boxed{s \approx \frac{\gamma_{\mathrm{XT}}^{\mathrm{dil}}}{\gamma_{\mathrm{MT}}^{\mathrm{dil}}} \cdot \frac{1}{1-\Phi} \cdot \frac{K^{\mathrm{dil}} a_{\mathrm{MT}}^{\mathrm{dil}}}{P^{\mathrm{dil}}(a_{\mathrm{X}}^{\mathrm{dil}}) + K^{\mathrm{dil}} a_{\mathrm{MT}}^{\mathrm{dil}}}.} \tag{87}$$

In the low-ligand limit $a_{\mathrm{X}}^{\mathrm{dil}} \to 0$, the binding polynomial satisfies $P^{\mathrm{dil}}(a_{\mathrm{X}}^{\mathrm{dil}}) \to 1$. If, in addition, the dilute-phase ligand activity coefficient is close to unity ($\gamma_{\mathrm{XT}}^{\mathrm{dil}} \simeq 1$), the susceptibility reduces to

$$s \approx \frac{1}{\gamma_{\mathrm{MT}}^{\mathrm{dil}}(1-\Phi)} \frac{K^{\mathrm{dil}} a_{\mathrm{MT}}^{\mathrm{dil}}}{1 + K^{\mathrm{dil}} a_{\mathrm{MT}}^{\mathrm{dil}}}. \qquad \text{(low ligand limit)} \tag{88}$$

In practice, activities of free and ligated macromolecules in the dilute phase are unknown, and the concentration-based equilibrium constant $K \equiv c_{\mathrm{MX}}^{\mathrm{dil}}/(c_{\mathrm{M}}^{\mathrm{dil}} c_{\mathrm{X}}^{\mathrm{dil}}) = K^{\mathrm{dil}}/c_0$ is more accessible. We therefore compare our experimental data to the following form:

$$\boxed{s \approx \frac{1}{1-\Phi} \frac{K c_{\mathrm{MT}}^{\mathrm{dil}}}{1 + K c_{\mathrm{MT}}^{\mathrm{dil}}} = \frac{1}{1-\Phi} \frac{c_{\mathrm{MT}}^{\mathrm{dil}}}{K_D + c_{\mathrm{MT}}^{\mathrm{dil}}} \qquad (\gamma_{\mathrm{MT}}^{\mathrm{dil}} = \gamma_{\mathrm{M}}^{\mathrm{dil}} = \gamma_{\mathrm{MX}}^{\mathrm{dil}} = 1),} \tag{89}$$

where $K_D \equiv 1/K$ is the binding affinity. This expression is presented as Eq. 4 in the main text. In the strong-binding limit ($K \to \infty$), $s \to 1/(1-\Phi)$, which is approximately one, consistent with the observed susceptibility of binding peptides to Bik1.

It is also useful to define a characteristic ligand concentration,

$$c^* \equiv \frac{c_{\mathrm{MT}}^{\mathrm{dil}}}{s} = (1-\Phi)(K_D + c_{\mathrm{MT}}^{\mathrm{dil}}), \tag{90}$$

which shows that the ligand concentration required to sufficiently perturb phase separation scales with the sum of the binding affinity $K_D$ and the dilute-phase macromolecule concentration $c_{\mathrm{MT}}^{\mathrm{dil}}$. In the strong-binding limit ($K_D \to 0$), $c^*$ approaches $(1-\Phi)c_{\mathrm{MT}}^{\mathrm{dil}}$.

As a final remark, this analysis can be extended to systems with multiple binding sites of distinct affinities, where the binding polynomial generalizes to

$$P(a_{\mathrm{X}}) = 1 + K_1 a_{\mathrm{X}} + K_1 K_2 a_{\mathrm{X}}^2 + \cdots ,$$

and the same framework applies with $K$ replaced by an effective multivalent binding term. The binding polynomial relates the free and total macromolecule concentrations in each phase, and such generalization requires only minor modifications of Eqs. 79 and 86.

*4. Summary*

We summarize below the expressions for the lignad–scaffold susceptibility from the polyphasic linkage theory.

(Polyphasic linkage, single binding site)

$$s \approx \frac{\gamma^{\rm dil}_{\rm XT}}{\gamma^{\rm dil}_{\rm MT}} \cdot \frac{1}{1-\Phi} \cdot \frac{K^{\rm dil} a^{\rm dil}_{\rm MT}}{P^{\rm dil}(a^{\rm dil}_{\rm X}) + K^{\rm dil} a^{\rm dil}_{\rm MT}}$$

(Low-ligand limit ($P^{\rm dil}(a^{\rm dil}_{\rm X}) \to 1, \gamma^{\rm dil}_{\rm XT} \to 1$) and $\gamma_{\rm MT} = 1$)

$$s \;\to\; \frac{1}{1-\Phi}\frac{K\, c^{\rm dil}_{\rm MT}}{1 + K\, c^{\rm dil}_{\rm MT}} = \frac{1}{1-\Phi}\frac{c^{\rm den}_{\rm MT}}{K_D + c^{\rm den}_{\rm MT}}$$

Keys

- $\Phi$: volume fraction of the dense phase
- $K^{\rm dil} = a^{\rm dil}_{\rm MX}/(a^{\rm dil}_{\rm M} a^{\rm dil}_{\rm X})$: activity-based equilibrium constant in the dilute phase
- $K = c^{\rm dil}_{\rm MX}/(c^{\rm dil}_{\rm M} c^{\rm dil}_{\rm X})$: concentration-baed equilibrium constant in the dilute phase
- $a^\alpha_i$: activity of species $i$ in phase $\alpha$; $a^\alpha_i = \gamma^\alpha_i c^\alpha_i / c_0$
  - MT: Total macromolecules (free + bound)
  - XT: Total ligands (free + bound)
  - M: Free macromolecules
  - X: Free ligands
- $c^\alpha_i$: concentration of species $i$ in phase $\alpha$

Assumptions

- $a^{\rm den}_{\rm MT}$, $a^{\rm den}_{\rm X}$ = const. (stationary dense-phase activity)

#### H. Summary of susceptibility models

Predictions of the susceptibility are summarized in Table S6 with the consistent notations across the models. Depending on the scaffold–solute interaction, our experiment shows that the dilute-phase susceptibility spans a wide range between $|s| = 10^{-5}$–$10^{0}$ when the dilute-phase protein concentration $c_1^{\mathrm{dil}} = \mathcal{O}(10^1)$ $\mu$M and solute volumes $v_2 = 0.1 - 1$ nm$^3$.

TABLE S6. **Predicted susceptibility forms for distinct scaffold–solute interactions.**

| Type of Scaffold–Solute Interaction | Functional Form | Typical Magnitude |
|---|---|---|
| Crowding | $s \approx -A(q) \cdot N_A v_2 c_1^{\mathrm{dil}}(\mathbf{c})$ for $\phi_1^{\mathrm{dil}} = N_A v_1 c_1^{\mathrm{dil}}(\mathbf{c}) < \phi_c(q)$ <br> For BSA/PEG, $q = 0.88; A = 5, \phi_c = 0.04$ | $\mathcal{O}(10^{-2} \sim 10^{-1})$ |
| Promiscuous Interaction | $s \approx -\frac{1}{1-\Phi}\left(\frac{v_2}{v_0}\right)\left(N_A v_1 c_1^{\mathrm{dil}}(\mathbf{c})\right) \cdot \left(1 + \chi^{\Delta} + h\right)$ | $\mathcal{O}(10^{-5} \sim 10^{-3})$ |
| Specific binding | $s \approx \frac{1}{\gamma_{\mathrm{MT}}} \frac{1}{1-\Phi} \frac{c_1^{\mathrm{dil}}}{K_D + c_1^{\mathrm{dil}}}$ | $\mathcal{O}(10^{-1} \sim 1)$ |

Keys: $q$ is the size ratio between solute and scaffold, $q \equiv R_2/R_1$; $\chi^{\Delta} \equiv \chi_{12} - \chi_{13} - \chi_{23}$; $h \equiv \frac{k_2-1}{k-1}\frac{v_0}{v_2}\frac{1}{\phi_1}$; $K_D$ is the dissociation constant of the binding reaction $M + X \rightleftharpoons MX$; $N_A$ is Avogadro's number; $c_i^{\alpha}$ is the concentration of component $i$ in phase $\alpha$; $\phi_i^{\alpha}$ is the volume fraction of component $i$ in phase $\alpha$; $k$ is the protein partition coefficient; $k_2$ is the solute partition coefficient; $v_i$ is the molecular volume of component $i$; $v_0$ is the reference volume.

### V. CAPTIONS FOR SUPPLEMENTARY MOVIES

**Movie S1.**

Time-lapse confocal microscopy of Bik1 droplets dissolving in a gradient chamber. A peptide concentration gradient (N-acetylated EETF) is created by placing two hydrogels pre-equilibrated with solutions of different peptide concentrations ($\Delta c = 325$ $\mu$M). At later times, the local peptide concentration increases, causing Bik1 droplets to swell and ultimately dissolve. Bik1 is fluorescently labeled. The movie is played at 0.01× real time (100× slower than real time). The solution contains 366 $\mu$M Bik1 monomer equivalent, 20 mM Tris, 400 mM NaCl, pH 7.5.

**Movie S2.**

Induction of Bik1 condensation from a homogeneous phase via ATP hydrolysis using apyrase, imaged by confocal microscopy. The time-lapse shows that ATP consumption and concomitant production of ADP, AMP, and phosphate trigger condensation of Bik1. Bik1 is fluorescently labeled. The initial solution contains 366 $\mu$M Bik1 monomer equivalent, 20 mM ATP, 10 $\mu$M apyrase, 20 mM Tris, 350 mM NaCl, pH 6.8.

**Appendix A: Relation between the hessians of Helmholtz and Gibbs free energies**

Here, we derive the relation between the hessian of the Helmholtz free energy (fixed $V$ and $T$) and the hessian of the Gibbs free energy (fixed $P$ and $T$). The Gibbs free energy is obtained by the Legendre transform of the Helmholtz free energy $F$,

$$G(T, P, \{N_i\}) = F(T, V, \{N_i\}) + PV,$$

where $V$ is a function of $T$, $P$, and $\{N_i\}$:

$$V = V(T, P, \{N_i\}).$$

We first differentiate $G$ with respect to $N_i$ at fixed $T$, $P$, and $N_{\ell \neq i}$:

$$\begin{aligned}\left(\frac{\partial G}{\partial N_i}\right)_{T,P} &= \left(\frac{\partial F}{\partial N_i}\right)_{T,V} + \left(\frac{\partial F}{\partial V}\right)_{T,\{N_i\}} \left(\frac{\partial V}{\partial N_i}\right)_{T,P} + P\left(\frac{\partial V}{\partial N_i}\right)_{T,P} \\ &= \left(\frac{\partial F}{\partial N_i}\right)_{T,V} + \left[\left(\frac{\partial F}{\partial V}\right)_{T,\{N_i\}} + P\right] \left(\frac{\partial V}{\partial N_i}\right)_{T,P}.\end{aligned}$$

The term in brackets vanishes because of the thermodynamic relation $P = -\left(\frac{\partial F}{\partial V}\right)_{T,\{N_i\}}$.
Therefore,

$$\left(\frac{\partial G}{\partial N_i}\right)_{T,P} = \left(\frac{\partial F}{\partial N_i}\right)_{T,V} = \mu_i.$$

Taking another derivative with respect to $N_j$ at fixed $T$ and $P$ yields

$$\left(\frac{\partial^2 G}{\partial N_i \partial N_j}\right)_{T,P} = \left(\frac{\partial \mu_i}{\partial N_j}\right)_{T,P}.$$

Since

$$\mu_i = \left(\frac{\partial F}{\partial N_i}\right)_{T,V},$$

the derivative at fixed pressure contains both the direct dependence on $N_j$ and the indirect dependence through $V(T, P, \{N_i\})$:

$$\left(\frac{\partial \mu_i}{\partial N_j}\right)_{T,P,N_{i\neq j}} = \left(\frac{\partial \mu_i}{\partial N_j}\right)_{T,V,N_{i\neq j}} + \left(\frac{\partial \mu_i}{\partial V}\right)_{T,\{N_k\}} \left(\frac{\partial V}{\partial N_j}\right)_{T,P,N_{i\neq j}}.$$

It remains to evaluate $(\partial V/\partial N_j)_{T,P}$. Differentiating $P = -\left(\frac{\partial F}{\partial V}\right)_{T,\{N_i\}}$, with respect to $N_j$ at fixed $T$ and $P$ gives

$$0 = -\left(\frac{\partial^2 F}{\partial V \partial N_j}\right)_{T,V} - \left(\frac{\partial^2 F}{\partial V^2}\right)_{T,\{N_i\}} \left(\frac{\partial V}{\partial N_j}\right)_{T,P}$$

since $P$ is fixed. Thus,

$$\left(\frac{\partial V}{\partial N_j}\right)_{T,P} = -\frac{\left(\frac{\partial^2 F}{\partial V \partial N_j}\right)_{T,V}}{\left(\frac{\partial^2 F}{\partial V^2}\right)_{T,\{N_i\}}}.$$

Substituting this result gives

$$\boxed{\left(\frac{\partial^2 G}{\partial N_i \partial N_j}\right)_{T,P,N_{i\neq j}} = \left(\frac{\partial^2 F}{\partial N_i \partial N_j}\right)_{T,V,N_{i\neq j}} - \frac{\left(\frac{\partial^2 F}{\partial N_i \partial V}\right)_{T,\{N_k\}} \left(\frac{\partial^2 F}{\partial V \partial N_j}\right)_{T,V,N_{i\neq j}}}{\left(\frac{\partial^2 F}{\partial V^2}\right)_{T,\{N_k\}}}.} \tag{A1}$$

This is the constant-pressure correction to the fixed-volume hessian.

We now rewrite Eq. A1 in terms of concentration hessians. Let

$$F(T, V, \{N_i\}) = V f(T, \mathbf{c}), \qquad c_i = \frac{N_i}{V}.$$

We define the fixed-$V, T$ concentration hessian as

$$H_{ij} \equiv V \left(\frac{\partial^2 F}{\partial N_i \partial N_j}\right)_{T,V,N_{i\neq j}} = \left(\frac{\partial \mu_i}{\partial c_j}\right)_{T,V,\{c_{i\neq j}\}},$$

and the fixed-$P, T$ concentration hessian as

$$\mathcal{H}_{ij} \equiv V \left(\frac{\partial^2 G}{\partial N_i \partial N_j}\right)_{T,P,N_{i\neq j}} = \left(\frac{\partial \mu_i}{\partial c_j}\right)_{T,P,\{c_{i\neq j}\}},.$$

The mixed derivative involving $V$ follows from the dependence of concentration on volume at fixed particle number:

$$\left(\frac{\partial c_k}{\partial V}\right)_{T,\{N_k\}} = -\frac{N_k}{V^2} = -\frac{c_k}{V}.$$

Thus,

$$\left(\frac{\partial^2 F}{\partial N_i \partial V}\right)_{T,\{N_k\}} = \left(\frac{\partial \mu_i}{\partial V}\right)_{T,\{N_k\}} = -\frac{1}{V} \sum_k H_{ik} c_k,$$

and

$$\left(\frac{\partial^2 F}{\partial V \partial N_j}\right)_{T,V,N_{i\neq j}} = -\frac{1}{V} \sum_k H_{jk} c_k.$$

The volume curvature is

$$\left(\frac{\partial^2 F}{\partial V^2}\right)_{T,\{N_k\}} = \frac{1}{V} \sum_{kl} c_k H_{kl} c_l.$$

Substitution into Eq. A1 gives

$$\boxed{\mathcal{H}_{ij} = H_{ij} - \frac{\left(\sum_k H_{ik} c_k\right)\left(\sum_l H_{jl} c_l\right)}{\sum_{kl} c_k H_{kl} c_l}.} \tag{A2}$$

Equivalently,

$$\boxed{\mathcal{H} = H - \frac{(H\mathbf{c})(H\mathbf{c})^T}{\mathbf{c}^T H \mathbf{c}}.} \tag{A3}$$

[1] K. Lau, B. Bouchri, and F. Pojer, Purification of 10xHis-SuperTEV v1 (2023).

[2] J. R. Rumble, ed., *CRC handbook of chemistry and physics*, 102nd ed. (CRC Press, Boca Raton London New York, 2021).

[3] E. Gasteiger, C. Hoogland, A. Gattiker, S. Duvaud, M. R. Wilkins, R. D. Appel, and A. Bairoch, Protein identification and analysis tools on the expasy server, in *The Proteomics Protocols Handbook*, edited by J. M. Walker (Humana Press, Totowa, NJ, 2005) pp. 571–607.

[4] S. M. Meier, A.-M. Farcas, A. Kumar, M. Ijavi, R. T. Bill, J. Stelling, E. R. Dufresne, M. O. Steinmetz, and Y. Barral, Multivalency ensures persistence of a +TIP body at specialized microtubule ends, Nature Cell Biology **25**, 56 (2023).

[5] S. R. Engel, S. Aleksander, R. S. Nash, E. D. Wong, S. Weng, S. R. Miyasato, G. Sherlock, and J. M. Cherry, Saccharomyces genome database: Advances in genome annotation, expanded biochemical pathways, and other key enhancements, Genetics , iyae185 (2025).

[21] H. N. Lekkerkerker, R. Tuinier, and M. Vis, *Colloids and the depletion interaction* (Springer Nature, 2024).

[22] G. J. Fleer and R. Tuinier, Analytical phase diagrams for colloids and non-adsorbing polymer, Advances in Colloid and Interface Science **143**, 1 (2008).

[23] H. Lekkerkerker, Osmotic equilibrium treatment of the phase separation in colloidal dispersions containing non-adsorbing polymer molecules, Colloids and surfaces **51**, 419 (1990).

[24] H. N. W. Lekkerkerker, W. C.-K. Poon, P. N. Pusey, A. Stroobants, and P. B. Warren, Phase Behaviour of Colloid + Polymer Mixtures, Europhysics Letters **20**, 559 (1992).
[25] J. Wyman, Linked Functions and Reciprocal Effects in Hemoglobin: A Second Look, in *Advances in Protein Chemistry*, Vol. 19 (Elsevier, 1964) pp. 223–286.
[26] J. Wyman and S. J. Gill, *Binding and Linkage: Functional Chemistry of Biological Macromolecules* (Univ. Science Books, Mill Valley, Calif, 1990).
